# Site-specific decreases in DNA methylation in replicating cells following exposure to oxidative stress

**DOI:** 10.1101/2022.05.15.491578

**Authors:** Annika R. Seddon, Andrew B. Das, Mark B. Hampton, Aaron J. Stevens

## Abstract

Oxidative stress is a common feature of inflammation-driven cancers, and promotes genomic instability and aggressive tumour phenotypes. It is known that oxidative stress transiently modulates gene expression through the oxidation of transcription factors and associated regulatory proteins. Activated neutrophils produce hypochlorous acid and chloramines that can disrupt DNA methylation via methionine oxidation. The goal of the current study was to determine whether chloramine exposure results in sequence-specific modifications in DNA methylation that enable long-term alterations in transcriptional output. Proliferating Jurkat T-lymphoma cells were exposed to sublethal doses of glycine chloramine and differential methylation patterns were compared using Illumina EPIC 850K bead chip arrays. There was a substantial genome-wide decrease in methylation four hours after exposure that correlated with altered RNA expression for 24 and 48 hours, indicating sustained impacts on exposed cells. A large proportion of the most significant differentially methylated CpG sites were situated towards chromosomal ends, suggesting that these regions are most susceptible to inhibition of maintenance DNA methylation. This may contribute to epigenetic instability of chromosomal ends in rapidly dividing cells, with potential implications for the regulation of telomere length and cellular longevity.

## Introduction

The functional relationship between inflammation and cancer is widely accepted, however, the molecular and cellular mechanisms that contribute towards this relationship remain poorly defined. During pathogenic invasion, enhanced cell proliferation occurs in an environment of increased inflammation, and a state of chronic inflammation increases the risk of cancer (1). Proliferating cells that are exposed to an inflammatory microenvironment may develop genetic or epigenetic changes that are propagated in subsequent cell generations, even after inflammation subsides. DNA methylation patterns are often modified in human cancers resulting in the silencing of tumour suppressor genes and/or the activation of oncogenes (2). In an attempt to gain insight into how the environment may influence disease outcomes, many studies investigate how exposure to environmental factors can change patterns of DNA methylation. However, few studies have investigated the mechanisms behind the modification of DNA methylation and how these changes become established in the methylome. Many of the environmental factors associated with epigenetic patterning, including inflammation, are also recognised as risk factors in cancer (3).

Neutrophils are a rich source of oxidants and excessive or prolonged oxidant production can lead to tissue damage and chronic disease states (4). There is now considerable evidence to implicate neutrophils in all stages of neoplastic disease progression, from initiation through to malignancy (5). Neutrophil oxidants can directly damage DNA leading to mutation, they can activate key enzymes and transcription factors pivotal for tumour growth and repress anti-cancer T-lymphocytes (6–9). While oxidants have been shown to interfere with T-lymphocyte functional properties during inflammation, less is known about their potential to disrupt the metabolic pathways essential for maintaining epigenetic fidelity in these cells (10). One study has shown that histone methylation patterns in T-lymphocytes could be modified due to the depletion of methionine by tumour cells, however the impact on DNA methylation patterns were not investigated (11). Similarly, in an inflammatory environment, there is potential for neutrophil oxidants to oxidise methionine to methionine sulfoxide, restricting the availability of methionine not only to T-lymphocytes but also to tumour cells. Altered methionine levels can impact on DNA methylation machinery and conserved changes in methylation and subsequent gene expression could contribute to the transformed phenotypes observed in both these cell types as a consequence of neutrophil exposure.

Hypochlorous acid (HOCl), a major oxidant produced by the neutrophil enzyme myeloperoxidase (MPO), reacts with endogenous amines to yield chloramines (12). These species are cell permeable, longer lived and more selective in their reactivity than HOCl (13). Chloramines react readily with thiol groups and free methionine and cause cell damage and enzyme inactivation (14–16). We have previously observed that glycine chloramine (GlyCl) directly inhibits DNA methyltransferase (DNMT) activity, and oxidises methionine leading to depletion of the methyl donor, *S*-adenosyl methionine (SAM), resulting in a global decrease in 5-methylcytosine levels without significantly impacting cell viability (15). Through the use of Illumina850K EPIC arrays we have now discovered that oxidative stress associated with GlyCl exposure causes site-specific alterations in DNA methylation and long-term gene expression, and that the affected regions are enriched towards chromosomal ends.

## Results

We investigated DNA methylation and gene expression changes in cultured Jurkat T-lymphoma cells following exposure to glycine chloramine (GlyCl). Since glycine chloramine is quickly consumed by cell and media thiols and methionine, and it acts to impair maintenance methylation during new DNA synthesis, cells were synchronized with a thymidine block to maximize the number undergoing DNA synthesis at the time of oxidant exposure. Thymidine halts cell division at the G_0_/G_1_ phase of the cell cycle, and we have previously demonstrated that a large proportion of cells enter S-phase two hours after release from thymidine block (17). We therefore exposed cells to GlyCl immediately after the thymidine was removed.

### Glycine chloramine sensitivity and cell proliferation

Jurkat cells were treated with either a single bolus of 200 µM GlyCl (treatment) or left untreated (control). Exposure to GlyCl corresponded with a significant decrease in both percentage growth at 24 and 48 h (*p* = 0.02 and 0.01) (Figure 1a), and a small decrease in cellular viability at 24, 48 and 72 h (*p* = 0.02, *p* = 0.01 and *p* = 0.03 respectively) (Figure 1b). The synchronization process itself results in a ∼20% loss in cell viability.

**Figure 1:**
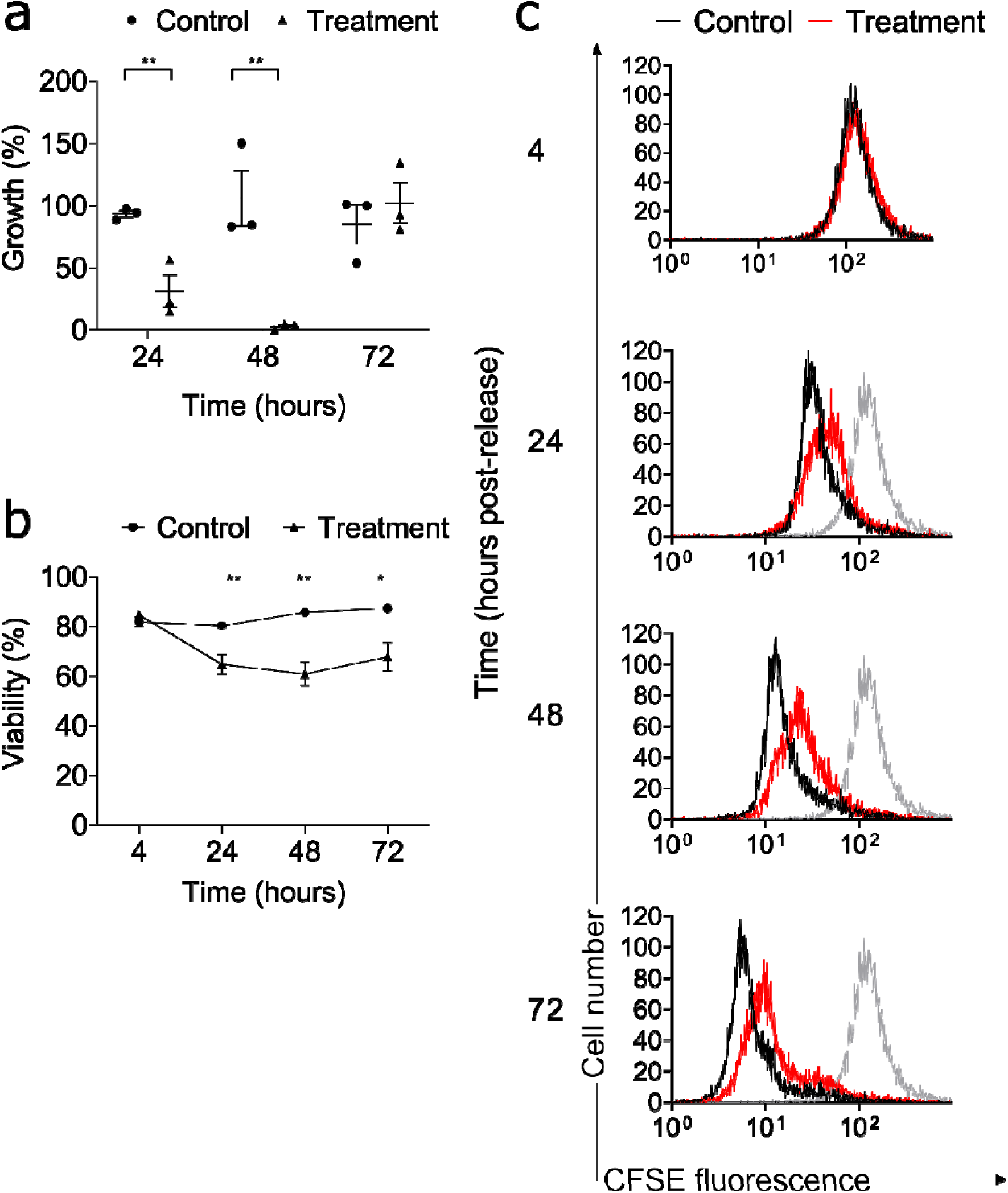
Viability, growth and CFSE dilution of GlyCl treated cells at each major time point. Cell measures were conducted at 24, 48 and 72 hours post-release. Circles represent control samples and triangles represent treatment samples. Data are means and SE of 3 independent experiments. Significant differences were determined with paired t-tests and are denoted with asterisks * = *p* <0.05, ** = *p* <0.01. **a,** Cell growth after treatment **b,** Cellular viability after treatment **c,** CFSE dilution of GlyCl treated cells at 24, 48 and 72 hours post-release. The number of cells is displayed on the y-axis and the concentration of CFSE fluorescence is on the x-axis. Representative cell histograms are shown for control (black) and treated (red) cells. The grey histograms represent the control cells from the 4 h time point for reference.

Cell proliferation was monitored with the cell permeable fluorescent dye CFSE over 72 hours for the control and treatment samples, with fluorescence intensity halving at each cell division. GlyCl treatment delayed cellular proliferation compared to control 24 h post treatment (Figure 1c), but subsequently they appeared to proliferate at a normal rate, with the treated cells at 72 h having the same concentration of CFSE as the control cells at 48 h (Figure 1c). These observations are consistent with the percentage growth rate (Figure 1a), and we conclude that after oxidant treatment a large number of viable cells that had undergone at least one round of cell division remained.

### Glycine chloramine alters DNA methylation

Principal component and hierarchical clustering analysis demonstrated that the 4 h treatment replicates clustered separately from all the other groups. (Figure 2a & b). This indicates that treatment corresponded with the most significant source of variation, and that consistent changes were observed within these samples.

**Figure 2:**
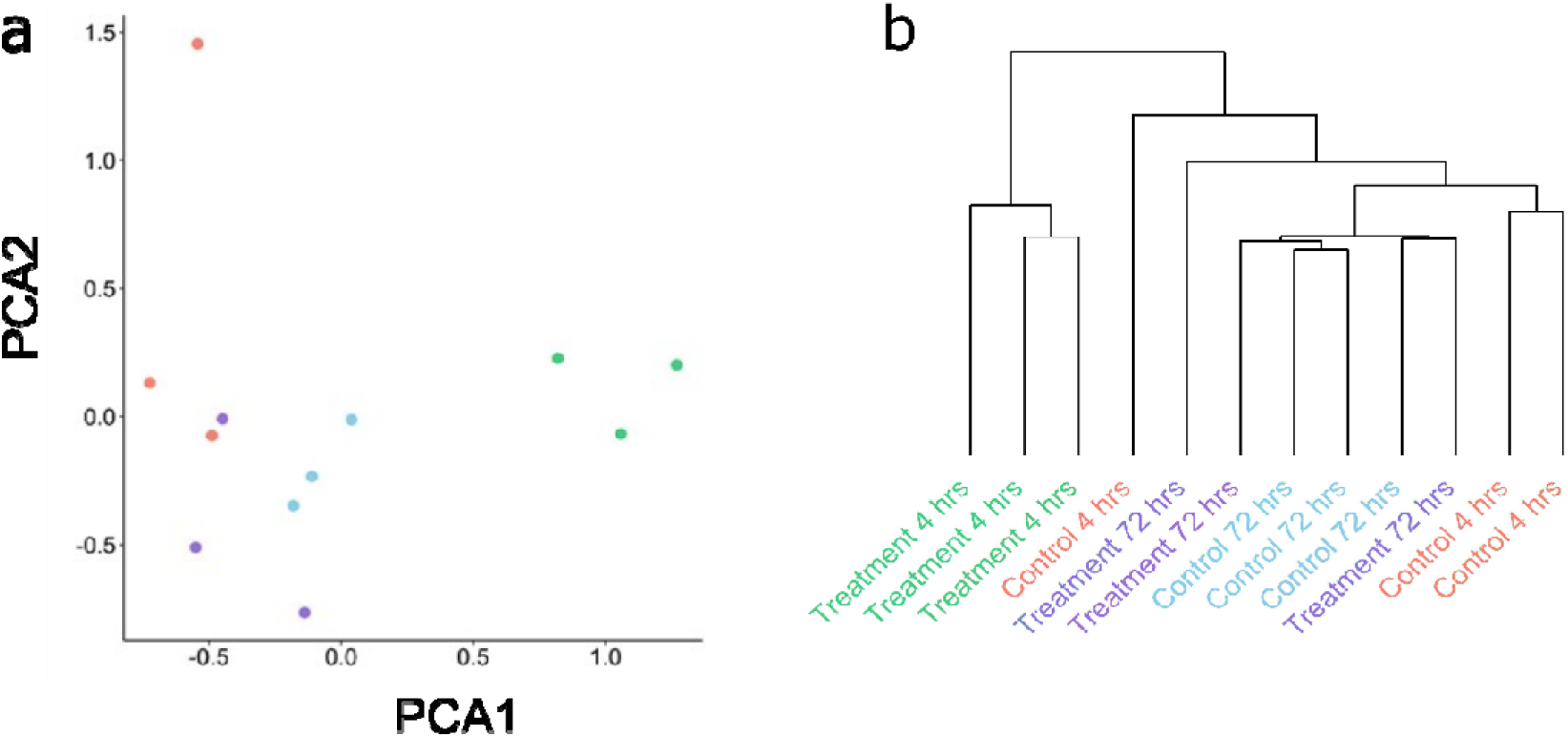
Unsupervised assessment of data variability. **a,** Multidimensional scaling of M-values, with the distances for leading log_2_FC in dimension 1 represented on the x-axis and the leading log_2_FC in dimension 2 are represented on the y-axis. Red and blue dots represent control samples (4h and 72h time points respectively), green and purple dots represent GlyCl treated samples (4h and 72h time points, respectively).**b,** Hierarchical clustering of β-values for all probes. The relative change in β-values is represented on the y-axis and individual samples are represented on the x-axis.

At 4 h, the largest and most significant fold changes were decreases in methylation in the treatment group, compared to the control (Figure 3a). Both the effect size and significance was considerably reduced at 72 h, with very few probes achieving genome-wide significance (Figure 3b). These results indicated that the majority of significant changes in DNA methylation due to GlyCl treatment observed at 4 h had reverted by 72 h (Figure 3a & b).

**Figure 3:**
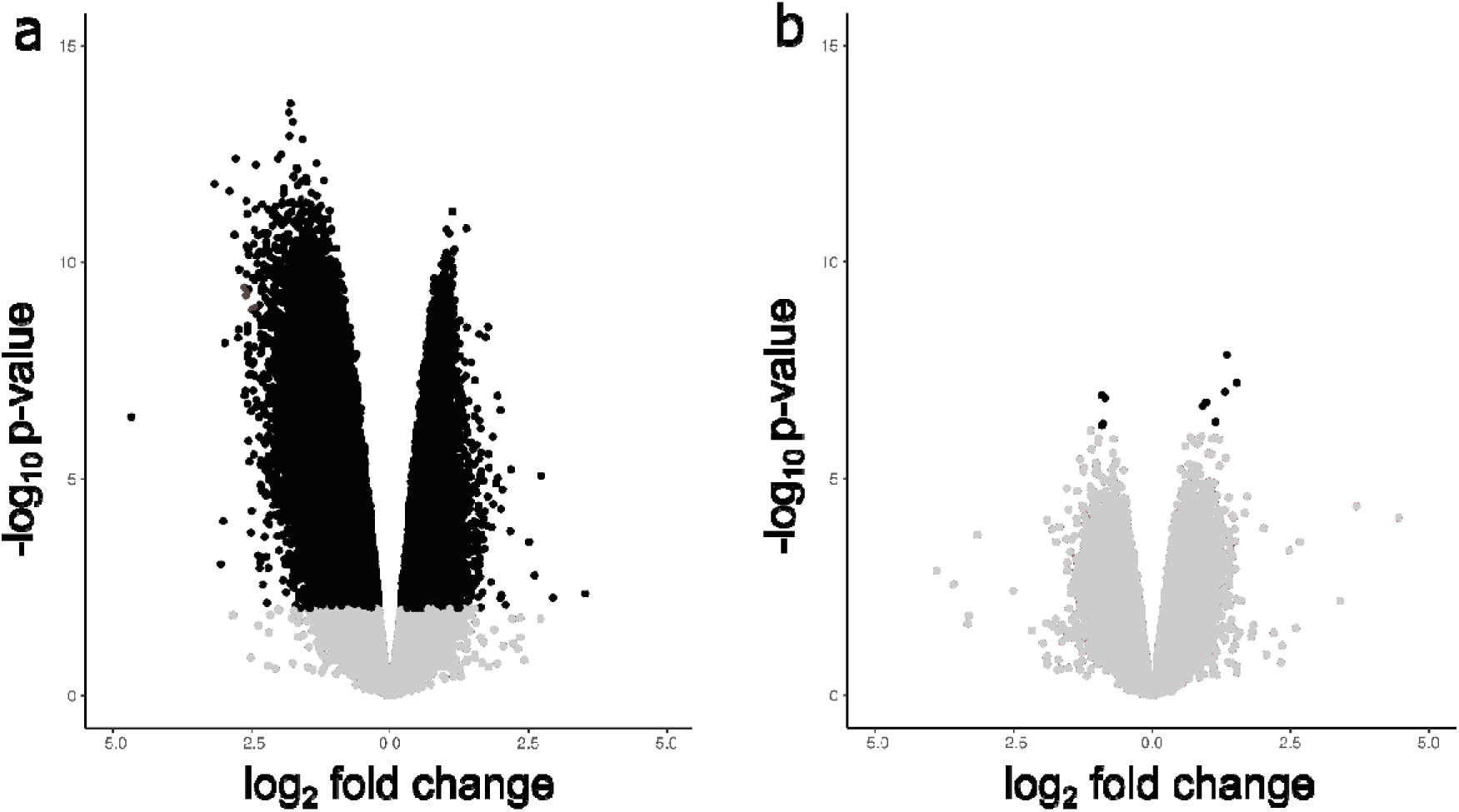
Volcano plots. Volcano plot displaying log_2_ fold changes (M-values) for all probes on the x-axis versus statistical significance on the y-axis -log_10_ p-value. Black dots represent probes with an adj. *p* <0.05. **a,** GlyCl exposure at 4 h **b,** GlyCl exposure at 72 h.

Our previous mechanistic studies indicated that chloramines affect methylation through depletion of SAM over two hours (15). To determine if GlyCl can impact methylation by an alternate mechanism, thymidine blocked Jurkat cells were exposed to 200 µM GlyCl at 2 hours post-release and methylation patterns assessed 2 hours later. Under these conditions the cells would have sufficient SAM to undertake the necessary maintenance methylation, but it provides an opportunity for the oxidant to impact DNA methylation through an alternate mechanism (17). There was no significant difference in percentage growth or viability observed in the treatment cells at 24 or 72 h however, a significant decrease in percentage growth was observed at 48 h *(p* = 0.03) (Supplementary Figure 1a & b). There were no distinctive groupings observed in principal component analysis and hierarchical clustering analysis revealed a lack of clustering in the treatment or control samples and groupings appeared random between replicates (Supplementary Figure 2a & b). Probe-wise comparisons of significance versus effect size change confirmed that GlyCl treatment at 2 hours post-release from thymidine block did not correspond with significant changes in DNA methylation at either 4 or 72 hours (Supplementary Figure 3a & b and Supplementary Tables 1 & 2). No further analyses were undertaken with this dataset.

### Glycine chloramine causes site-specific changes in DNA methylation

Epigenetic variability across a population of cells can result in transcriptional heterogeneity and promote diverse functional states (18, 19). We therefore investigated the differential variability of all probes (DVP) after GlyCl exposure. At 4 h, the treatment group demonstrated an increase in differential variability at 5,910 positions compared to control, and a decrease at 13,769 positions. At 72 h, 1,892 positions demonstrated an increase and 2,743 demonstrated a decrease. Twenty-three DVPs were observed at both 4 and 72 h, however only six of these DVPs (26%) demonstrated a consistent direction of change between the two time points (Supplementary Table 3).

We next identified site-specific DNA methylation changes that were associated with GlyCl exposure, by using a linear regression model that accounted for time and biological replicate. At 4 h, using a log_2_FC cut-off of 1, there were 92,665 CpG sites that displayed a significant decrease in methylation and 66,403 that demonstrated an increase. These DNA methylation changes were mapped by genomic location, and it appeared that many significant changes were concentrated at specific gene loci, and towards the chromosomal ends (Figure 4 and Supplementary Figure 4). In order to select the DNA methylation changes that were most likely to be of biological interest, the log_2_FC was increased to 2. This returned 124 CpG sites that demonstrated decreased methylation and no CpG sites that demonstrated an increase, which indicates that the largest changes in differential methylation were all decreases. We then further filtered to only include probes that demonstrated a minimum of a 20% mean change between treatment and control (Table 1 and Supplementary Figure 5).

**Figure 4:**
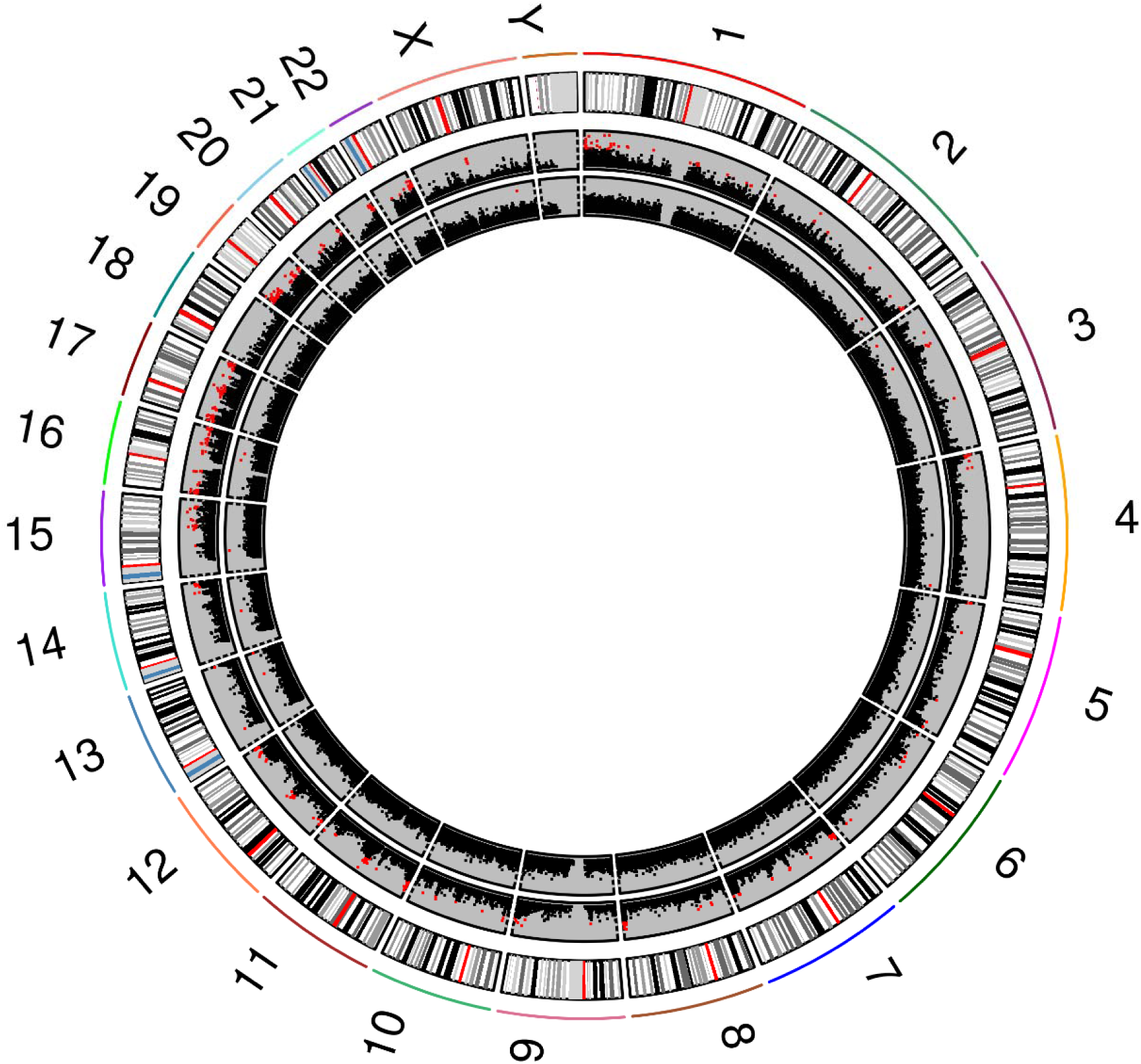
Genome-wide circular plot of all CpG sites. All CpG sites are ordered per chromosome position, and *p*-values as the log_10_ (*p* values) are presented on the y-axis. Positions that reached genome-wide significance as determined using the “Benjamini, Hochberg” method within Limma are represented by the red squares. The 4-hour time point is represented in the outer ring and the 72-hour time point is represented on the inner ring. The bands in the outermost ring show the position of the centromere (red) and the states of chromatin packing (black representing the most densely packed chromatin through to light grey representing more loosely packed chromatin).

**Table 1:**
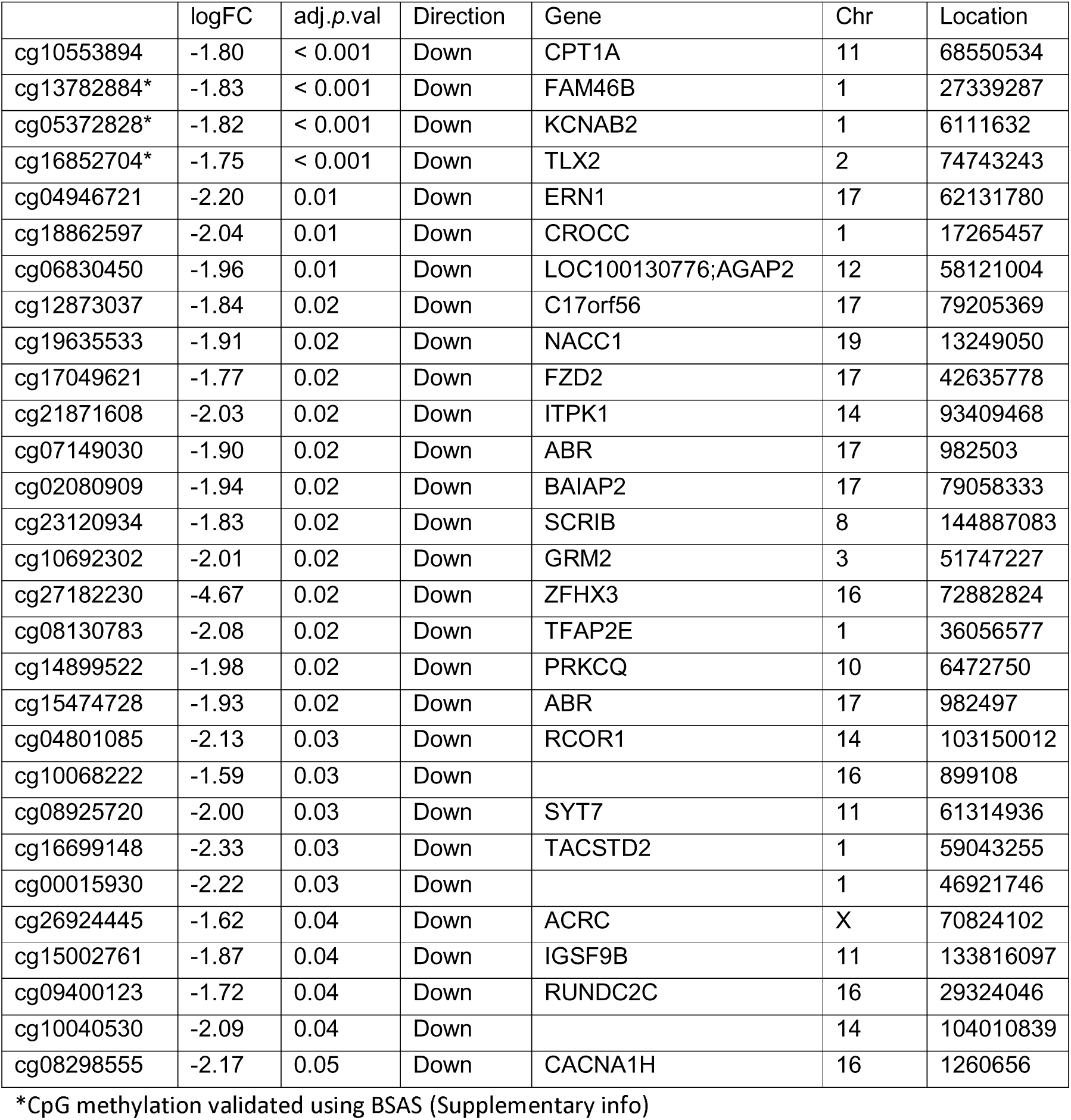
Top most significant differentially methylated probes with a Log_2_FC ∼2 and a >20% mean change between treatment and control at 4 h.

At 72 h, six CpG sites demonstrated a significant increase in differential methylation and four CpG sites demonstrated a significant decrease (adj. *p* <0.05), when assessed using a log_2_FC cut-off of 1 (Supplementary Table 4). Two of these differentially methylated CpG sites were also significant at 4 h. These CpG sites corresponded to the *ERC2* and the *MLF1IP* genes, and had a smaller log_2_FC at 4 h than at 72 h. Furthermore, neither exhibited a consistent direction of change (Supplementary Figure 6a & b).

To identify differentially methylated gene regions (DMRs), we limited analysis to investigate loci containing five or more consecutive probes that displayed a significant average change in differential methylation greater than 10%. Eighty-five significant DMRs were observed at 4 h, and 14 of these consisted of 10 or more adjacent significant CpGs (Supplementary Table 5). The top two DMRs with the highest number of significant adjacent CpGs mapped to the *MAPK8IP3* gene region (18 CpGs) (Figure 5a) and the *TFAP2E* gene region (16 CpGs) (Figure 5b). No significant DMRs were observed at 72 h under these parameters. When the parameters were relaxed to detect changes in methylation greater than 5%, there were four significant DMRs (Supplementary Table 6) however, the direction of methylation changes were opposing between the 4 and 72 hour time points.

**Figure 5:**
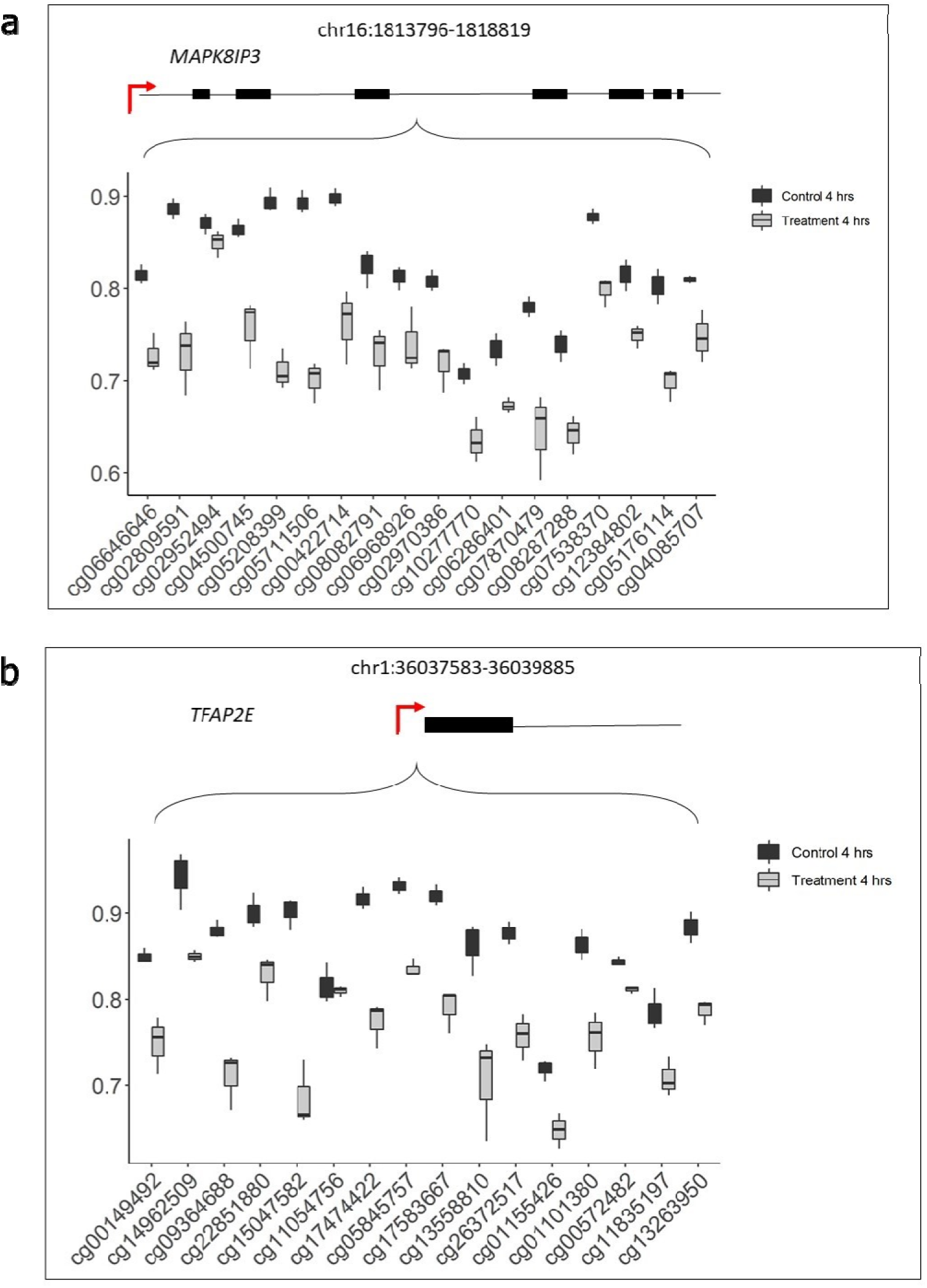
Differentially methylated gene regions. **a**, DMR corresponding to the MAPK8IP3 gene (chromosome 16:1813796-1818819, width:5024 bp) that displayed a significant change in methylation (β-values (y-axis)) across 18 CpGs (x-axis) between the treatment (grey) and control (black) for the 4 hour time point. **b,** DMR corresponding to TFAP2E gene (chromosome 1:36037583-36039885, width:2303 bp) that displayed a significant change in methylation(β-values (y-axis)) across 16 CpGs (x-axis) between the treatment (grey) and control (black) for the 4 hour time point. Gene structure is placed on top of each graph, exons are shown as black bars, and the transcriptional start site is marked by a red arrow. The CpGs cg15047582 (TFAP2E) and cg05208399 (MAPK) were validated using BSAS (Supplementary info)

The differentially methylated CpGs at 4 h appeared to be concentrated towards the chromosomal ends (Figure 4, Supplementary Figure 4). To investigate this observation, the data was categorized in regions representing 10% bins for each chromosome. There was a considerable enrichment in the number of significant probes (49%) that were located within 10% of the end of the chromosomes, even after correction for probe bias (Figure 6a). This observation also corresponded with a substantial decrease in the log_2_FC of significant probes towards the chromosomal ends (Figure 6b). Specifically, these regions correspond with between 2-5 million base pairs (bp) from each chromosome end, and are towards the outer limit of CpG sites that are included in the array. The array does not include telomeric or the subtelomeric regions, which are typically characterised as within 50,000 bp upstream from the telomeric regions.

**Figure 6:**
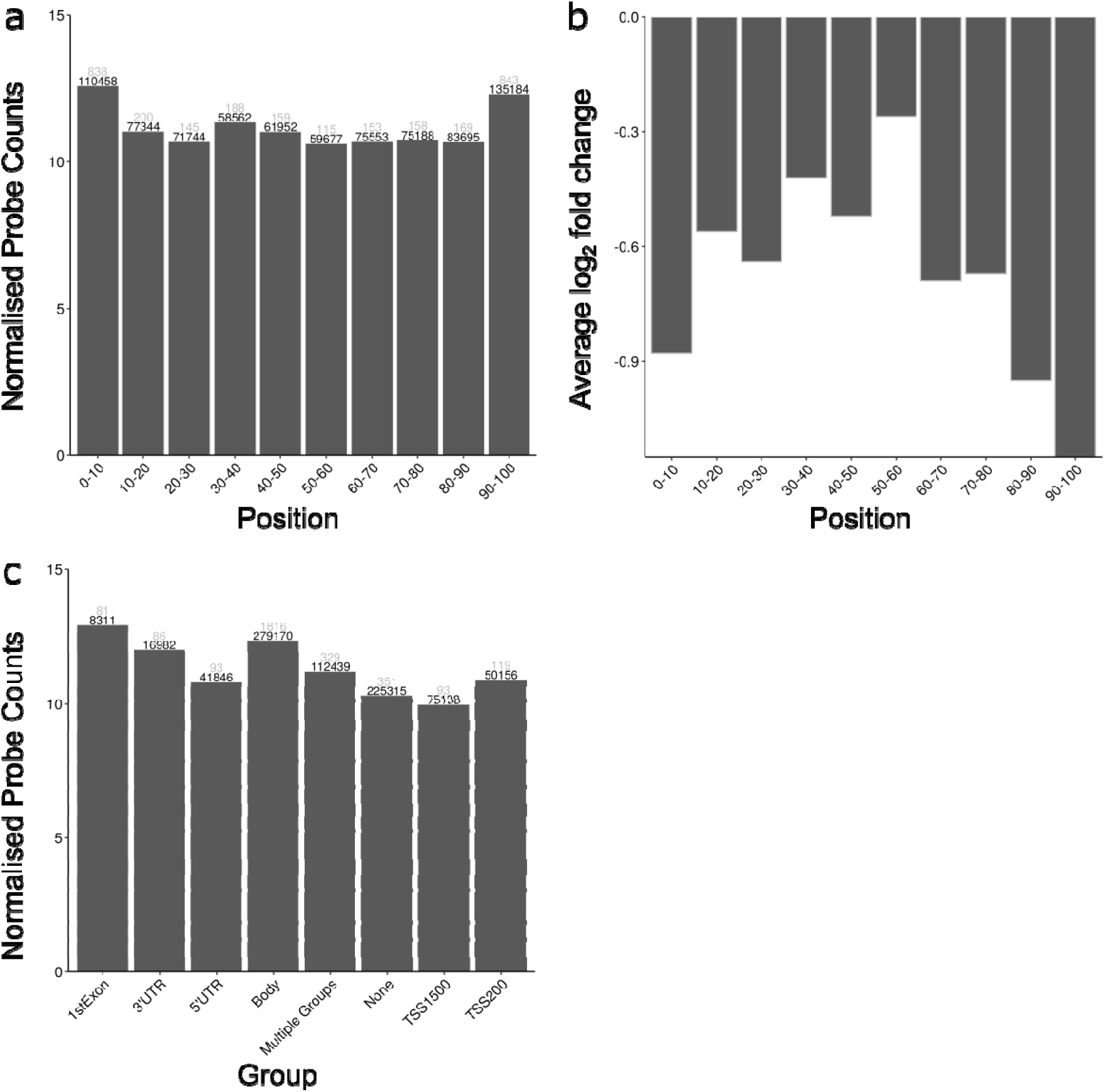
Significant differentially methylated CpGs (log_2_FC >1.5) plotted by genomic features. **a,** The normalised number of significant differentially methylated CpGs plotted by percent of total chromosome length. The x-axis shows the distance from the start of all chromosomes as a percentage. The y-axis shows the normalised number of CpGs that showed significant change. Simple Scaling Normalization (SSN) was adapted from the LUMI pipeline to control for probe bias. Black numbers represent the total number of probes that bind in each region, and grey numbers represent the number of significant differentially methylated probes that bind in that region. **b,** Average log_2_ Fold Change for the significant probes (as determined by Limma) binned into 10 percent intervals for all chromosomes**. c,** Significant differentially methylated CpGs plotted by gene element which included: Gene body, 5’untranslated regions (UTR), 3’UTR, first exon, area from the transcriptional start site (TSS) to − 200 nucleotides upstream of TSS, and the region between 200 and 1500 nucleotides upstream of TSS, multiple groups encompasses CpGs that are not unique to a single group. The y-axis shows the normalised (SSN) number of CpGs that showed significant change. Black numbers represent the total number of probes that bind in each region, and grey numbers represent the number of significant differentially methylated probes that bind in that region.

The majority of significant differentially methylated probes (64%) occurred within gene bodies or the first exon, with a substantial under-representation from non-transcribed gene elements (Figure 6c). There was no substantial enrichment in significant probes that occurred in relation to CpG islands (data not shown).

### Gene expression change in response to GlyCl

We next used RNA sequencing to investigate the impact of GlyCl exposure on differential gene expression in the same Jurkat samples used in the DNA methylation analysis. The statistical approach was consistent with that used to detect significant differential methylation, where changes were contrasted between the groups (treatment versus control), and the effect of biological replicate was accounted for. In order to minimise the number of significant differentially expressed genes, while still selecting for results of highest biological significance, thresholds containing an absolute log_2_FC > 1.15 and an adjusted *p* value < 0.05 were incorporated into the statistical model. Gene expression changes were assessed at 4, 24 and 48 h, low RNA yield meant that there was insufficient sample available for an accurate assessment at 72 h.

Hierarchical sample clustering and principal component analysis indicated that 4 and 24 h treatment samples were the most distinct from all other groups (Supplementary Figure 7a &b).

At 4 h, there were 277 genes up regulated in the treatment group relative to the control group, and 203 genes down regulated, and these exhibited a similar significance and magnitude of change (Supplementary Table 7). Genes that demonstrated decreased expression were associated with 27 GO terms and two KEGG terms, after correction for multiple testing (Supplementary Table 8). However, genes demonstrating increased expression were not associated with any GO or KEGG terms.

At 24 h there was the largest number of significant differentially expressed genes observed over the duration of the experiment, with 2155 genes up regulated in the treatment group relative to the control group, and 1902 genes down regulated. The largest fold changes all corresponded with increased expression, and the genes that showed decreased expression generally corresponded with smaller *p-*values and had a lower magnitude of change (Supplementary Table 9).At 48 h, the magnitude of changes and number of significant changes substantially reduced, with 121 genes up regulated in the treatment group relative to the control group, and 61 genes down regulated (Supplementary Table 10, Supplementary Figure 8).

To investigate the relationship between DNA methylation changes and gene expression after GlyCl treatment, the significant DNA methylation changes (Log_2_FC >1.5) observed at 4 h were correlated against significant changes in differential gene expression (Log_2_FC >1.5) at the subsequent time points. We limited our analysis to only assess significant DNA methylation sites that were located within the promoter regions of genes that displayed significant differential expression.

In total, there were 47,133 probes that displayed significant DNA methylation, and were located within gene promoter regions. Of these, there were 560 differentially methylated CpGs located within the promoter regions of 266 differentially expressed genes at 4 h. A significant, negative correlation (adj. *p <* 0.05) was observed between 69 CpGs and 48 unique genes (Supplementary Table 11).

There were 4751 differentially methylated CpGs from 4 h located within the promoter regions of 2255 differentially expressed genes at 24 h. A significant negative correlation (adj. *p <* 0.05) was observed between 398 CpGs and gene expression in 400 genes (Supplementary Table 12). No significant correlations were observed between methylation levels at 4 h and gene expression at 48 h. A summary of correlations between promoter DNA methylation and gene expression are presented in Table 2.

**Table 2:** Summary of correlations between promoter DNA methylation at 4 h and gene expression at the indicated time point.

| Time point | Total number of assessed probes * | Total number of corresponding genes** | Number of genes showing significant correlation *** | Percent of genes showing positive correlation | Percent of genes showing negative correlation |
| --- | --- | --- | --- | --- | --- |
| 4 h | 560 | 266 | 83 | 41.5 | 58.9 |
| 24 h | 4751 | 2255 | 787 | 63.9 | 37.7 |
| 48 h | 251 | 106 | 0 | 0 | 0 |

\* Total number of probes that showed significant DNA methylation change, and were located within the promoter region of genes that showed a significant change in gene expression
\*\* Total number of genes corresponding with the assessed probes
\*\*\* Total number of genes showing a significant pairwise sample correlation between DNA methylation values and gene expression values (see methods).

There was no apparent bias of significantly correlated genes towards specific genomic locations at 4 h (Figure 7). However, when the 4 h DNA methylation was correlated with the 24 h gene expression levels, significant correlations were frequently clustered together along the chromosome at adjacent gene regions, for example, the cluster of *HIST* genes on 6p22.2. (Figure 8).

**Figure 7:**
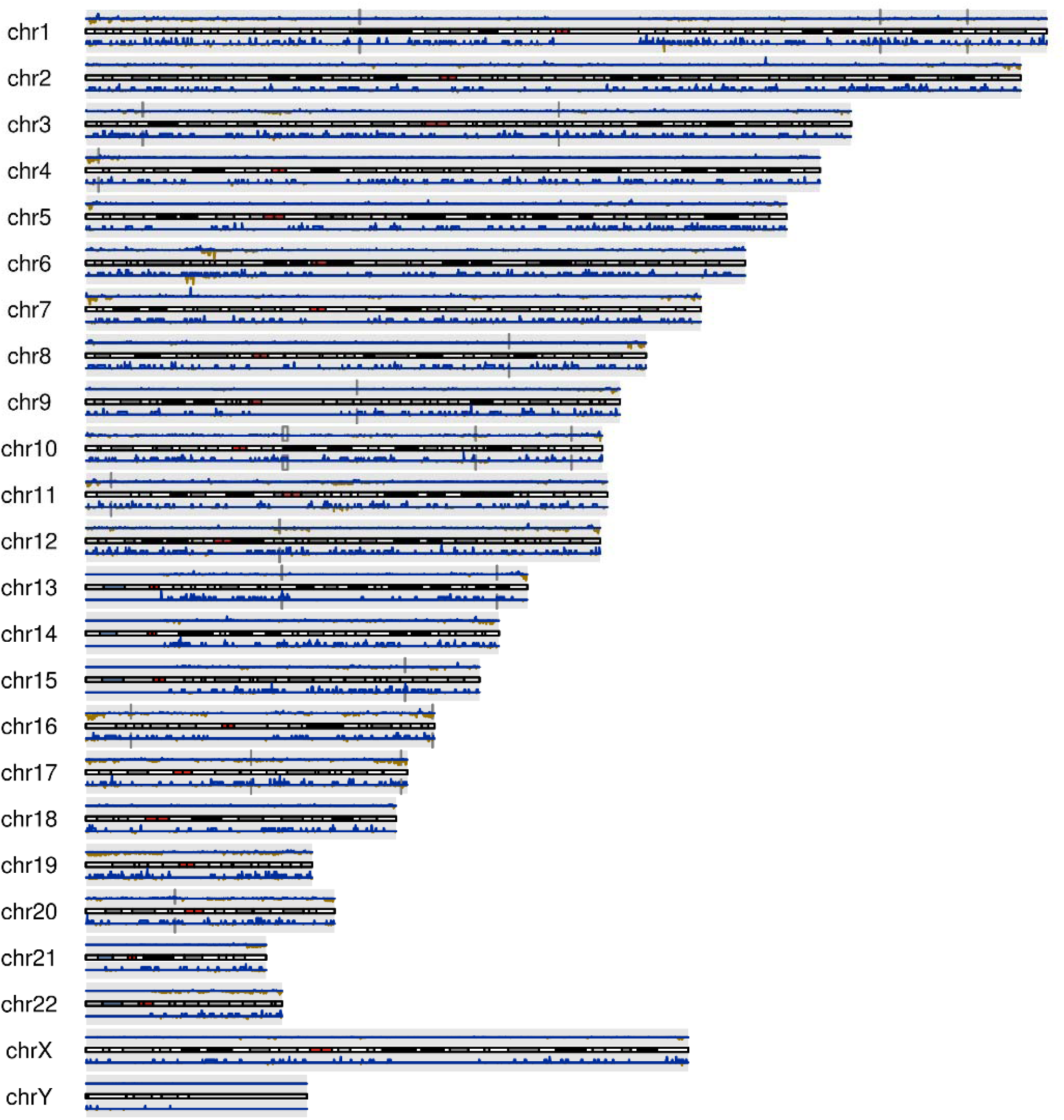
Genome-wide alignment of significant methylation logFC and significant gene expression logFC at 4h. Blue peaks represent positive logFC (treatment vs control) and yellow peaks represent negative logFC (treatment vs control). Data is presented as density of logFC over one million base pair windows. Dark Grey rectangles represent significant correlation between promoter methylation and gene expression changes. The horizontal bars show the position of the centromere (red) and the states of chromatin packing (black lines representing the most densely packed chromatin through to light grey representing more loosely packed chromatin).

**Figure 8:**
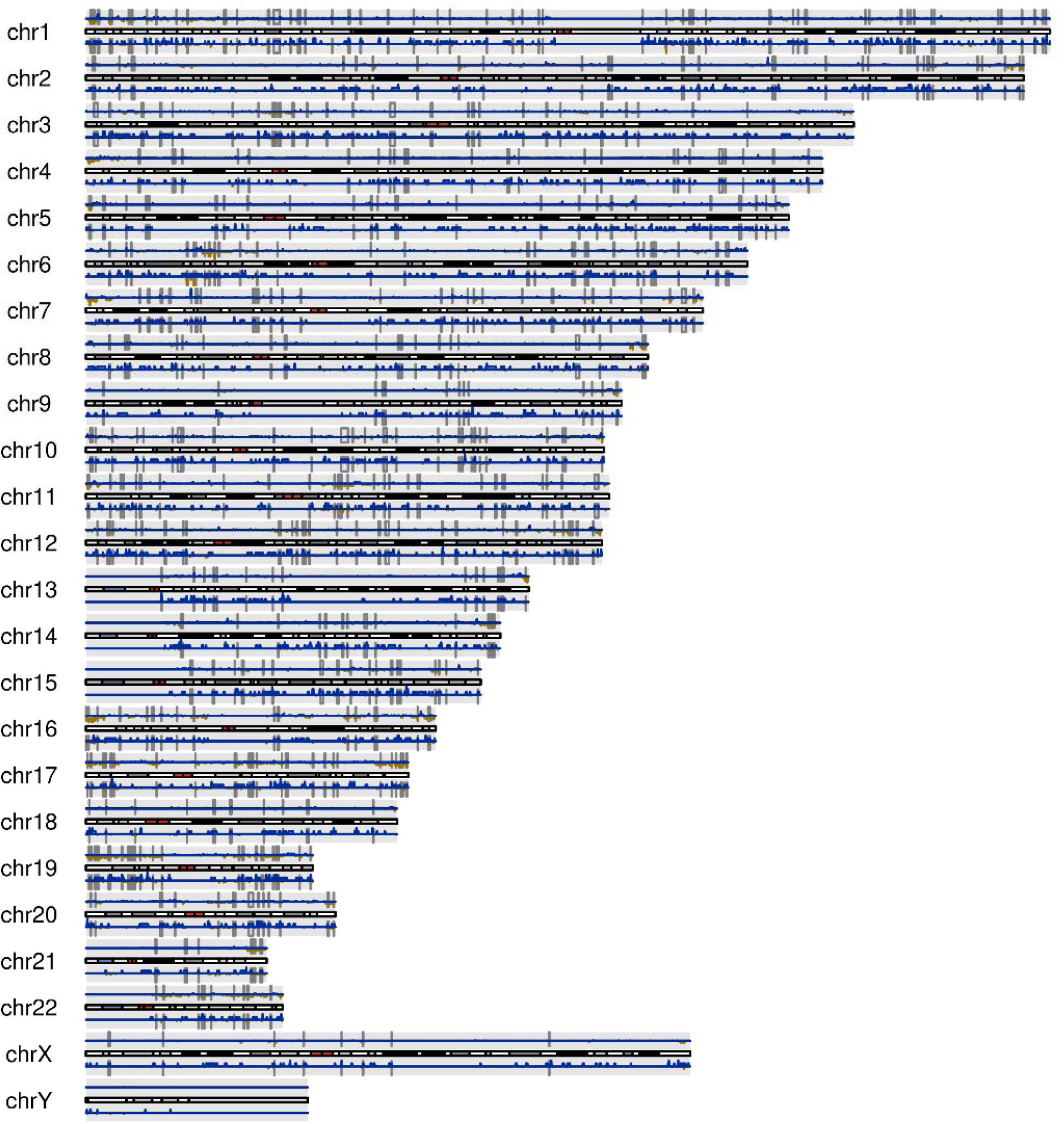
Genome-wide alignment of significant methylation logFC at 4 h and significant gene expression logFC at 24 h. Blue peaks represent positive logFC (treatment vs control) and yellow peaks represent negative logFC (treatment vs control). Data is presented as density of logFC over one million base pair windows and dark grey rectangles represent significant correlation between promoter methylation and gene expression changes. The horizontal bars show the position of the centromere (red) and the states of chromatin packing (black lines representing the most densely packed chromatin through to light grey representing more loosely packed chromatin).

### Pathway analysis of DNA methylation regulated gene expression changes

Pathway analysis was conducted with the top most significant CpG sites that demonstrated decreased methylation at 4 h within the KEGG and GO databases. Fifty-two significant KEGG pathways were identified that had a FDR <0.05 and 13 significant GO pathways were identified for this dataset (Supplementary Table 13).

To investigate whether genes that demonstrated a significant correlation between promoter methylation and gene expression were enriched for similar biological function, these genes were also assessed using GO and KEGG databases. There were 10 significant GO pathways associated with genes that demonstrated a significant negative correlation between samples at 4 h and 53 genes that demonstrated a significant negative correlation between DNA methylation at 4 h and gene expression at 24 h (Supplementary Table 14). Numerous pathways from both time points represented DNA break repair mechanisms. Interestingly, several of the top most significant pathways are of direct relevance in the development and progression of Alzheimer’s disease, including microtubule based processes (20), extracellular vesicles (21), 3’,5’-cyclic-nucleotide phosphodiesterase activity (22), cytokine stimulus (23), and negative regulation of cell-cell adhesion (24). Other significant pathways were involved in the development of various cancers, and in acute inflammatory response pathways.

To further investigate what biological disease outcomes might be influenced by the epigenetic regulation of gene expression following GlyCl treatment, we assessed the significantly correlated genes using WebGestalt (25) against the OMIM database. This analysis confirmed that the regulation of several key Alzheimer’s and cancer related genes might be linked with the corresponding methylation levels. There were also numerous important genes involved in non-insulin-dependent (type II) diabetes mellitus, a disease linked with inflammation (26) (Supplementary Table 15).

## Discussion

Epigenetic modification of DNA is a means by which environmental stimuli can modulate human gene activity, but the exact mechanisms behind these processes remain unclear. We investigated whether cellular proliferation in an inflammatory environment can trigger epigenetic changes that lead to long-term changes in gene expression, including propagation in subsequent cell generations. By using methylation arrays, we have shown that glycine chloramine exposure results in large genome-wide decreases in methylation as well as increased heterogeneity of methylation patterns in the cell population after completion of DNA replication. This was associated with significant changes in gene expression 24 and 48 h after oxidant exposure, and while most of the significant methylation changes observed were corrected during subsequent rounds of cell division, some changes were conserved. The failure to restore correct methylation patterns could lead to the propagation of these aberrant patterns and subsequent alteration in cell function. Furthermore, the chromosomal position of the most significant differentially methylated CpG sites were concentrated in gene bodies within 2-5 million base pairs of the chromosomal ends, indicating that sequence-specific alterations in methylation can occur through interference in a pathway that acts on molecular methylation machinery.

At sites of infection and inflammation the heme enzyme myeloperoxidase generates HOCl, and its primary targets are most likely to be extracellular methionine and amines (27, 28). Resultant chloramines, such as the GyCl used in this study, will therefore be prominent under physiological conditions. Their lower reactivity provides them the opportunity to pass into neighbouring cells and alter cell function (14, 27). It is well known that oxidants can impact gene expression through direct oxidation of transcription factors and associated regulatory proteins (29, 30). HOCl exposure activates several transcription factors in bacterial systems, resulting in the upregulation of genes that protect the organism from oxidative stress (31–34). Likewise in mammalian cells, HOCl-induced modified gene expression has been observed through its action on stress-sensing transcription factors such as Nrf2 (NF-E2-related factor 2) and NF-κB (nuclear factor kappa-light-chain-enhancer of activated B cells), both through the oxidation of the regulatory proteins Keap-1 and IκB (35–39). These changes are typically transient, with thiol reduction occurring quickly after removal of the oxidant. The epigenetic changes we have observed in our study provide an alternate mechanism of redox regulation of gene expression and one that will last substantially longer than that achieved through direct effects of transcription factor activity.

Glycine chloramine exposure prior to the cells entering into the stage of DNA replication had a complex effect on 5-methylcytosine in synchronised Jurkat cells. Large genome-wide changes in differential methylation and differential variability were observed at 4 h, and abnormal methylation was still detected at 72 h, after all cells had undergone at least one round of cell division. We have previously shown that GlyCl treatment depletes methionine and SAM levels substantially two hours after treatment, and anticipated that exposure would be required prior to cellular DNA replication (15). This was confirmed by our delayed GlyCl exposure experiment, which demonstrated no significant changes in DNA methylation. This exposure occurred at a time at which DNMT would be most active and before SAM levels could become significantly depleted by treatment. This suggests that the effect of GlyCl on SAM levels is the critical driver of the methylation changes observed rather than direct inhibition on DNMT activity.

The top most significant CpGs all demonstrated a decrease in differential DNA methylation, with a substantial number of genes involved in proliferation, tumour progression and cell death (40–47). Pathway analysis reinforced this observation where enriched pathways included vascular endothelial growth factor (VEGF) signalling (86%), mitogen-activated protein kinases (MAPKs) (71%) and apoptosis (67%). Two CpG sites corresponding to the *ERC2* and the *MLF1IP* genes demonstrated a significant change at both 4 and 72 h, but were not consistent in the direction of change, and also had a smaller log_2_FC at 4 h than at 72 h. This may suggest that the most commonly occurring methylation changes (those with the largest log_2_FC) are either detrimental to cell division and are corrected by the cell, or they trigger an apoptotic pathway and are not represented at 72 h. However, changes that were more highly represented (larger log_2_FC) at 72 h than at 4 h, may be indicative of cells that have acquired a reproductive advantage.

Many of the changes in methylation observed at 4 h were consistent across relatively large numbers of adjacent CpGs mapped to the same gene region. There were 18 adjacent CpGs that demonstrated a significant decrease in methylation mapping to the *MAPK8IP3* gene. The *MAPK8IP3* gene encodes the MAPK 8 interacting protein 3 also known as JNK/stress-activated protein kinase-associated protein 1 (JSAP1). MAPK8IP3 is a scaffolding protein for the mitogen-activated protein kinase (MAPK)/c-Jun N-terminal kinases (JNK). MAPKs are involved in a broad range of signalling pathways including proliferation, differentiation, autophagy and apoptosis which are pivotal for tumour invasiveness and progression (48). Oxidative stress has been shown to modify MAPK pathways in other cell types and can either promote a protective effect against oxidation (49, 50), or trigger apoptosis (51). It should be noted that because Jurkat cells are an acute T-cell leukemia cell line they may be prone to developing changes in cancer related pathways. Jurkat cells have also undergone substantial genomic rearrangement, and may be more tolerant of changes in DNA methylation than a primary cell (52).

Taken together, our results show that a single exposure of GlyCl at the onset of replication had a significant initial impact on DNA methylation at important genes that can either confer a survival advantage or induce cell death. These enriched pathways were not observed at the 72 h, indicating that the replenishment of methionine/SAM levels allowed for restoration of methylation by active DNMTs.

To investigate if the DNA methylation changes observed in response to GlyCl treatment could regulate gene expression, we sequenced RNA transcripts from matching cell aliquots. The results demonstrated substantial changes in gene expression corresponding with GlyCl, which has not been previously reported.

There was a strong negative correlation observed between probes demonstrating a significant change in promoter DNA methylation at 4h and gene expression at both 4 and 24 h. These genes were highly concentrated within regions of open chromatin, which is where the majority of DNA methylation changes were observed. Unfaithful methylation in gene-rich areas may have important implications for replication origin selection, as epigenetic markers have been shown to favour open chromatin structures (53, 54).

Pathway analysis indicated that some of the top most significant genes contribute towards aspects of cell division. Changes in these genes should be interpreted with caution since GlyCl appeared to stall cellular replication, and the treatment and control samples were no longer synchronised. At 48 h, both treatment and control cells had settled into an equivalent asynchronous state, but significant changes were still detectable, indicating that cell cycle differences would not be the sole factor in altered gene expression. GlyCl is also likely to have a detrimental effect on genetic stability, as indicated by the significant up regulation of gene pathways involved in DNA repair. Although this study was carried out using only one cell type, the mechanism of inhibition of methylation should be the same in all cells.

One of the most interesting findings of our study was the observation that significant methylation changes were highly concentrated towards the chromosomal ends. The precise mechanisms by which GlyCl targets methylation in these regions is unclear, but we did not observe this effect for hydrogen peroxide (17). We were limited by the scope of the bead chip array, which does not cover telomeric and subtelomeric regions, but this provides an interesting avenue for further research. Regions close in proximity to the telomeres have been shown to regulate telomere length, impacting cell longevity and function (55). It is worth noting that the TERT gene, responsible for maintaining telomere length during replication, demonstrated a significant decrease in gene expression 24 h after treatment. This effect demonstrated a positive correlation with promoter methylation, and is consistent with previous observations (56). Telomeres contain unmethylated repetitive sequences found at the terminal ends of each chromosome that preserve the integrity of the chromosome by protecting against chromosomal degradation, redundant DNA repair and recombination or fusion events (57). While telomere shortening is considered a by-product of somatic cell division, altered telomere length is often observed in ageing and age-related pathologies, particularly chronic inflammation and cancer (58–64). Increased telomere length is a common characteristic of advanced cancer types that often have reactivated telomerases that repair and lengthen end caps allowing for cellular immortality (65). Numerous studies have reported associations between epigenetic changes in CpG rich subtelomeric regions and the regulation of telomere length (66–70). Methylation of the *TERT* promoter plays an integral role in activation of telomere maintenance machinery in the context of human cancer, and has important clinical and biological implications (71). This CpG was located in a CpG rich region upstream of the *TERT* promotor, called *TERT* Hypermethylated Oncological Region (THOR) (56). Methylation at this site has been demonstrated to direct TERT expression, and has a unique cancer specific signature (56).

Overall, this study indicates that immune cell-derived oxidants can cause sequence-specific alterations in DNA methylation with resulting long-term changes in gene expression. These experiments were performed by exposing cells to a single oxidant bolus. In a physiological setting, cells would likely be subjected to repeated or continuous oxidant exposure, and at sub-lethal doses, this is even more likely to result in epigenetic changes that are inherited by daughter cells through failure to reinstate methylation and gene expression patterns after cell division. This provides a mechanistic explanation for the epigenetic reprogramming that has been reported in inflammation-associated diseases.

## Materials and Methods

### Study design

Treatment with glycine chloramine (GlyCl) occurred immediately after release from the cell cycle block. Growth rate was determined by live cell counts and flow cytometry was used to determine cell viability at 24, 48 and 72 hours post release. At 4 and 72 hours post-release, 5 x 10^6^ cells were harvested and stored for DNA and RNA extractions. All harvested cells underwent a slow centrifugation step (1000 x g for 5 min) to avoid harvesting late apoptotic or dead cells. This procedure was also performed with GlyCl treatment occurring 2 hours after release from cell cycle block. Independent biological replicates were obtained by repeating the experiments on different days with different oxidant preparations.

### Cell culture procedure

Jurkat E6.1 human suspension T-cell lymphoma cells were grown in RPMI 1640 medium supplemented with 10% v/v heat-inactivated FBS, 100 U/mL penicillin and 100 µg/mL streptomycin at 37° C in a humidified incubator with 5% CO_2_. Cell concentrations were not permitted to exceed 1 x 10^6^/mL and were not seeded at concentrations below 2.5 x 10^5^/mL. A log phase of growth was maintained by sub-culturing the cells every two days.

### Cell synchronization

Cell cycle arrest of Jurkat cells was performed by using an excess of thymidine as previously described (15, 72). Briefly, Jurkat cells (1 x 10^6^/mL) were treated with 1 mM thymidine for 18 hours. Cells were washed twice in PBS, resuspended in fresh media supplemented with 50 µM cytidine to promote progression out of G_1_ into S-phase. The cells were then split into either a treatment or control flasks and the treatment flask received a single bolus of 200 µM GlyCl while the control received the volumetric equivalent of PBS.

### Cell viability and proliferation

Cell viability was assessed using flow cytometry prior to thymidine block (pre-block), immediately after release from block (post-block) and at 4, 24, 48 and 72 h post release. Growth rate was determined by live cell counts at 24, 48 and 72 h post-release (current day count - previous day count)/previous day count x100. The percentage of viable cells was assessed by the exclusion of PI. Cell cycle transitions were observed by fixation with ice-cold 70% v/v ethanol and subsequent incubation with PI. At each major time point, cell proliferation profiles were compared between treatment and control by labelling cells with carboxyfluorescein diacetate succinimidyl ester (CFSE; Invitrogen, Auckland, New Zealand), as previously described (72).

### Glycine chloramine preparation

Glycine chloramine (GlyCl) was prepared immediately before treatment by adding hypochlorous acid dropwise to a 10 mM excess of glycine in an equivalent volume of PBS while gently vortexing. The concentration of glycine chloramine was determined using 5-thio-2-nitrobenzoic acid (TNB) by measuring the change in absorbance at 412 nm, using the molar extinction coeffcient for TNB (14; 100M^-1^ cm^-1^) and adjusting for the 1:2 stoichiometry of the reaction (GlyCl:TNB).

### Sample preparation for methylation array

DNA was extracted from 5 x 10^6^ Jurkat cells at 4 and 72-hour time points using the GeneJet genomic DNA purication kit (Thermo Scientific, Finland) as per the manufacturer’s protocol for cultured mammalian cells. Sodium bisulfite conversion on 1000 ng of extracted DNA from the 4 and 72 hour time points was performed using Zymo Research EZ DNA Methylation Kit (Zymo Research, Irvine,CA) according to the manufacturer’s specifications recommended for use on the Illumina Infinium MethylationEPIC 850K array (Illumina, Inc., San Diego, CA, USA). Genome wide DNA methylation profiles were assayed with the Illumina Infinium MethylationEPIC 850K kit, at the AgResearch Ltd (Invermay, New Zealand). Analysis was performed on all samples in a single batch.

### Data processing

Data analyses were performed using statistical software programs and packages as previously described (17, 73). R statistical program (www.R-project.org) and Minfi and Limma Bioconductor software packages were used for all statistical analyses. All packages, programs and workflows were based upon published scripts (74). All datasets passed quality control and were normalized using preprocessQuantile function within Minfi (75). However, one control sample had an abnormal methylation profile, which was likely due to low DNA quality and quantity. Inclusion of this sample may have skewed the results and artificially inflated statistical significance. Therefore, this replicate was removed from the analysis, and the data presented correspond to three replicates. Probes that produced detection *p*-values > 0.05 for 1% or more samples were considered unreliable and filtered out of the dataset. Probes identified as having polymorphic hybridizing potential and homology to common single nucleotide polymorphisms (SNPs) were also excluded (76). Probes were annotated using the IlluminaHumanMethylationEPICmanifest (77) and genomic locations were converted to GRCh38 using the Bioconductor package liftOver v.1.16.0. Multidimensional scaling of the top 1000 methylation values was performed using pairwise distance method for gene selection and hierarchical clustering was performed on a “minkowski” distance matrix calculated using the β-values for all probes, regardless of significance.

### Validation of the MethylationEPIC 850K array using BSAS

Bisulfite-based amplicon sequencing (BSAS) was utilized as an alternative DNA methylation detection technique, to validate CpG sites detected using the EPIC array, utilizing the protocols published by Nobel *et. al.*(78). Six CpG sites were selected for validation, based on their differential methylation status in the treatment group compared to the control group. Four of these CpGs originiated from the list of top differentially methylated CpGs at the four hour time point, and two from the list of differentially methylated regions. Three additional CpGs, which did not reach genome wide sequence level using the EPIC array were also included in the amplified target regions, bringing the total to nine CpGs, across differing levels of significance. Primers were designed to target regions of both methylated and non-methylated DNA, of an approximate size of 250 bp. The 5’ end of each primer sequence contained a 33 base pair sequence for Illumina barcoding during high-throughput sequencing. The forward and reverse primers were matched within two degrees (Celsius), for *T*_m_ which ranged from 52 °C -57 °C for all primers (Supplementary File 2). PCR protocol for bisulfite converted DNA followed that described in our previous publication (79).

PCR products were visualized by gel electrophoresis and purified using MagBio HighPrep™ PCR Clean-up System, according the manufacturer’s recommendations. DNA was eluted using 10 mM Tris pH 8.5 and quantified using the Qubit HS kit (Thermo Fisher). Preparation of the sequence libraries, and sequencing was performed by Massey Genome Service (New Zealand), using the Illumina MiSeq™.

Illumina MiSeq™ sequence reads were quality processed by the service provider according to their internal protocols, in brief this included removal of PhiX genomic DNA using Bowtie2, adapter removal using BBduk, and trimming using SolexaQA++. Sequences were further clipped using Trim Galore (RRID:SCR_011847) (Version 0.6.5) and aligned to bisulfite converted reference sequences using Bowtie2 (version 2.4.5). The Bismark package (80) was then used to align individual reads to their references sequence and produce counts for methylated and unmethylated CpG sites for validation. The methylation counts for each CpG were imported into R/minfi and combined into a methylset data frame and a coverage level greater than eight counts across unmethylated and methylated counts was verified. Beta and Methylation values were then calculated in Minfi, according to the protocols for the EPIC array. An identical statistical approach (general linear model, and adjustment for multiple testing) was then performed on the BSAS values, as with the EPIC array to detect differential methylation between treatment and control samples at the two time points. Epic array and BSAS beta values were visualized using a scatterplot with a regression line for all samples, and for individual groups. A Bland Altman analysis (81) was then used to visualize the agreement between the two methods.

### Data visualization and bioinformatic analysis of DNA methylation

Differentially variable positions (DVPs) were identified using the DiffVar algorithm within the missMethyl package (82, 83). The methylation status of each probe was calculated using normalized probe signals represented as methylation values (M-values) and β-values. M-values were generated within Minfi as the log_2_ ratio of the signal intensities of methylated probe divided by the unmethylated probe. β-values (average DNA methylation level for each probe) were used for data visualization and were generated by dividing the methylated probe signal with the sum of the methylated and unmethylated probe signals.

Differentially methylated positions that correlated with treatment were identified using a linear regression model within the Limma package, with adjustment for multiple testing. This was calculated by comparing the methylation measurements of the control samples with the treatment samples, for each of the two time points (4 hours post release, and 72 hours post release). Samples that originated from the same cell passage were treated as a biological replicate and the correlation coefficient between these samples was incorporated into the statistical design using Limma’s duppcorr function. The top most significant, differentially methylated CpG positions were identified using log fold change (logFC) weighted thresholds incorporated within the statistical design. Adjustment for multiple correction was performed using the “Benjamini, Hochberg” (BH) method within Limma (84). We prioritized significant results a filtering criteria that excluded probes with less than 10% mean difference in methylation between treatment and control samples, and probes that had a mean methylation value less than 10% or greater than 90% (in the control), as changes within the excluded ranges are unlikely to be of biological significance.

Pathway analysis was performed by comparison with the Kyoto Encyclopedia of Genes and Genomes (KEGG) (85) database using the missmethyl R package, with correction for probe bias. Differentially methylated regions were interrogated within the Minfi package using the statistical package DMRcate (86). A methylation differential cut-off of 10 was used, and unless stated otherwise significance was determined using a false discovery rate (FDR) of 0.05 in conjunction with a *p-*value cut-off of 0.05.

### Chromosomal probe position

To investigate where significant changes in DNA methylation were localised on the chromosomes, the probe position was classified as a percentage of chromosome length. The top most significant probes at the 4-hour time point were determined as probes that demonstrated log_2_FC > 1.5. This resulted in 2,968 significant differentially methylated probes. Each chromosome was divided into 10 bins as a percentage of the total length and the number of significant probes that were located in each bin was summed across all chromosomes. Simple Scaling Normalization was adapted from the LUMI pipeline to control for probe bias (87). The log_2_ of the total significant probe counts per region was subtracted from the log_2_ ratio of the total probes per region divided by total probes.

### RNA Sequencing

Cell aliquots were pelleted and frozen at -80°C for RNA extraction using the NucleoSpin RNA plus kit (Macherey-Nagel, Duren, Germany) according to the manufacturer’s recommendations. RNA quality was checked using the Agilent RNA Screen Tape and sequencing was outsourced to Custom Science (Auckland, New Zealand) according to their protocol for LncRNA-seq. For shipping, RNA was treated using the RNAstable Tube Kit (Biomatrica). Quality control and filtering of sequence data was performed by the service provider according to their internal protocols. In brief, this utilised cutadapt for adapter and low quality read filtering from the raw data (Pass Filter Data), removal of primers and adapters sequences, removal of the end sequences with base quality less than 20, removal of sequences with N content greater than 10%, and removal of reads with lengths less than 75bp after filtering. BWA (v. 0.7.12) (88) was also used to filter out host sequences based on host genome sequence. Filtered fastq files were then processed and aligned to the human reference genome (Gencode release 29) based on the GRCh38.p12, using the STAR spliced read aligner (v. 2.7.8a) (89), and a two-pass alignment mode. This resulted in a mean of ∼22 million (+/- ∼ 5 million) uniquely-mapped read pairs per sample. A gene matrix was built from the resulting Sam sorted, BAM files using GenomicAlignments (v. 1.26.0) (90) package in R (v. 4.0.3) and *summarizeoverlaps* within edgeR (v. 3.32.1) (91) utilising the “union” mode for paired-end, stranded sequencing. This was performed by making a gene level transcription database against the same gencode v. 29 annotation gene transfer file used for alignments. Filtering was performed by *filterByExpr* within edgeR and resulted in 20,374 genes retained for further analysis. An average of 29 million (range: 23 - 31 million) read pairs were successfully assigned in each sample. Differentially expressed genes were called using DESeq2 package (v. 1.30.1)(92) with adjusted p-value_cut-off of 0.05, and an internal log_2_FC of 0.2. The statistical model used to detect differential expression was consistent with the approach used for identifying differential methylation, where data was grouped by time and treatment. The effect of biological replicate was incorporated into the statistical model as a covariate, and differentially expressed genes were then identified by contrasting samples for each time point.

Gene set enrichment analyses for the DEGs at each time point were performed using goseq v. 1.44. This approach is significantly more stringent due to the ability to correct for gene length bias, and significance adjustment for multiple testing (93). GO categories with fewer than five genes in the final GO ranking list were excluded (94), with Gene Ontology and KEGG annotation drawn from the org.Hs.eg.db v. 3.10.0 database (95). The web based bioinformatic platform, WebGestalt (25) was used to assess the potential overrepresentation of significant results with biological disease, within the Online Mendelian Inheritance in Man (OMIM) database (96).

### Integrative analysis of DNA methylation and gene expression

To identify which significant promoter DNA methylation changes associated with significant gene expression changes, correlative analysis of the DNA methylation and RNAseq data was carried out.

The significant effect of GlyCl treatment was already determined for the gene expression and methylation data in the respective analyses, therefore, the correlation was performed on all paired samples, regardless of condition. The correlation was performed using methylation values (M-values) of significant CpGs against gene expression data, which was transformed and normalized using the DESeq2 ’regularized log’ transformation. The Pearson correlation (r) was calculated between the paired samples for methylation and RNA sequencing data for pairwise observations. Significance adjustment for multiple testing was determined using the “BH” method. Methylation probes were selected if they had an adjusted *p*-value less than 0.05 and absolute log_2_FC greater than 1.5. They were further filtered to those that occurred within regions 2000 bp upstream and 400 bp downstream of the transcription start site using the GenomicFeatures package v. 1.44.2 (90). Genomic locations were then converted using LiftOver, and methylation sites were annotated by location using the same gene transcription file previously used for RNA sequencing. Significant promoter methylation was merged with significant RNA transcripts using their ensemble gene identification number. This ensured that the promoter filtered methylation probes and RNA gene transcripts were accurately assigned to the same corresponding gene. In the instance where multiple CpGs bound in the same gene promoter region, the correlation was performed on each CpG value separately. Gene-enrichment analysis, of significant correlated genes was performed using web-based functional annotation bioinformatics tool, DAVID Resources 6.8 (97, 98). Goseq was not used for this analysis, as there was no need to prioritise gene length corrections. To investigate what biological disease outcomes might be influenced by the epigenetic regulation of gene expression following GlyCl treatment we assessed the significantly correlated genes using WebGestalt (25) against the OMIM database

## Funding

This work was supported by the Health Research Council of New Zealand and the Postgraduate Tassel Scholarship for Cancer Research.

## Conflict of Interest Disclosure

The authors declare no conflicts of interest

## Notes

https://apc01.safelinks.protection.outlook.com/?url=https%3A%2F%2Fwww.ncbi.nlm.nih.gov%2Fgeo%2Fquery%2Facc.cgi%3Facc%3DGSE199236&amp;data=04%7C01%7Caaron.stevens%40otago.ac.nz%7C7fc4340a2a184ed4317408da0d0e72f9%7C0225efc578fe4928b1579ef24809e9ba%7C0%7C0%7C637836652977513579%7CUnknown%7CTWFpbGZsb3d8eyJWIjoiMC4wLjAwMDAiLCJQIjoiV2luMzIiLCJBTiI6Ik1haWwiLCJXVCI6Mn0%3D%7C3000&amp;sdata=QzgOjGtkpxG7SGcSbGFEFssG4Xwthx9lU4fP8iZr7Z4%3D&amp;reserved=0

### Competing Interest Statement

The authors have declared no competing interest.

