## Supplemental Figures and Tables 1 for "Site-specific decreases in DNA methylation in replicating cells following exposure to oxidative stress"

**GlyCl exposure experiments 2 hours post-release from thymidine block**

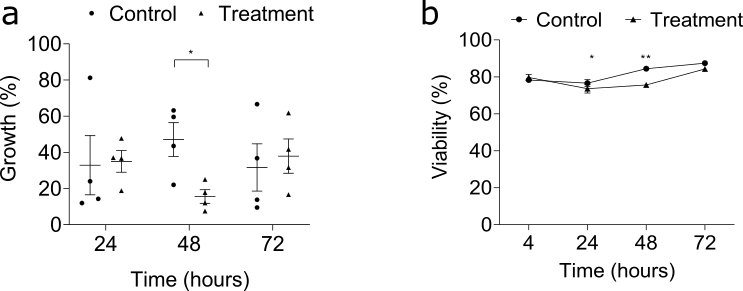

**Supplementary Figure 1: Viability and growth of GlyCl treated cells at 2 hours post release at each major time point.** Cell measures were conducted at 24, 48 and 72 hours post-release. Circles represent control samples and triangles represent treatment samples. Data are means and SE of 3 or 4 independent experiments. Significant differences were determined with paired t-tests and are denoted with asterisks * = p <0.05, ** = p <0.01. A) Cell growth after treatment B) Cellular viability after treatment.

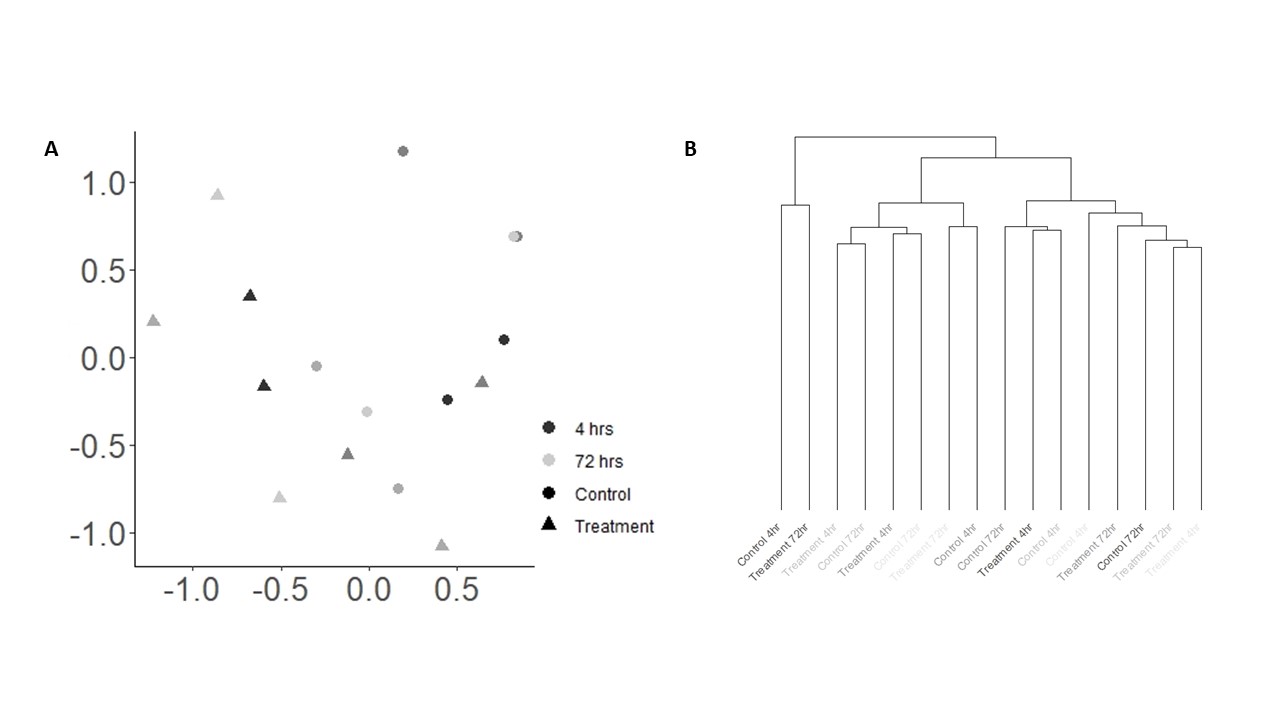

**Supplementary Figure 2:** **Unsupervised assessment of data variability in S-phase GlyCl exposure dataset**. (A) Multidimensional scaling of β-values, with the distances for leading Log_2_FCin dimension 1 represented on the x-axis and the leading Log_2_FC in dimension 2 are represented on the y-axis. Dots represent the control group, and triangles represent the treatment group. (B) Hierarchical clustering of β-values for all CpG sites. The relative change in β-values is represented on the y-axis and the individual samples are represented on the x-axis. Samples are coloured according to the replicate (three in total), where cells from each replicate originated from the same stock.

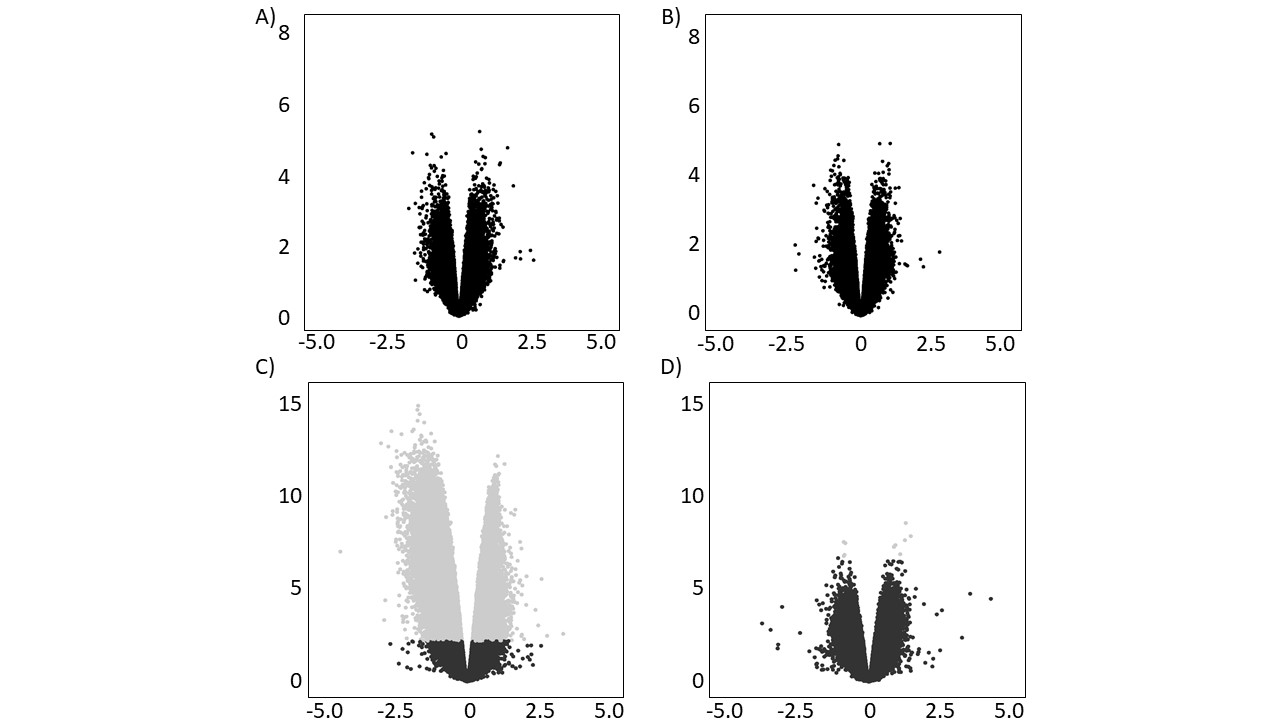

**Supplementary Figure 3:** **Volcano plots of the GlyCl T2 exposure dataset compared to the GlyCl T0 exposure dataset.** Volcano plot displaying log_2_ fold changes (M-values) for all probes on the x-axis versus statistical significance on the y-axis -log_10_ p-value. Grey dots represent probes with an adj. *p* <0.05. A) T2 GlyCl at the 4 hour time B) T2 GlyCl at the 72 hour time hour time point.

**Supplementary Table 1: Top 20 most significant differentially methylated CpGs at the 4 hour time point for GlyCl exposure 2 hours post release.**

|  | logFC | adj.P.Val | Direction | Gene | Chr | basepair |
| --- | --- | --- | --- | --- | --- | --- |
| cg01617955 | -1.12926 | 0.485001 | Down | LRBA; | chr4 | 1.52E+08 |
| cg17578309 | -0.82049 | 0.999985 | Down | TBC1D16 | chr17 | 77964703 |
| cg03791426 | -0.50481 | 0.999985 | Down | OLFML2B | chr1 | 1.62E+08 |
| cg21781319 | 0.827154 | 0.999985 | Up | MIR633;MIR548W | chr17 | 61020813 |
| cg00010266 | -0.84144 | 0.999985 | Down | MFSD3 | chr8 | 1.46E+08 |
| cg22627029 | 0.761977 | 0.999985 | Up | HLA-DRB6 | chr6 | 32520615 |
| cg20325092 | -0.88284 | 0.999985 | Down | CAPS2 | chr12 | 75699992 |
| cg22410262 | 0.715479 | 0.999985 | Up |  | chr11 | 64198213 |
| cg08743671 | -0.61903 | 0.999985 | Down | NDUFV1 | chr11 | 67378570 |
| cg08596797 | 0.62808 | 0.999985 | Up | SMAD6 | chr15 | 66994027 |
| cg06096382 | 0.629486 | 0.999985 | Up | HECW1 | chr7 | 43151725 |
| cg12650153 | 0.740166 | 0.999985 | Up | IKBKAP | chr9 | 1.12E+08 |
| cg22842599 | -0.42897 | 0.999985 | Down | LMTK2 | chr7 | 97749133 |
| cg24454158 | 0.541849 | 0.999985 | Up | FMN2 | chr1 | 2.4E+08 |
| cg07173352 | 1.163018 | 0.999985 | Up | LOC100652768;TAGLN | chr11 | 1.17E+08 |
| cg22423294 | -0.69394 | 0.999985 | Down | TFDP1 | chr13 | 1.14E+08 |
| cg04513006 | -0.62288 | 0.999985 | Down | ESRP2 | chr16 | 68270252 |
| cg21620282 | -0.72636 | 0.999985 | Down | CHGA | chr14 | 93389628 |
| cg17786993 | 1.145626 | 0.999985 | Up |  | chr4 | 1.75E+08 |
| cg22075325 | -0.5113 | 0.999985 | Down |  | chr14 | 1E+08 |

**Supplementary Table 2: Top 20 most significant differentially methylated CpGs at the 72 hour time point for GlyCl exposure 2 hours post release.**

|  | logFC | AveExpr | t | P.Value | adj.P.Val | B | Direction | Gene | chr | Basepair |
| --- | --- | --- | --- | --- | --- | --- | --- | --- | --- | --- |
| cg24328079 | 2.055593 | -4.28822 | 6.542062 | 4.87E-06 | 0.999989 | -1.39634 | Up | NMI | chr2 | 1.52E+08 |
| cg16582649 | -0.95206 | 4.434818 | -6.527 | 5.01E-06 | 0.999989 | -1.40196 | Down | FTHL17; | chrX | 31090073 |
| cg08039116 | 0.633455 | 4.6257 | 6.489899 | 5.38E-06 | 0.999989 | -1.4159 | Up | HECW1; | chr7 | 43152254 |
| cg24044884 | -1.04238 | 2.808218 | -6.15333 | 1.03E-05 | 0.999989 | -1.5482 | Down | NSG1; | chr4 | 4387109 |
| cg02252248 | -0.69744 | -0.09307 | -6.12618 | 1.09E-05 | 0.999989 | -1.55935 | Down |  | chr17 | 47073164 |
| cg17403512 | 0.532708 | 5.497289 | 6.059007 | 1.24E-05 | 0.999989 | -1.58724 | Up |  | chr15 | 28700396 |
| cg11781282 | -1.07452 | 2.613951 | -6.046 | 1.27E-05 | 0.999989 | -1.5927 | Down | UBXN4 | chr2 | 1.37E+08 |
| cg15094236 | -0.50763 | -3.00546 | -6.04581 | 1.27E-05 | 0.999989 | -1.59278 | Down | LOC285768 | chr6 | 1070389 |
| cg15798516 | 0.566694 | 3.770574 | 6.043259 | 1.28E-05 | 0.999989 | -1.59385 | Up | LOC647288 | chr13 | 75814711 |
| cg21970086 | -3.34998 | -1.88365 | -6.01009 | 1.37E-05 | 0.999989 | -1.60784 | Down | PRUNE2; | chr9 | 79318390 |
| cg16983152 | 0.502969 | 3.634839 | 5.964594 | 1.50E-05 | 0.999989 | -1.62721 | Up | BAT2 | chr6 | 31592053 |
| cg19063654 | -0.60112 | -2.33441 | -5.85336 | 1.87E-05 | 0.999989 | -1.67547 | Down | HAVCR2 | chr5 | 1.57E+08 |
| cg12041841 | 0.403774 | 4.088602 | 5.845258 | 1.90E-05 | 0.999989 | -1.67904 | Up |  | chr1 | 1.15E+08 |
| cg16712664 | 0.501416 | 5.082761 | 5.798163 | 2.09E-05 | 0.999989 | -1.69989 | Up |  | chr16 | 30787355 |
| cg12348202 | 0.820392 | 5.194545 | 5.78777 | 2.13E-05 | 0.999989 | -1.70453 | Up | PTPRN2; | chr7 | 1.58E+08 |
| cg18363721 | 0.690065 | -4.6168 | 5.784595 | 2.14E-05 | 0.999989 | -1.70595 | Up | SLC25A38 | chr3 | 39424765 |
| cg06658404 | 0.617751 | -4.23271 | 5.781932 | 2.16E-05 | 0.999989 | -1.70714 | Up |  | chr16 | 54408014 |
| cg13420422 | -0.72058 | 2.991216 | -5.74998 | 2.30E-05 | 0.999989 | -1.72148 | Down |  | chr7 | 1.41E+08 |
| cg25637520 | 0.620938 | 3.539763 | 5.737274 | 2.36E-05 | 0.999989 | -1.72721 | Up | KDM4B | chr19 | 5043247 |
| cg19030994 | -1.0474 | 2.12766 | -5.69482 | 2.57E-05 | 0.999989 | -1.74648 | Down | PDCD6 | chr5 | 273394 |

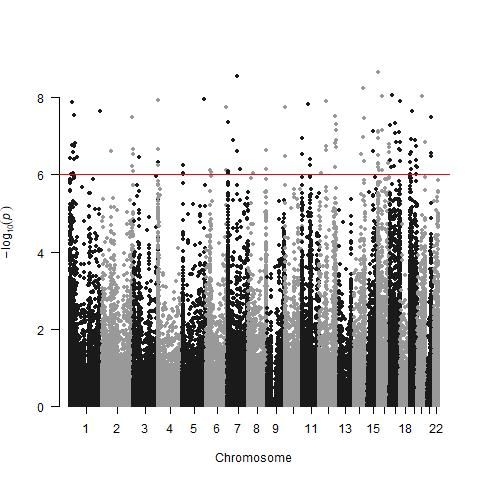

**Supplementary Figure 4: Manhattan plot of all CpG sites in the 4 hour time point.** All CpG sites are ordered per chromosome position along the x-axis, and p-values as the log_10_ (p-values) are presented on the y-axis. Genome-wide significance was determined using the “Benjamini, Hochberg" method within Limma is represented by the horizontal red line.

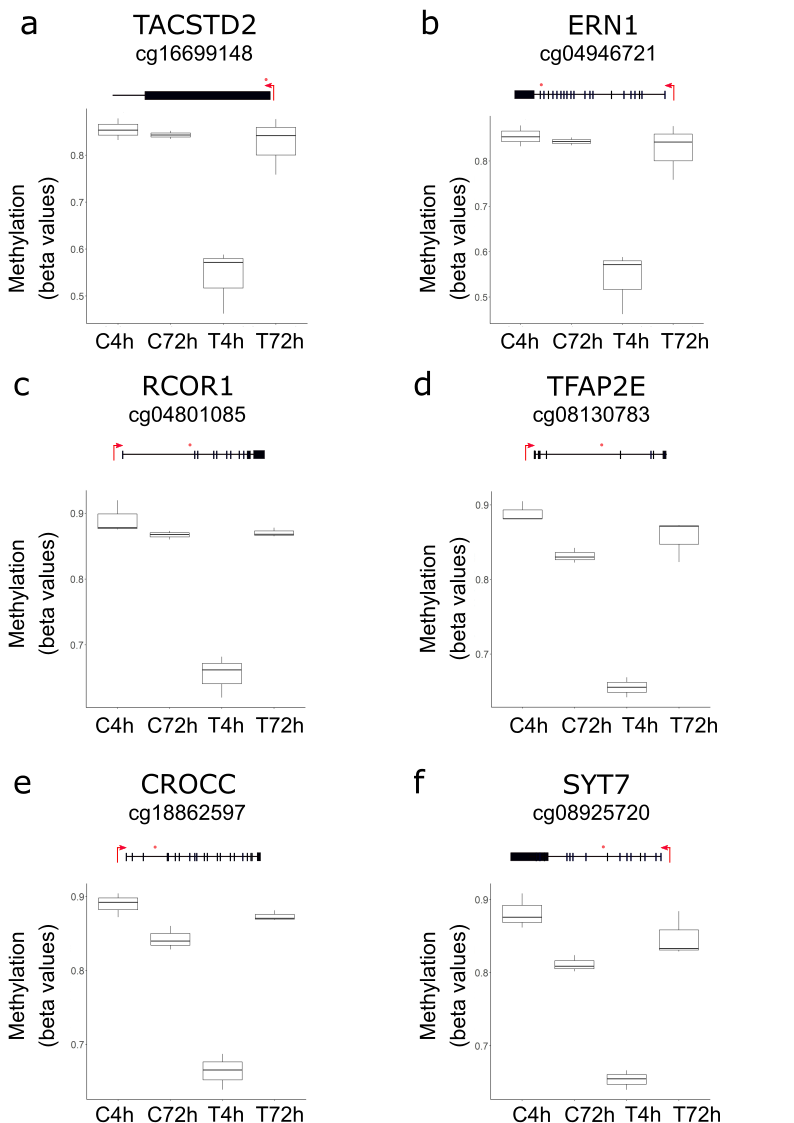

**Supplementary Figure 5: Box plots showing β-values of a subset of the top differentially methylated CpG sites at the 4 hour time point.** CpG sites were selected that demonstrated the largest, significant fold changes at the 4 hour time point. β-values are represented on the y-axis and group on the x-axis. The CpG sites are marked with an orange dot in the representative scheme placed above each graph. The transcriptional start site is marked by a red arrow and exons are marked with vertical lines and boxes.**a**, chr1:59043255; **b**, chr17:62131780; **c**, chr14:103150012; **d**, chr1:36056577; **e**, chr1:17265457; **f**, chr11:61314936

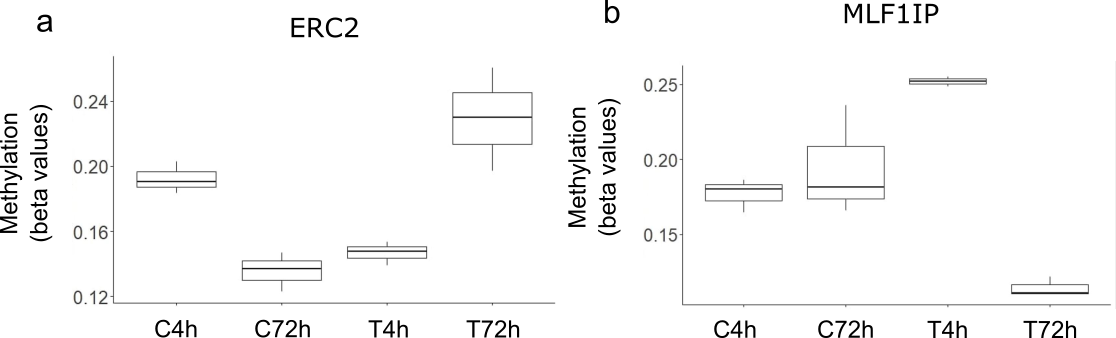

**Supplementary Figure 6: Box plots showing the β-values of the two differentially methylated CpG sites at both the 4 hour and 72 hour time points.** β-values are represented on the y-axis and group on the x-axis**. a,** ERC2 **b,** MLF1IP

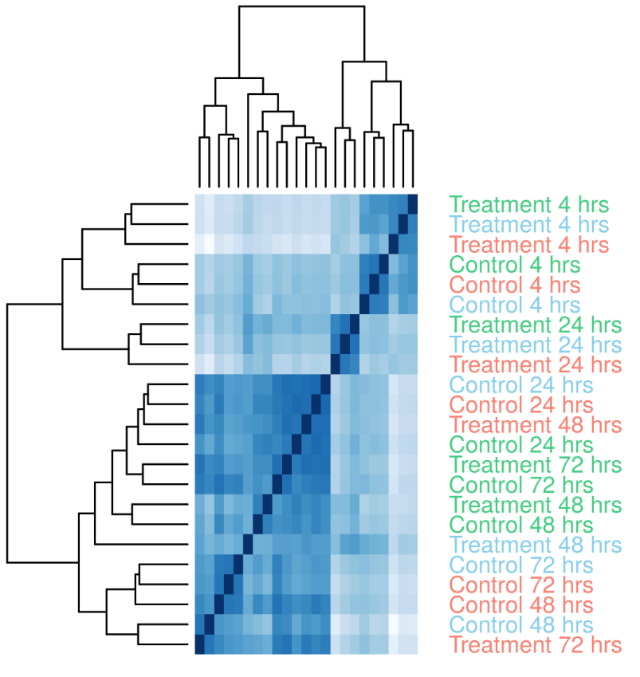
**a**

**b**

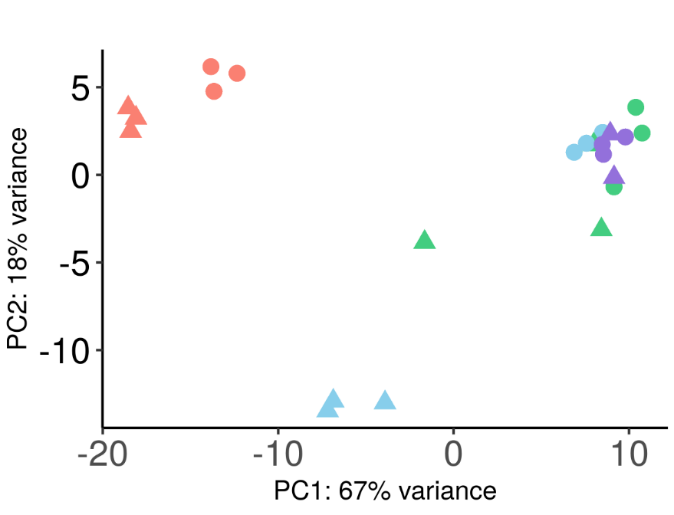

**Supplementary Figure 7. Unsupervised assessment of RNA variability in the GlyCl exposure dataset**. Hierarchical clustering and principle component analysis of regularized log transformed, filtered counts. The 4 hour control samples grouped closely with the 4 hour treatment group, which likely reflects that at this time point both cell populations were in the S-phase of replication. Both PCA and hierarchical clustering indicated that there were less substantial differences between samples from the 48 hr and 72 h samples, which did not cluster into distinct groups. This pattern was reflected in the significance and magnitude of differentially expressed genes observed at each time point. **a,** Heatmap representing hierarchical clustering at the sample level with samples coloured according to replicate. **b,** Principle component anlaysis with treatment attributed to PC1 and time attribute to PC2. Dots represent control samples and triangles represent the treatment samples. Red represents samples from the 4 hr time point, light blue represents the 24 hr time point, green represents the 48 hr time point and purple represents the 72 hr time point.

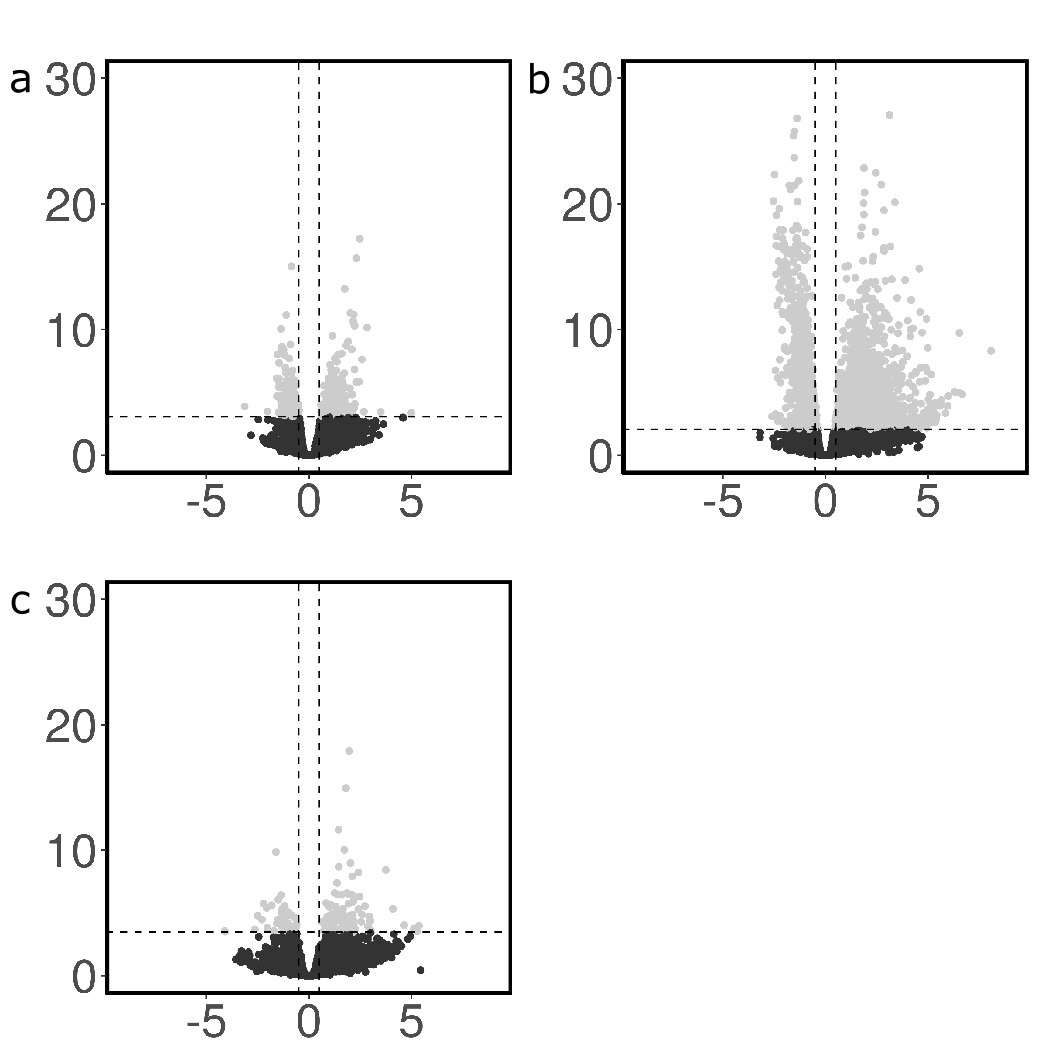

**Supplementary Figure 8:** **Significance plots of each time point for RNA sequencing analysis.** At the 4 hour time point there was a relatively even spread in the number of genes showing increased and decreased expression, and these exhibited a similar significance and magnitude of change. The largest effect on differential gene expression occurred at the 24-hour time point, where the genes that showed decreased expression corresponded with smaller *p*-values and had a lower magnitude of change. At the 48-hour time point, the magnitude of changes and number of significant changes substantially reduced, which is consistent with the pattern of DNA methylation changes. The majority of the remaining significant changes corresponded with decreased gene expression**.** Figure shows volcano plot displaying log_2_ fold changes for all probes on the x-axis versus statistical significance on the y-axis -log_10_ p-value. Grey dots represent probes with an adj. *p* <0.05. **a,** 4 hr time **b,** 24 hr time point. **c,** 48 hr time point.

**Supplementary Table 3: Differentially variable probes (DVPs) ordered by t-statistic significance that were common in both the 4 and 72 hour time points.**

|  | 4 hours | | |  | 72 hours | | | |  |
| --- | --- | --- | --- | --- | --- | --- | --- | --- | --- |
| Probe ID | LVR | t | adj.*p*-val. |  | LVR | | t | adj.*p*-val. | Gene |
| cg06521531 | 6.72 | 7.73 | 0.01 |  | -5.85 | | -8.26 | 0.01 | FHL1 |
| **cg13981054** | 6.07 | 7.61 | 0.01 |  | 6.48 | | 7.66 | 0.01 |  |
| **cg10568558** | -3.31 | -7.03 | 0.01 |  | -3.64 | | -5.56 | 0.03 | UBE4B |
| **cg14612785** | -7.35 | -6.52 | 0.02 |  | -4.41 | | -5.34 | 0.04 | EIF4E3 |
| cg27445386 | 2.31 | 6.26 | 0.02 |  | -3.99 | | -5.10 | 0.04 | CNPY3 |
| cg13835576 | 5.84 | 6.25 | 0.02 |  | -6.10 | | -6.50 | 0.02 | BLK |
| cg00276792 | -8.36 | -6.21 | 0.02 |  | | 6.87 | 5.19 | 0.04 |  |
| **cg27560282** | -2.85 | -5.91 | 0.02 |  | | -6.10 | -5.11 | 0.04 | ICK |
| cg24598449 | 3.64 | 5.87 | 0.02 |  | | -3.24 | -4.98 | 0.04 | FBXO6 |
| cg09016212 | 4.51 | 5.87 | 0.02 |  | | -2.85 | -8.41 | 0.01 | GRASP |
| cg03913456 | 4.70 | 5.84 | 0.02 |  | | -5.15 | -6.21 | 0.03 | NCAPH |
| cg00360072 | 2.16 | 5.78 | 0.02 |  | | -2.29 | -5.35 | 0.04 | HDAC4 |
| cg21267439 | -3.67 | -5.73 | 0.02 |  | | 5.24 | 5.48 | 0.04 | NT5DC1 |
| **cg15372625** | -2.91 | -5.72 | 0.02 |  | | -3.02 | -5.19 | 0.04 |  |
| cg05352535 | 4.96 | 5.68 | 0.02 |  | | -3.60 | -6.36 | 0.02 | PVRL3 |
| cg23161691 | 7.55 | 5.66 | 0.02 |  | | -3.58 | -5.55 | 0.03 | BAHCC1 |
| cg21388527 | -2.68 | -5.63 | 0.02 |  | | 3.25 | 8.48 | 0.01 | EPHA6 |
| cg18558388 | 5.22 | 5.62 | 0.02 |  | | -6.73 | -5.23 | 0.04 | SAP30L |
| cg00823357 | 4.98 | 5.59 | 0.02 |  | | -3.25 | -5.11 | 0.04 | EFNB1 |
| cg23444251 | 3.01 | 5.50 | 0.02 |  | | -3.43 | -5.04 | 0.04 | FGFR2 |
| cg16688031 | 3.69 | 5.40 | 0.02 |  | | -2.11 | -5.66 | 0.03 |  |
| **cg20898865** | -2.18 | -5.28 | 0.02 |  | | -3.09 | -8.18 | 0.01 |  |
| cg03324815 | -1.85 | -5.26 | 0.02 |  | | 2.33 | 5.20 | 0.04 | MZF1 |

**LVR, Log Variance Ratio

**Probes that demonstrated a consistent direction of change are in bold typeface.

**Supplementary Table 4: Top 20 most significant differentially methylated CpGs at the 72 hour time point using a log_2_FC cut off of ~ 1.**

| Probe ID | logFC | adj.*p*.Val | Direction | Gene | CHR | Basepair |
| --- | --- | --- | --- | --- | --- | --- |
| cg00297451 | 1.352419 | 0.011303 | Up | ARHGAP11A | 15 | 32908400 |
| cg16381597 | 1.533733 | 0.022432 | Up | ZNF142;BCS1L | 2 | 219524462 |
| cg27633753 | 1.32058 | 0.022432 | Up | OCRL | X | 128674222 |
| cg03343631 | -0.91007 | 0.022432 | Down | RASA3 | 13 | 114764489 |
| cg18538954 | -0.85446 | 0.022432 | Down |  | 16 | 56511477 |
| cg14058432 | 0.972789 | 0.02329 | Up | PELI2 | 14 | 56584702 |
| cg13722120 | 0.918361 | 0.024522 | Up | ERC2 | 3 | 56487225 |
| cg07171505 | 1.150163 | 0.047183 | Up | C16orf70 | 16 | 67144003 |
| cg07934941 | -0.88631 | 0.047183 | Down | MLF1IP | 4 | 185654596 |
| cg06851253 | -0.90945 | 0.048603 | Down | CHCHD4;TMEM43 | 3 | 14166363 |
| cg21975824 | -1.11975 | 0.056204 | Down | RBM26 | 13 | 79980660 |
| cg13055288 | 0.897001 | 0.056511 | Up | SLC7A2 | 8 | 17396259 |
| cg13953717 | 0.694856 | 0.056511 | Up | MIR99AHG | 21 | 17657943 |
| cg08046288 | 1.144429 | 0.056511 | Up |  | 1 | 199082308 |
| cg11216153 | -0.9647 | 0.056511 | Down | HADH | 4 | 108921304 |
| cg02768742 | 1.084081 | 0.056511 | Up | LARS | 5 | 145562530 |
| cg19407454 | -0.68892 | 0.056511 | Down | IQCE | 7 | 2643258 |
| cg03050188 | 1.20667 | 0.056511 | Up |  | 2 | 11531606 |
| cg11979743 | -0.97845 | 0.059587 | Down | FAM110A | 20 | 814510 |
| cg16603817 | 0.769654 | 0.059587 | Up |  | 5 | 148827511 |

**Supplementary Table 5: Significant DMR observed at 4 hours after GlyCl exposure obtained using the dmrcate algorithm within R.**

| Chr | start | end | width | no.cpgs | minfdr | maxβfc | meanβfc | overlapping.promoters |
| --- | --- | --- | --- | --- | --- | --- | --- | --- |
| chr16 | 1813796 | 1818819 | 5024 | 18 | 3E-79 | -2E-01 | -1E-01 | MAPK8IP3-014, MAPK8IP3-015, MAPK8IP3-013, MAPK8IP3-012, MAPK8IP3-011, MAPK8IP3-010, MAPK8IP3-009 |
| chr1 | 36037583 | 36039885 | 2303 | 16 | 2E-183 | -2E-01 | -1E-01 | TFAP2E-001, RP4-728D4.2-002 |
| chr22 | 50644361 | 50647567 | 3207 | 16 | 6E-103 | -2E-01 | -1E-01 | SELO-002, RP3-402G11.28-001, RP3-402G11.27-001 |
| chr17 | 981518 | 984184 | 2667 | 16 | 2E-197 | -2E-01 | -1E-01 | ABR-026, ABR-025 |
| chr21 | 43372534 | 43374650 | 2117 | 14 | 2E-185 | -2E-01 | -1E-01 | C2CD2-001, C2CD2-009 |
| chr20 | 61296949 | 61299808 | 2860 | 14 | 5E-98 | -2E-01 | -1E-01 | RP11-93B14.5-002, RP11-93B14.5-001, RP11-93B14.5-003, SLCO4A1-007, SLCO4A1-008, SLCO4A1-003, SLCO4A1-004, SLCO4A1-001, SLCO4A1-002 |
| chr1 | 9793553 | 9796095 | 2543 | 12 | 2E-71 | -2E-01 | -1E-01 | CLSTN1-006 |
| chr11 | 606568 | 609744 | 3177 | 11 | 8E-90 | -2E-01 | -1E-01 |  |
| chr8 | 1.42E+08 | 1.42E+08 | 2742 | 11 | 8E-66 | -2E-01 | -1E-01 | AGO2-007 |
| chr17 | 8382941 | 8386145 | 3205 | 11 | 1E-102 | -2E-01 | -1E-01 | MYH10-009 |
| chr11 | 984447 | 986709 | 2263 | 10 | 5E-64 | -2E-01 | -1E-01 | AP2A2-014, AP2A2-015 |
| chr21 | 45160229 | 45161603 | 1375 | 10 | 4E-94 | -2E-01 | -1E-01 | PDXK-005, PDXK-011, PDXK-010, PDXK-014 |
| chr15 | 74314149 | 74315710 | 1562 | 10 | 5E-129 | -2E-01 | -1E-01 | PML-018, PML-022, PML-023, PML-017, PML-019 |
| chr19 | 1624494 | 1627485 | 2992 | 10 | 4E-81 | -1E-01 | -1E-01 | NA |
| chr2 | 2.43E+08 | 2.43E+08 | 2037 | 9 | 4E-85 | -2E-01 | -1E-01 | ING5-011 |
| chr16 | 69969137 | 69971338 | 2202 | 9 | 2E-53 | -2E-01 | -1E-01 | WWP2-018, WWP2-019 |
| chr16 | 2500696 | 2503678 | 2983 | 9 | 1E-59 | -2E-01 | -1E-01 | RP11-715J22.3-001 |
| chr17 | 17464853 | 17465682 | 830 | 8 | 1E-119 | -1E-01 | -1E-01 | NA |
| chr22 | 46593349 | 46594435 | 1087 | 8 | 3E-96 | -1E-01 | -1E-01 | PPARA-203 |
| chr19 | 3522506 | 3523009 | 504 | 8 | 3E-131 | -2E-01 | -1E-01 | FZR1-003, FZR1-002, SNORD38.4-201, FZR1-004 |
| chr16 | 4748507 | 4749556 | 1050 | 8 | 3E-143 | -2E-01 | -1E-01 | ANKS3-028, ANKS3-009, ANKS3-010 |
| chr17 | 79166676 | 79168726 | 2051 | 8 | 4E-40 | -1E-01 | -1E-01 | AZI1-004, AZI1-007 |
| chr7 | 1.57E+08 | 1.57E+08 | 1672 | 8 | 3E-69 | -2E-01 | -1E-01 | NA |
| chr16 | 69368764 | 69370169 | 1406 | 8 | 8E-83 | -2E-01 | -1E-01 | RP11-343C2.12-001 |
| chr16 | 941948 | 943632 | 1685 | 8 | 4E-59 | -2E-01 | -1E-01 | LMF1-015 |
| chr17 | 79215047 | 79216949 | 1903 | 7 | 6E-77 | -2E-01 | -1E-01 | C17orf89-005, C17orf89-004, C17orf89-003, C17orf89-002, C17orf89-007 |
| chr17 | 74386817 | 74387581 | 765 | 7 | 5E-92 | -2E-01 | -1E-01 | NA |
| chr7 | 6448148 | 6449981 | 1834 | 7 | 1E-62 | -2E-01 | -1E-01 | NA |
| chr17 | 78312907 | 78314035 | 1129 | 7 | 3E-62 | -1E-01 | -1E-01 | RNF213-201, RNF213-014 |
| chr17 | 76421538 | 76422976 | 1439 | 7 | 2E-77 | -2E-01 | -1E-01 | DNAH17-005, DNAH17-012, AC061992.1-201 |
| chr22 | 50664592 | 50666063 | 1472 | 7 | 6E-75 | -2E-01 | -1E-01 | TUBGCP6-004, TUBGCP6-005, TUBGCP6-003 |
| chr11 | 3173868 | 3175636 | 1769 | 7 | 1E-51 | -2E-01 | -1E-01 | NA |
| chr17 | 78326557 | 78328252 | 1696 | 7 | 3E-68 | -2E-01 | -1E-01 | RNF213-004, RNF213-008 |
| chr7 | 1.57E+08 | 1.57E+08 | 894 | 7 | 6E-56 | -1E-01 | -1E-01 | NA |
| chr13 | 24796915 | 24798240 | 1326 | 7 | 2E-40 | -1E-01 | -1E-01 | SPATA13-002 |
| chr16 | 57562120 | 57563395 | 1276 | 6 | 1E-83 | -2E-01 | -1E-01 | NA |
| chr16 | 87525178 | 87525542 | 365 | 6 | 2E-94 | -2E-01 | -2E-01 | ZCCHC14-001, RP11-482M8.1-002, RP11-482M8.1-001, ZCCHC14-002 |
| chr11 | 1.17E+08 | 1.17E+08 | 578 | 6 | 8E-84 | -2E-01 | -1E-01 | SIDT2-019, SIDT2-020, SIDT2-018, SIDT2-004 |
| chr16 | 84822583 | 84823224 | 642 | 6 | 4E-64 | -1E-01 | -1E-01 | NA |
| chr8 | 1.42E+08 | 1.42E+08 | 489 | 6 | 9E-66 | -2E-01 | -1E-01 | NA |
| chr20 | 60976987 | 60978651 | 1665 | 6 | 7E-63 | -2E-01 | -1E-01 | NA |
| chr12 | 1.25E+08 | 1.25E+08 | 642 | 6 | 2E-51 | -1E-01 | -1E-01 | SCARB1-009 |
| chr8 | 1.45E+08 | 1.45E+08 | 1407 | 6 | 3E-45 | -2E-01 | -1E-01 | NA |
| chr21 | 45547032 | 45548059 | 1028 | 6 | 1E-64 | -2E-01 | -1E-01 | PWP2-006, PWP2-005 |
| chr5 | 1033518 | 1034624 | 1107 | 6 | 5E-78 | -2E-01 | -1E-01 | NKD2-004, NKD2-005, NKD2-006 |
| chr7 | 6191780 | 6192094 | 315 | 6 | 1E-61 | -2E-01 | -1E-01 | NA |
| chr16 | 75563009 | 75564221 | 1213 | 6 | 3E-64 | -1E-01 | -1E-01 | CHST5-201 |
| chr17 | 79204738 | 79206371 | 1634 | 6 | 2E-79 | -2E-01 | -1E-01 | AC027601.1-001, ENTHD2-013, ENTHD2-012 |
| chr12 | 1.32E+08 | 1.32E+08 | 922 | 6 | 1E-47 | -1E-01 | -1E-01 | PUS1-011 |
| chr16 | 2647662 | 2648415 | 754 | 6 | 8E-40 | -1E-01 | -1E-01 | PDPK1-017, CTD-3126B10.1-001, PDPK1-018 |
| chr1 | 2205213 | 2205902 | 690 | 6 | 3E-34 | -2E-01 | -1E-01 | NA |
| chr2 | 2.4E+08 | 2.4E+08 | 661 | 6 | 1E-44 | -2E-01 | -1E-01 | NA |
| chr4 | 858299 | 859471 | 1173 | 5 | 6E-85 | -2E-01 | -1E-01 | GAK-010, GAK-011 |
| chr19 | 46056709 | 46057298 | 590 | 5 | 4E-76 | -2E-01 | -1E-01 | NA |
| chr14 | 1.03E+08 | 1.03E+08 | 1072 | 5 | 7E-74 | -1E-01 | -1E-01 | NA |
| chr7 | 1.51E+08 | 1.51E+08 | 625 | 5 | 2E-74 | -2E-01 | -1E-01 | AGAP3-005, AGAP3-013, AGAP3-012 |
| chr1 | 6333502 | 6334700 | 1199 | 5 | 1E-54 | -1E-01 | -1E-01 | NA |
| chr3 | 47450718 | 47451026 | 309 | 5 | 4E-57 | -1E-01 | -1E-01 | PTPN23-009 |
| chr19 | 4688549 | 4689839 | 1291 | 5 | 2E-59 | -2E-01 | -1E-01 | DPP9-016, DPP9-014 |
| chr7 | 1.02E+08 | 1.02E+08 | 616 | 5 | 1E-60 | -1E-01 | -1E-01 | RP11-514P8.2-001, ORAI2-003, ORAI2-005 |
| chr6 | 52267928 | 52268951 | 1024 | 5 | 2E-76 | -2E-01 | -1E-01 | NA |
| chr16 | 31235535 | 31236039 | 505 | 5 | 9E-57 | -2E-01 | -1E-01 | RP11-388M20.9-001 |
| chr16 | 2497840 | 2499223 | 1384 | 5 | 4E-47 | -1E-01 | -1E-01 | NA |
| chr11 | 3122878 | 3123614 | 737 | 5 | 4E-56 | -2E-01 | -1E-01 | OSBPL5-017 |
| chr2 | 2.42E+08 | 2.42E+08 | 793 | 5 | 5E-59 | -2E-01 | -1E-01 | PASK-013 |
| chr19 | 5243839 | 5244269 | 431 | 5 | 3E-47 | -2E-01 | -1E-01 | NA |
| chr20 | 60881152 | 60881634 | 483 | 5 | 2E-63 | -2E-01 | -1E-01 | RP11-157P1.4-001, ADRM1-004 |
| chr19 | 46443657 | 46444050 | 394 | 5 | 2E-51 | -1E-01 | -1E-01 | NA |
| chr17 | 78354382 | 78355552 | 1171 | 5 | 2E-45 | -1E-01 | -1E-01 | RNF213-006, RNF213-017 |
| chr17 | 78853966 | 78854418 | 453 | 5 | 3E-70 | -2E-01 | -1E-01 | NA |
| chr1 | 6149049 | 6149517 | 469 | 5 | 8E-58 | -1E-01 | -1E-01 | NA |
| chr20 | 60733611 | 60733814 | 204 | 5 | 7E-41 | -1E-01 | -1E-01 | SS18L1-201 |
| chr1 | 6336239 | 6337401 | 1163 | 5 | 2E-45 | -1E-01 | -1E-01 | NA |
| chr17 | 48431653 | 48434445 | 2793 | 5 | 2E-30 | -1E-01 | -1E-01 | XYLT2-005, XYLT2-006, XYLT2-007 |
| chr17 | 77963988 | 77964734 | 747 | 5 | 3E-43 | -2E-01 | -1E-01 | CTD-2529O21.1-001 |
| chr4 | 955469 | 956377 | 909 | 5 | 1E-36 | -1E-01 | -1E-01 | DGKQ-005 |
| chr16 | 84828558 | 84829007 | 450 | 5 | 8E-75 | -2E-01 | -1E-01 | NA |
| chr11 | 64029923 | 64031188 | 1266 | 5 | 3E-57 | -2E-01 | -1E-01 | PLCB3-003 |
| chr11 | 66475076 | 66475721 | 646 | 5 | 3E-54 | -2E-01 | -1E-01 | NA |
| chr7 | 893267 | 894202 | 936 | 5 | 1E-29 | -1E-01 | -1E-01 | SUN1-033, SUN1-032, SUN1-010 |
| chr11 | 987821 | 990566 | 2746 | 5 | 1E-59 | -3E-01 | -1E-01 | NA |
| chr12 | 1.33E+08 | 1.33E+08 | 1638 | 5 | 4E-51 | -2E-01 | -1E-01 | NA |
| chr16 | 2294493 | 2296200 | 1708 | 5 | 7E-35 | -1E-01 | -1E-01 | ECI1-008 |
| chr7 | 1.59E+08 | 1.59E+08 | 723 | 5 | 2E-32 | -1E-01 | -1E-01 | VIPR2-003 |
| chr17 | 7811107 | 7812665 | 1559 | 5 | 2E-42 | -3E-01 | -1E-01 | CHD3-010, CHD3-011, CHD3-005, SCARNA21-201, CHD3-017, CHD3-016 |

**Supplementary Table 6: Significant DMR observed at 72 hours after GlyCl exposure obtained using the dmrcate algorithm within R.**

| Chr | start | end | width | no.cpgs | minfdr | maxβfc | meanβfc | overlapping.promoters |
| --- | --- | --- | --- | --- | --- | --- | --- | --- |
| chr20 | 1276826 | 1277814 | 989 | 5 | 9.09E-05 | 0.355145 | 0.059969 | NA |
| chr7 | 1E+08 | 1E+08 | 794 | 5 | 2.11E-06 | 0.206638 | 0.051375 | ZAN-006, ZAN-001, ZAN-004, ZAN-002, ZAN-003, ZAN-005, ZAN-201, ZAN-202, ZAN-203, ZAN-204 |
| chr8 | 72459908 | 72460431 | 524 | 5 | 0.013293 | -0.44966 | -0.09736 | RP11-1102P16.1-002, RP11-1102P16.1-001 |
| chr10 | 12390599 | 12391599 | 1001 | 6 | 0.005069 | -0.35303 | -0.06175 | CAMK1D-002, CAMK1D-001, CAMK1D-003 |

**Supplementary Table 7. Top 100 most significant differentially expressed genes at the 4 hour time point, ordered by fold change.**

| Log_2_FC | adj.*p*.Val | Gene | CHR | start | end |
| --- | --- | --- | --- | --- | --- |
| 4.99 | 0.032 | AC019211.1 | chr2 | 220105656 | 220450778 |
| 3.50 | 0.028 | NXPH3 | chr17 | 49575858 | 49583827 |
| -3.13 | 0.014 | FBXL22 | chr15 | 63597353 | 63602428 |
| 2.83 | 0.000 | LHFPL4 | chr3 | 9498361 | 9553802 |
| 2.68 | 0.027 | PRB3 | chr12 | 11265924 | 11269805 |
| 2.59 | 0.000 | MIR99AHG | chr21 | 15928296 | 16645065 |
| 2.48 | 0.000 | ALPK3 | chr15 | 84816680 | 84873482 |
| 2.45 | 0.001 | FOXR1 | chr11 | 118971707 | 118981291 |
| 2.31 | 0.000 | AC244131.2 | chr12 | 11212219 | 11251389 |
| 2.30 | 0.001 | TMEM186 | chr16 | 8780384 | 8797648 |
| 2.23 | 0.010 | LINC02204 | chr15 | 70570958 | 70586606 |
| 2.23 | 0.000 | SLC6A13 | chr12 | 220621 | 262873 |
| 2.22 | 0.000 | COL20A1 | chr20 | 63293186 | 63334851 |
| 2.22 | 0.016 | TREHP1 | chr11 | 118688033 | 118690599 |
| 2.21 | 0.000 | CFAP161 | chr15 | 81007033 | 81149175 |
| 2.18 | 0.000 | HIST3H3 | chr1 | 228424845 | 228425325 |
| 2.16 | 0.000 | AC008897.2 | chr5 | 75320155 | 75336914 |
| 2.13 | 0.003 | CYP27A1 | chr2 | 218781749 | 218815293 |
| 2.10 | 0.024 | AC090844.3 | chr17 | 40012226 | 40014705 |
| 2.09 | 0.000 | AC011451.3 | chr19 | 9383581 | 9407055 |
| 2.02 | 0.000 | TEX29 | chr13 | 111316184 | 111344249 |
| -2.02 | 0.027 | IGHV4-4 | chr14 | 106011922 | 106012420 |
| 1.99 | 0.001 | AL512306.3 | chr1 | 204603035 | 204616565 |
| 1.98 | 0.019 | AC104170.1 | chr1 | 51518309 | 51561629 |
| 1.97 | 0.034 | LINC01563 | chr17 | 21075556 | 21090615 |
| 1.92 | 0.038 | LINC00346 | chr13 | 110863987 | 110870251 |
| 1.91 | 0.000 | IL12RB2 | chr1 | 67307364 | 67397090 |
| 1.89 | 0.013 | SYBU | chr8 | 109573978 | 109691791 |
| 1.83 | 0.003 | SLC22A14 | chr3 | 38282294 | 38318575 |
| 1.82 | 0.000 | AC096719.1 | chr4 | 21949015 | 22330330 |
| 1.79 | 0.018 | LINC01562 | chr1 | 51195095 | 51235096 |
| 1.78 | 0.003 | MAEL | chr1 | 166975582 | 167022214 |
| 1.78 | 0.003 | PRB1 | chr12 | 11351823 | 11395566 |
| 1.78 | 0.024 | ZNF532 | chr18 | 58862600 | 58986480 |
| 1.78 | 0.018 | AC010325.1 | chr19 | 50776141 | 50793142 |
| 1.78 | 0.009 | AC012574.1 | chr8 | 23707141 | 23789487 |
| 1.77 | 0.022 | LINC02145 | chr5 | 6310441 | 6339884 |
| 1.76 | 0.026 | SMPD4P1 | chr22 | 20602595 | 20656914 |
| 1.75 | 0.000 | KCND3 | chr1 | 111770662 | 111989155 |
| 1.74 | 0.001 | ZNF843 | chr16 | 31432593 | 31443160 |
| 1.72 | 0.000 | ACSBG1 | chr15 | 78167468 | 78245688 |
| 1.70 | 0.001 | LRRN2 | chr1 | 204617170 | 204685733 |
| 1.69 | 0.012 | PM20D1 | chr1 | 205828022 | 205850132 |
| 1.67 | 0.010 | GMPR | chr6 | 16238580 | 16295549 |
| 1.66 | 0.012 | RPL3L | chr16 | 1943974 | 1957606 |
| 1.66 | 0.001 | FAM71F2 | chr7 | 128672288 | 128687872 |
| 1.65 | 0.001 | SLC35D2 | chr9 | 96313444 | 96383710 |
| 1.65 | 0.022 | AC074351.1 | chr7 | 54759425 | 54804928 |
| 1.64 | 0.000 | AC008758.4 | chr19 | 12379189 | 12401274 |
| 1.63 | 0.006 | AL133216.1 | chr10 | 38453181 | 38466176 |
| 1.63 | 0.026 | LINC01277 | chr6 | 142966421 | 143038077 |
| 1.62 | 0.038 | AC005020.1 | chr7 | 99638242 | 99638767 |
| 1.61 | 0.022 | COL8A1 | chr3 | 99638475 | 99799226 |
| 1.61 | 0.003 | ACOT12 | chr5 | 81330005 | 81394179 |
| 1.60 | 0.000 | ABCC6 | chr16 | 16148928 | 16223522 |
| 1.60 | 0.003 | SLC22A1 | chr6 | 160121789 | 160158718 |
| 1.59 | 0.038 | LINC00578 | chr3 | 177441921 | 177752305 |
| 1.59 | 0.023 | TEX48 | chr9 | 114666434 | 114682066 |
| 1.59 | 0.020 | DPYS | chr8 | 104330324 | 104467053 |
| 1.58 | 0.016 | PRRT1B | chr9 | 131545514 | 131558620 |
| 1.58 | 0.007 | CFAP100 | chr3 | 126394939 | 126436556 |
| -1.57 | 0.000 | IGHV1-3 | chr14 | 106005095 | 106005574 |
| 1.56 | 0.001 | AC019186.1 | chr2 | 157725708 | 157736005 |
| 1.56 | 0.006 | AC068305.2 | chr12 | 58920639 | 59064238 |
| -1.55 | 0.003 | AC093673.1 | chr7 | 143379692 | 143380495 |
| 1.54 | 0.005 | TNN | chr1 | 175067858 | 175148066 |
| 1.53 | 0.004 | AC022762.1 | chr11 | 6108135 | 6185576 |
| 1.52 | 0.039 | C9orf131 | chr9 | 35041095 | 35045991 |
| -1.52 | 0.000 | CHRNA9 | chr4 | 40335329 | 40355217 |
| 1.52 | 0.001 | AL132857.2 | chr14 | 36320753 | 36656163 |
| 1.51 | 0.005 | FGF2 | chr4 | 122826708 | 122898236 |
| 1.51 | 0.024 | SOD1P3 | chr8 | 125951861 | 125952314 |
| 1.50 | 0.006 | LINC00243 | chr6 | 30798654 | 30830659 |
| 1.49 | 0.011 | ANKRD20A19P | chr13 | 23939557 | 23946782 |
| 1.48 | 0.004 | DEPTOR | chr8 | 119873717 | 120050913 |
| 1.47 | 0.033 | LIPE-AS1 | chr19 | 42397128 | 42652355 |
| 1.47 | 0.002 | AL592494.1 | chr1 | 121494379 | 121510383 |
| 1.46 | 0.001 | PHTF1 | chr1 | 113696831 | 113759489 |
| -1.46 | 0.030 | AL034374.1 | chr6 | 53350158 | 53350705 |
| -1.46 | 0.000 | TRARG1 | chr17 | 1279663 | 1300987 |
| -1.46 | 0.001 | AL138899.2 | chr1 | 158195633 | 158196131 |
| -1.45 | 0.000 | HSPA8 | chr11 | 123057489 | 123063230 |
| 1.45 | 0.001 | INTS6-AS1 | chr13 | 51452367 | 51552364 |
| 1.45 | 0.006 | LINC01524 | chr20 | 52210645 | 52650431 |
| -1.44 | 0.001 | PGBD4P1 | chr7 | 142722358 | 142722764 |
| 1.43 | 0.000 | TUBB6 | chr18 | 12307669 | 12344320 |
| 1.43 | 0.035 | WDR97 | chr8 | 144107726 | 144118315 |
| 1.41 | 0.000 | MYO18B | chr22 | 25742144 | 26031041 |
| 1.40 | 0.004 | SLC4A9 | chr5 | 140360202 | 140375143 |
| 1.40 | 0.004 | AC125618.1 | chr3 | 180566718 | 180601935 |
| -1.40 | 0.003 | AC020661.1 | chr15 | 40906811 | 40910337 |
| -1.39 | 0.006 | GRAMD2A | chr15 | 72159807 | 72197785 |
| 1.38 | 0.045 | AL354740.1 | chr6 | 34248568 | 34286768 |
| 1.38 | 0.016 | HHLA2 | chr3 | 108296490 | 108378285 |
| 1.37 | 0.000 | KCNJ10 | chr1 | 159998651 | 160070483 |
| -1.36 | 0.000 | AL138899.1 | chr1 | 158197922 | 158203877 |
| 1.35 | 0.006 | AL354892.1 | chr6 | 169725504 | 169738992 |
| 1.35 | 0.046 | AP000253.1 | chr21 | 31653593 | 31659500 |
| 1.34 | 0.040 | DCC | chr18 | 52340172 | 53535903 |
| -1.33 | 0.013 | CRYBB1 | chr22 | 26599278 | 26618088 |
| 1.33 | 0.039 | FER1L5 | chr2 | 96642737 | 96704887 |
| 1.33 | 0.000 | COL4A4 | chr2 | 227002711 | 227164453 |
| -1.33 | 0.000 | DHFR2 | chr3 | 94047836 | 94063389 |
| 1.33 | 0.026 | CA15P1 | chr22 | 19031564 | 19034564 |
| 1.31 | 0.035 | SYNPO2 | chr4 | 118850688 | 119061247 |
| 1.31 | 0.015 | LINC00624 | chr1 | 147258885 | 147517875 |
| 1.31 | 0.034 | ENPP7P8 | chr11 | 71711740 | 71781183 |
| 1.31 | 0.013 | PDE6G | chr17 | 81650459 | 81663112 |
| 1.31 | 0.024 | AC011498.1 | chr19 | 4457962 | 4471493 |
| -1.31 | 0.000 | AC004585.1 | chr17 | 40517026 | 40527002 |
| 1.29 | 0.012 | DOK7 | chr4 | 3463311 | 3501473 |
| -1.29 | 0.011 | STYK1 | chr12 | 10618939 | 10674318 |
| 1.28 | 0.019 | EFNB2 | chr13 | 106489731 | 106535662 |
| 1.27 | 0.000 | SMIM15-AS1 | chr5 | 61162070 | 61232040 |
| 1.27 | 0.030 | C1S | chr12 | 6988259 | 7071032 |
| 1.27 | 0.022 | FUT6 | chr19 | 5830610 | 5839731 |
| 1.26 | 0.022 | AC244033.2 | chr1 | 150045660 | 150067701 |
| 1.25 | 0.000 | SNORC | chr2 | 232857270 | 232878708 |
| 1.25 | 0.019 | CLDN12 | chr7 | 90383721 | 90513402 |
| 1.24 | 0.019 | AC105383.1 | chr4 | 133075311 | 133149116 |
| 1.24 | 0.001 | AL589740.1 | chr6 | 97283303 | 98399872 |
| 1.23 | 0.040 | TNFSF13B | chr13 | 108251240 | 108308484 |
| 1.23 | 0.041 | AL109804.1 | chr20 | 3811384 | 3823882 |
| -1.23 | 0.000 | GIMAP7 | chr7 | 150514830 | 150521073 |
| 1.23 | 0.009 | MSANTD1 | chr4 | 3244369 | 3271738 |
| 1.23 | 0.036 | LINC00535 | chr8 | 93213302 | 93700433 |
| 1.23 | 0.001 | PPP2R3A | chr3 | 135965673 | 136147891 |
| 1.22 | 0.008 | NLRC4 | chr2 | 32224453 | 32265854 |
| 1.22 | 0.001 | RSPH9 | chr6 | 43645046 | 43672599 |
| 1.22 | 0.040 | RRH | chr4 | 109827994 | 109844604 |
| -1.21 | 0.000 | AC010422.6 | chr19 | 12525720 | 12580975 |
| 1.20 | 0.006 | SPTBN5 | chr15 | 41848144 | 41894077 |
| 1.20 | 0.001 | GRIK5 | chr19 | 41998321 | 42069498 |
| 1.19 | 0.013 | AC093423.3 | chr1 | 89633140 | 89933250 |
| -1.19 | 0.001 | RPL36AL | chr14 | 49618519 | 49620685 |
| -1.18 | 0.029 | GIMAP4 | chr7 | 150567277 | 150573955 |
| -1.18 | 0.038 | BAIAP2-DT | chr17 | 81029130 | 81034881 |
| -1.18 | 0.043 | TRBV28 | chr7 | 142720660 | 142721160 |
| 1.18 | 0.039 | AC007823.1 | chr3 | 179340322 | 179341887 |
| -1.18 | 0.000 | CD2 | chr1 | 116754385 | 116769228 |
| 1.17 | 0.005 | METTL21EP | chr13 | 102880099 | 102896033 |
| -1.17 | 0.001 | FAM30A | chr14 | 105917979 | 105932642 |
| -1.16 | 0.001 | CD40LG | chrX | 136648193 | 136660390 |
| 1.16 | 0.001 | AC008758.5 | chr19 | 12391949 | 12441082 |
| 1.16 | 0.007 | AC010542.3 | chr16 | 66469812 | 66517312 |
| -1.15 | 0.002 | GZMA | chr5 | 55102648 | 55110252 |
| 1.15 | 0.000 | GATM | chr15 | 45361124 | 45402327 |
| 1.15 | 0.000 | HAVCR2 | chr5 | 157085832 | 157142869 |
| 1.15 | 0.002 | AC015813.6 | chr17 | 58095672 | 58101479 |
| -1.15 | 0.000 | WDR6 | chr3 | 49007062 | 49015953 |
| 1.14 | 0.010 | ANKDD1B | chr5 | 75611459 | 75671846 |
| -1.14 | 0.003 | NPAS4 | chr11 | 66421004 | 66426707 |
| 1.13 | 0.003 | AC108471.2 | chr4 | 39639140 | 39666644 |
| 1.13 | 0.036 | AC009292.2 | chr15 | 67832725 | 67873866 |
| -1.13 | 0.004 | APOL1 | chr22 | 36253010 | 36267530 |
| -1.12 | 0.000 | SMIM33 | chr5 | 139470778 | 139474772 |
| 1.12 | 0.001 | TMEM236 | chr10 | 17752252 | 17800868 |
| 1.12 | 0.043 | DRAIC | chr15 | 69463026 | 69843120 |
| 1.12 | 0.001 | AC097103.2 | chr3 | 139389815 | 139583319 |
| -1.12 | 0.001 | TNFSF10 | chr3 | 172505508 | 172523507 |
| -1.11 | 0.000 | PTGER3 | chr1 | 70852353 | 71047808 |
| 1.11 | 0.001 | PKN2-AS1 | chr1 | 87620803 | 88685204 |
| 1.10 | 0.013 | NME9 | chr3 | 138261437 | 138329886 |
| -1.10 | 0.000 | GPR174 | chrX | 79144663 | 79175315 |
| -1.10 | 0.000 | SAMD9 | chr7 | 93099513 | 93118023 |
| 1.10 | 0.000 | AP3B2 | chr15 | 82659281 | 82709914 |
| 1.10 | 0.000 | C9orf43 | chr9 | 113410054 | 113429684 |
| 1.09 | 0.022 | KCNH8 | chr3 | 19148454 | 19535646 |
| 1.09 | 0.050 | CDH26 | chr20 | 59958427 | 60034011 |
| 1.09 | 0.000 | AL606489.1 | chr1 | 204377850 | 204435846 |
| -1.09 | 0.000 | CD1D | chr1 | 158179947 | 158184896 |
| 1.09 | 0.017 | AC093151.3 | chr1 | 41242373 | 41284861 |
| -1.09 | 0.048 | ACKR3 | chr2 | 236567787 | 236582358 |
| -1.08 | 0.018 | COA3 | chr17 | 42795147 | 42798704 |
| 1.07 | 0.003 | PARD6G-AS1 | chr18 | 80147924 | 80178432 |
| 1.07 | 0.001 | YPEL2 | chr17 | 59331689 | 59401729 |
| 1.07 | 0.006 | PRKG1 | chr10 | 50990891 | 52298350 |
| 1.07 | 0.018 | SGO1-AS1 | chr3 | 20174244 | 21145967 |
| 1.07 | 0.006 | C11orf65 | chr11 | 108308519 | 108467531 |
| 1.07 | 0.046 | ITGA10 | chr1 | 145891208 | 145910189 |
| 1.06 | 0.006 | AVIL | chr12 | 57797376 | 57818704 |
| 1.05 | 0.016 | DMGDH | chr5 | 78997606 | 79236038 |
| 1.04 | 0.004 | PIP5K1B | chr9 | 68705414 | 69009176 |
| 1.04 | 0.018 | AC090518.1 | chr15 | 56542952 | 56629592 |
| 1.04 | 0.000 | GVQW2 | chr6 | 138725211 | 138773652 |
| 1.04 | 0.001 | CYP3A5 | chr7 | 99648194 | 99680026 |
| 1.04 | 0.002 | SEPT7-AS1 | chr7 | 35751856 | 35800616 |
| 1.03 | 0.047 | OTOAP1 | chr16 | 22545698 | 22576865 |
| -1.03 | 0.007 | JCAD | chr10 | 30012800 | 30115494 |
| -1.03 | 0.006 | B3GALT6 | chr1 | 1232226 | 1235041 |
| 1.02 | 0.009 | AC109326.1 | chr17 | 43360041 | 43361361 |
| -1.02 | 0.027 | SPRY4 | chr5 | 142310427 | 142326455 |
| 1.02 | 0.003 | CRYM-AS1 | chr16 | 21300849 | 21339037 |
| 1.02 | 0.024 | ADCY10P1 | chr6 | 41101022 | 41140835 |
| 1.02 | 0.016 | MYH11 | chr16 | 15703172 | 15857033 |
| 1.01 | 0.047 | AC005532.1 | chr7 | 7255154 | 7277779 |
| -1.01 | 0.009 | CCR7 | chr17 | 40553769 | 40565472 |
| -1.01 | 0.019 | RNF144A-AS1 | chr2 | 6912277 | 6918709 |
| -1.01 | 0.022 | HNRNPH2 | chrX | 101408295 | 101414133 |
| 1.00 | 0.039 | Z84485.1 | chr6 | 36146698 | 36197205 |

**Supplementary Table 8: KEGG and GO pathway terms associated with significant genes that demonstrated decreased expression at the 4 hour time point.**

| Pathway | Description | n | DE | Adj.*p* |
| --- | --- | --- | --- | --- |
| GO:0002376 | immune system process | 55 | 1964 | 0.01 |
| GO:0006955 | immune response | 41 | 1298 | 0.01 |
| GO:0000315* | organellar large ribosomal subunit | 8 | 48 | 0.01 |
| GO:0005762* | mitochondrial large ribosomal subunit | 8 | 48 | 0.01 |
| GO:0032543* | mitochondrial translation | 11 | 113 | 0.01 |
| GO:0000313* | organellar ribosome | 9 | 69 | 0.01 |
| GO:0005761* | mitochondrial ribosome | 9 | 69 | 0.01 |
| GO:0070126 | mitochondrial translational termination | 9 | 73 | 0.01 |
| GO:0070125 | mitochondrial translational elongation | 9 | 72 | 0.01 |
| GO:1903039 | positive regulation of leukocyte cell-cell adhesion | 11 | 156 | 0.02 |
| GO:0048872 | homeostasis of number of cells | 12 | 195 | 0.02 |
| GO:0043624 | cellular protein complex disassembly | 12 | 183 | 0.02 |
| GO:0007159 | leukocyte cell-cell adhesion | 13 | 231 | 0.02 |
| GO:0140053* | mitochondrial gene expression | 11 | 138 | 0.03 |
| GO:0002521 | leukocyte differentiation | 16 | 347 | 0.03 |
| GO:0006414* | translational elongation | 10 | 111 | 0.03 |
| GO:0006415* | translational termination | 9 | 87 | 0.03 |
| GO:0022409 | positive regulation of cell-cell adhesion | 11 | 179 | 0.03 |
| GO:0007015 | actin filament organization | 13 | 303 | 0.04 |
| GO:0007155 | cell adhesion | 25 | 886 | 0.04 |
| GO:0022610 | biological adhesion | 25 | 888 | 0.04 |
| GO:0050863 | regulation of T cell activation | 12 | 219 | 0.05 |
| GO:0098609 | cell-cell adhesion | 18 | 525 | 0.05 |
| GO:0030155 | regulation of cell adhesion | 17 | 466 | 0.05 |
| GO:0001775 | cell activation | 29 | 948 | 0.05 |
| GO:0045321 | leukocyte activation | 27 | 847 | 0.05 |
| GO:0002684 | positive regulation of immune system process | 24 | 714 | 0.05 |
| KEGG:hsa05144 | Human Diseases; Infectious disease: parasitic Malaria - Homo sapiens (human) | 23 | 4 | 0.01 |
| KEGG:hsa04514 | Cell adhesion molecules - Homo sapiens (human) | 93 | 6 | 0.02 |

n, number of genes in the pathway

DE, number of genes that were differentially expressed

adjP, BH adjusted probability of differential expression

Light grey shading represent results from the GO database

* Pathways also differentially regulated at 24 hrs

**Supplementary Table 9. Top 100 most significant differentially expressed genes at the 24 hour time point, ordered by fold change.**

| Log_2_FC | adj.*p*.Val | Gene | CHR | start | end |
| --- | --- | --- | --- | --- | --- |
| 8.08 | < 0.0001 | AC007319.1 | chr2 | 187003220 | 187554663 |
| 6.66 | 0.0003 | TRIM61 | chr4 | 164954446 | 164977668 |
| 6.53 | < 0.0001 | AC096711.3 | chr4 | 132004810 | 132124031 |
| 6.49 | 0.0002 | AL160408.3 | chr1 | 234669523 | 234696153 |
| 6.30 | 0.0002 | MIR548XHG | chr21 | 18561265 | 18760003 |
| 5.99 | 0.0004 | ARMC3 | chr10 | 22928024 | 23038523 |
| 5.96 | 0.0016 | AL136537.2 | chr4 | 37001816 | 37020708 |
| 5.85 | 0.0047 | MIR137HG | chr1 | 97933474 | 98049863 |
| 5.66 | 0.0007 | AC244517.2 | chr5 | 141100242 | 141174391 |
| 5.60 | 0.0016 | LINC01630 | chr18 | 51346249 | 51643939 |
| 5.58 | 0.0015 | AC004009.1 | chr7 | 27361843 | 27409938 |
| 5.42 | 0.0062 | AC012355.1 | chr2 | 80699388 | 80870190 |
| 5.39 | 0.0016 | MEIS3 | chr19 | 47403124 | 47419523 |
| 5.39 | 0.0113 | MAGEB2 | chrX | 30215560 | 30220089 |
| 5.39 | 0.0010 | AADACL2-AS1 | chr3 | 151751443 | 151928175 |
| 5.36 | 0.0058 | AC022031.2 | chr18 | 55721063 | 55788761 |
| 5.35 | 0.0178 | RBMY1D | chrY | 21880076 | 21894526 |
| 5.34 | 0.0011 | ADAM29 | chr4 | 174829668 | 174978180 |
| 5.33 | 0.0013 | AC091806.1 | chrX | 40262917 | 40287720 |
| 5.28 | 0.0045 | LINC01241 | chr9 | 25780056 | 25812968 |
| 5.26 | 0.0021 | AC105053.1 | chr2 | 85815130 | 85825391 |
| 5.20 | 0.0026 | TGFBI | chr5 | 136028895 | 136063818 |
| 5.19 | 0.0017 | WNT11 | chr11 | 76186325 | 76210736 |
| 5.16 | < 0.0001 | MSR1 | chr8 | 16107878 | 16567490 |
| 5.14 | 0.0181 | KIAA1755 | chr20 | 38210488 | 38260772 |
| 5.13 | 0.0153 | C1orf158 | chr1 | 12746215 | 12763699 |
| 5.11 | 0.0161 | PPARG | chr3 | 12287368 | 12434356 |
| 5.11 | 0.0010 | XIST | chrX | 73820651 | 73852723 |
| 5.09 | 0.0003 | LIN7A | chr12 | 80792520 | 80937925 |
| 5.08 | 0.0093 | CPXM2 | chr10 | 123706207 | 123940267 |
| 5.06 | 0.0116 | SLC17A8 | chr12 | 100357079 | 100422059 |
| 5.05 | 0.0044 | AL161908.1 | chr9 | 126589594 | 126613700 |
| 5.05 | 0.0088 | NXPE2 | chr11 | 114678386 | 114706933 |
| 5.02 | 0.0006 | LINC01681 | chr1 | 170174403 | 170241287 |
| 5.02 | 0.0137 | AC026116.1 | chr12 | 63292625 | 63360037 |
| 4.99 | < 0.0001 | LGALS14 | chr19 | 39704306 | 39709444 |
| 4.99 | 0.0068 | NLRP2 | chr19 | 54953130 | 55001142 |
| 4.99 | 0.0040 | ZCCHC12 | chrX | 118823790 | 118826968 |
| 4.98 | 0.0028 | LINC01105 | chr2 | 5932687 | 6001275 |
| 4.97 | < 0.0001 | ZNF532 | chr18 | 58862600 | 58986480 |
| 4.95 | 0.0392 | ZFHX4 | chr8 | 76681219 | 76867285 |
| 4.93 | < 0.0001 | TDRG1 | chr6 | 40334954 | 40379887 |
| 4.91 | 0.0052 | AL844170.1 | chr1 | 84477039 | 84479157 |
| 4.89 | 0.0067 | AL713998.1 | chr6 | 67887646 | 67889339 |
| 4.89 | < 0.0001 | SSX2 | chrX | 52696896 | 52707189 |
| 4.87 | < 0.0001 | CCDC178 | chr18 | 32937402 | 33441101 |
| 4.83 | 0.0069 | LINC01228 | chr16 | 79798050 | 79827150 |
| 4.80 | 0.0130 | OR9Q1 | chr11 | 58023881 | 58181616 |
| 4.79 | 0.0047 | AC015574.1 | chr15 | 95638346 | 95825451 |
| 4.78 | 0.0309 | AC007342.4 | chr16 | 53373493 | 53384745 |
| 4.73 | 0.0230 | IL1R1 | chr2 | 102064544 | 102179874 |
| 4.72 | 0.0142 | LINC02422 | chr12 | 31877079 | 31887203 |
| 4.71 | < 0.0001 | TMEM196 | chr7 | 19719310 | 19773598 |
| 4.71 | 0.0261 | AC092783.1 | chr1 | 90510910 | 90533472 |
| 4.67 | 0.0244 | ARNT2 | chr15 | 80404350 | 80597937 |
| 4.66 | 0.0011 | AL591501.1 | chrX | 33726508 | 33942280 |
| 4.64 | 0.0123 | LIFR-AS1 | chr5 | 38556786 | 38671216 |
| 4.63 | < 0.0001 | SSX2B | chrX | 52751132 | 52790305 |
| 4.62 | 0.0089 | PDX1 | chr13 | 27920020 | 27926231 |
| 4.62 | < 0.0001 | COLEC12 | chr18 | 316740 | 500722 |
| 4.62 | 0.0414 | ELFN1 | chr7 | 1688119 | 1747954 |
| 4.60 | < 0.0001 | KLHDC7B | chr22 | 50545891 | 50551023 |
| 4.58 | 0.0169 | HOXD-AS2 | chr2 | 176121611 | 176137098 |
| 4.58 | < 0.0001 | NEBL | chr10 | 20779973 | 21174187 |
| 4.57 | 0.0181 | AL450992.1 | chr1 | 151994531 | 152042774 |
| 4.57 | 0.0224 | AC007491.1 | chr16 | 54628963 | 54657662 |
| 4.56 | 0.0113 | AC098934.4 | chr1 | 202810238 | 202810829 |
| 4.55 | 0.0445 | SPDYC | chr11 | 65170154 | 65173244 |
| 4.54 | 0.0159 | AL161733.1 | chr9 | 130933851 | 130945520 |
| 4.52 | 0.0091 | SCN4A | chr17 | 63938554 | 63972918 |
| 4.50 | 0.0194 | AC139491.2 | chr5 | 176143085 | 176185155 |
| 4.45 | 0.0186 | AC025575.1 | chr12 | 71034122 | 71104526 |
| 4.45 | 0.0194 | LINC01709 | chr1 | 101639548 | 101787572 |
| 4.40 | 0.0131 | LINC01684 | chr21 | 24428740 | 24547942 |
| 4.40 | < 0.0001 | AC123905.1 | chr12 | 71007773 | 71032083 |
| 4.40 | 0.0131 | AC008840.1 | chr5 | 95701249 | 95732295 |
| 4.40 | < 0.0001 | VAT1L | chr16 | 77788530 | 77980107 |
| 4.39 | 0.0010 | PAK3 | chrX | 110944285 | 111227361 |
| 4.39 | 0.0002 | TMEM92 | chr17 | 50271406 | 50281485 |
| 4.38 | 0.0235 | IGFN1 | chr1 | 201190825 | 201228952 |
| 4.38 | 0.0383 | LINC02140 | chr16 | 54366007 | 54370699 |
| 4.38 | 0.0108 | AL392023.2 | chr14 | 38034287 | 38194281 |
| 4.36 | 0.0249 | PLA2R1 | chr2 | 159932006 | 160062610 |
| 4.36 | 0.0295 | AL512638.1 | chr1 | 115471941 | 115476027 |
| 4.36 | 0.0325 | HAVCR1 | chr5 | 157029413 | 157059119 |
| 4.35 | 0.0032 | KCND2 | chr7 | 120273668 | 120750331 |
| 4.35 | 0.0305 | COL4A2 | chr13 | 110305812 | 110513209 |
| 4.31 | 0.0166 | AC091979.1 | chr5 | 158424585 | 158452758 |
| 4.30 | 0.0214 | SCN4B | chr11 | 118133377 | 118152888 |
| 4.27 | 0.0225 | LONRF3 | chrX | 118974614 | 119018355 |
| 4.27 | < 0.0001 | IL31RA | chr5 | 55851379 | 55922853 |
| 4.27 | 0.0300 | AC025887.2 | chr18 | 32470288 | 32745567 |
| 4.26 | < 0.0001 | TREHP1 | chr11 | 118688033 | 118690599 |
| 4.25 | 0.0034 | TMEM92-AS1 | chr17 | 50281577 | 50287855 |
| 4.22 | 0.0057 | DKFZp779M0652 | chr11 | 45771432 | 45772358 |
| 4.21 | 0.0328 | TEX26-AS1 | chr13 | 30881933 | 30933846 |
| 4.21 | < 0.0001 | DACH1 | chr13 | 71437966 | 71867192 |
| 4.18 | < 0.0001 | ANKRD34C-AS1 | chr15 | 79191707 | 79283945 |
| 4.16 | 0.0019 | AC100858.3 | chr8 | 124811226 | 124857516 |
| 4.16 | 0.0253 | AC019183.1 | chr18 | 1655177 | 1779955 |

**Supplementary Table 10. Top 100 most significant differentially expressed genes at the 48 hour time point, ordered by fold change.**

| Log_2_FC | adj.*p*.Val | Gene | CHR | start | end |
| --- | --- | --- | --- | --- | --- |
| 5.36 | 0.0228 | AC096711.3 | chr4 | 132004810 | 132124031 |
| 5.27 | 0.0452 | ZG16B | chr16 | 2830169 | 2839585 |
| 5.12 | 0.0332 | ACSL5 | chr10 | 112374018 | 112428380 |
| 4.65 | 0.0219 | CD5L | chr1 | 157830914 | 157898256 |
| -4.11 | 0.0452 | ADM | chr11 | 10304680 | 10307397 |
| 4.09 | 0.0027 | AC245100.3 | chr1 | 148288001 | 148288951 |
| -4.08 | 0.0467 | TNS3 | chr7 | 47275154 | 47582558 |
| 3.75 | <0.0001 | GLDN | chr15 | 51341629 | 51408013 |
| 3.02 | 0.0436 | SEMA3B | chr3 | 50267558 | 50277546 |
| 2.99 | 0.0125 | ZNF469 | chr16 | 88427471 | 88440757 |
| 2.96 | 0.0073 | IL31RA | chr5 | 55851379 | 55922853 |
| 2.88 | 0.0252 | RGS16 | chr1 | 182598623 | 182604408 |
| 2.74 | 0.0019 | NRIP1 | chr21 | 14961235 | 15065936 |
| -2.63 | 0.0373 | MIR210HG | chr11 | 565660 | 568457 |
| 2.60 | 0.0062 | TNF | chr6 | 31575567 | 31578336 |
| 2.55 | 0.0137 | CSTA | chr3 | 122325244 | 122341972 |
| -2.50 | 0.0065 | SLC32A1 | chr20 | 38724462 | 38729372 |
| 2.47 | 0.0006 | CARD6 | chr5 | 40841184 | 40860175 |
| 2.41 | 0.0028 | AC008696.2 | chr5 | 143605628 | 143828772 |
| 2.41 | <0.0001 | TEX41 | chr2 | 144667967 | 145262988 |
| -2.29 | 0.0109 | SLC2A1-AS1 | chr1 | 42959049 | 42983358 |
| 2.25 | 0.0341 | NEBL | chr10 | 20779973 | 21174187 |
| 2.22 | 0.0007 | AC109466.1 | chr5 | 164296696 | 165171643 |
| -2.21 | 0.0015 | PFKFB4 | chr3 | 48517684 | 48562015 |
| 2.18 | 0.0436 | AL691420.1 | chr9 | 115324932 | 115744330 |
| 2.18 | 0.0406 | LINC01828 | chr2 | 67086446 | 67311439 |
| 2.18 | 0.0117 | PARD3B | chr2 | 204545793 | 205620162 |
| 2.18 | 0.0221 | HIST1H2BA | chr6 | 25726777 | 25727292 |
| 2.17 | 0.0228 | UCP1 | chr4 | 140559434 | 140568805 |
| 2.17 | 0.0486 | FAM155A | chr13 | 107163510 | 107866735 |
| 2.16 | 0.0012 | RAB27B | chr18 | 54717860 | 54895516 |
| 2.14 | 0.0482 | MIR320A | chr8 | 22244962 | 22245043 |
| 2.14 | 0.0062 | DMBT1 | chr10 | 122560665 | 122643736 |
| 2.12 | 0.0004 | CPED1 | chr7 | 120988677 | 121297444 |
| 2.12 | <0.0001 | HIST3H3 | chr1 | 228424845 | 228425325 |
| -2.06 | 0.0025 | BNIP3 | chr10 | 131966455 | 131982013 |
| 2.04 | 0.0207 | FGF22 | chr19 | 639879 | 644371 |
| 2.04 | 0.0231 | LINC02105 | chr5 | 53776644 | 53819686 |
| 2.03 | <0.0001 | KRT1 | chr12 | 52674736 | 52680407 |
| 2.03 | 0.0449 | S100P | chr4 | 6693069 | 6697170 |
| -2.02 | 0.0314 | AL137145.2 | chr10 | 6277687 | 6335982 |
| 2.00 | 0.0065 | ARC | chr8 | 142611044 | 142614472 |
| 1.96 | <0.0001 | STXBP6 | chr14 | 24809656 | 25050297 |
| 1.94 | 0.0467 | SERPINE1 | chr7 | 101127089 | 101139266 |
| 1.93 | 0.0015 | TFEC | chr7 | 115935148 | 116159896 |
| 1.92 | 0.0452 | SPOCK1 | chr5 | 136975298 | 137598379 |
| 1.91 | 0.0004 | CD93 | chr20 | 23079349 | 23086340 |
| 1.86 | 0.0004 | SYNPO2 | chr4 | 118850688 | 119061247 |
| 1.85 | 0.0094 | AC016152.1 | chr12 | 98113014 | 98292445 |
| 1.82 | 0.0216 | CACNA1E | chr1 | 181317690 | 181808084 |
| -1.81 | 0.0018 | CFAP97D2 | chr13 | 114154691 | 114223084 |
| 1.81 | <0.0001 | CTSW | chr11 | 65879809 | 65883741 |
| 1.80 | 0.0257 | NRP1 | chr10 | 33177492 | 33336262 |
| 1.73 | <0.0001 | FBXO39 | chr17 | 6776223 | 6797101 |
| -1.70 | 0.0452 | KIT | chr4 | 54657918 | 54740715 |
| 1.69 | 0.0391 | CHODL | chr21 | 17901263 | 18267373 |
| 1.68 | 0.0332 | LHFPL6 | chr13 | 39209116 | 39603528 |
| 1.68 | 0.0101 | AC004828.1 | chr14 | 72552580 | 72595125 |
| 1.66 | 0.0027 | HIST1H1T | chr6 | 26107419 | 26108136 |
| 1.66 | 0.0052 | FAM19A1 | chr3 | 68004216 | 68545625 |
| 1.66 | 0.0118 | MIR4527HG | chr18 | 47285725 | 47594550 |
| 1.65 | 0.0332 | CTSS | chr1 | 150730196 | 150765957 |
| 1.64 | 0.0446 | CREB5 | chr7 | 28299321 | 28825894 |
| 1.61 | 0.0373 | AC092813.1 | chr1 | 75932479 | 76019356 |
| 1.61 | 0.0004 | ZBTB16 | chr11 | 114059593 | 114250676 |
| -1.61 | <0.0001 | RAG1 | chr11 | 36510709 | 36593156 |
| 1.59 | 0.0063 | AC008957.1 | chr5 | 36666214 | 36725195 |
| 1.58 | 0.0320 | Z92544.2 | chr16 | 689001 | 692554 |
| 1.56 | 0.0332 | FRK | chr6 | 115931149 | 116060758 |
| 1.56 | 0.0042 | CDKN1A | chr6 | 36676460 | 36687339 |
| -1.55 | 0.0181 | DARS-AS1 | chr2 | 135985176 | 136022593 |
| 1.53 | 0.0415 | CTSO | chr4 | 155924118 | 155953917 |
| -1.50 | 0.0109 | KCNJ4 | chr22 | 38426327 | 38455199 |
| -1.49 | 0.0009 | FAM110B | chr8 | 57994509 | 58204279 |
| 1.48 | 0.0120 | SGCD | chr5 | 155870344 | 156767788 |
| 1.46 | <0001 | MTUS2 | chr13 | 28820348 | 29505947 |
| 1.45 | <0001 | AP002518.2 | chr11 | 114210616 | 114356571 |
| 1.44 | 0.0118 | KCNK17 | chr6 | 39299001 | 39314553 |
| -1.44 | 0.0133 | ADGRG1 | chr16 | 57610652 | 57665580 |
| 1.44 | 0.0065 | TRARG1 | chr17 | 1279663 | 1300987 |
| 1.42 | 0.0452 | AC010168.1 | chr12 | 14665655 | 14757963 |
| 1.41 | 0.0036 | CFAP47 | chrX | 35919734 | 36385319 |
| 1.38 | 0.0073 | HBEGF | chr5 | 140332843 | 140346631 |
| 1.37 | <0001 | PLCH1 | chr3 | 155375580 | 155745067 |
| 1.36 | 0.0494 | TRGV8 | chr7 | 38330343 | 38330935 |
| 1.36 | 0.0004 | GZMA | chr5 | 55102648 | 55110252 |
| -1.35 | 0.0004 | CTTN | chr11 | 70398404 | 70436584 |
| -1.33 | 0.0049 | LGMN | chr14 | 92703807 | 92748702 |
| -1.32 | 0.0047 | GPR162 | chr12 | 6821545 | 6829972 |
| 1.32 | 0.0021 | NELL1 | chr11 | 20669551 | 21575681 |
| -1.31 | 0.0265 | TRIB1 | chr8 | 125430321 | 125438405 |
| -1.30 | 0.0117 | GABRG3 | chr15 | 26971282 | 27541991 |
| -1.29 | 0.0027 | NOTCH3 | chr19 | 15159038 | 15200981 |
| 1.26 | 0.0117 | S100A4 | chr1 | 153543613 | 153550136 |
| 1.26 | 0.0288 | GPR158 | chr10 | 25174802 | 25602226 |
| 1.25 | 0.0004 | GPA33 | chr1 | 167052836 | 167166479 |
| -1.24 | 0.0483 | CD4 | chr12 | 6786858 | 6820808 |
| -1.21 | 0.0185 | MAP1A | chr15 | 43510958 | 43531620 |
| 1.20 | 0.0406 | DPP6 | chr7 | 153887097 | 154894285 |
| -1.20 | 0.0063 | MAL | chr2 | 95025677 | 95053996 |

**Supplementary Table 11. List of probes that demonstrated a significant correlation between DNA methylation and gene expression at 4 hours.**

| corr_R | t | adjP | Gene_name |
| --- | --- | --- | --- |
| 1.00 | 20.95 | 0.02 | COA3 |
| -0.99 | -15.08 | 0.03 | C3orf20 |
| 0.99 | 13.54 | 0.03 | MRPL43 |
| -0.99 | -11.52 | 0.05 | DLEU7 |
| 0.98 | 10.24 | 0.05 | LIMA1 |
| -0.98 | -8.98 | 0.05 | MSANTD1 |
| 0.98 | 8.86 | 0.05 | SNN |
| -0.98 | -8.84 | 0.05 | PDE6G |
| 0.97 | 8.59 | 0.05 | PRKG1 |
| -0.97 | -8.26 | 0.05 | AP3B2 |
| 0.97 | 8.18 | 0.05 | SYBU |
| -0.97 | -8.12 | 0.05 | PDE6G |
| 0.97 | 8.09 | 0.05 | EFNB2 |
| -0.97 | -7.96 | 0.05 | PM20D1 |
| -0.97 | -7.96 | 0.05 | DNHD1 |
| -0.97 | -7.82 | 0.05 | HIST3H3 |
| -0.97 | -7.65 | 0.05 | PTGER3 |
| -0.97 | -7.62 | 0.05 | BCCIP |
| 0.97 | 7.60 | 0.05 | LIMA1 |
| -0.97 | -7.58 | 0.05 | SPIRE2 |
| -0.96 | -7.22 | 0.05 | CD93 |
| -0.96 | -7.14 | 0.05 | MRPL43 |
| -0.96 | -7.13 | 0.05 | SMC5-AS1 |
| 0.96 | 7.11 | 0.05 | PARP9 |
| -0.96 | -7.00 | 0.05 | PM20D1 |
| -0.96 | -6.63 | 0.05 | C5orf24 |
| 0.96 | 6.62 | 0.05 | SYBU |
| -0.96 | -6.47 | 0.05 | PARD6G-AS1 |
| -0.96 | -6.46 | 0.05 | ACP4 |
| 0.96 | 6.44 | 0.05 | RB1 |
| 0.95 | 6.36 | 0.05 | LYRM2 |
| 0.95 | 6.36 | 0.05 | LYRM2 |
| -0.95 | -6.26 | 0.05 | HORMAD2-AS1 |
| -0.95 | -6.20 | 0.05 | DHFR2 |
| 0.95 | 6.20 | 0.05 | DRAIC |
| -0.95 | -6.15 | 0.05 | MYH11 |
| 0.95 | 6.08 | 0.05 | RAC2 |
| -0.95 | -6.06 | 0.05 | DNHD1 |
| -0.95 | -6.03 | 0.05 | AVIL |
| 0.95 | 6.03 | 0.05 | AGAP2 |
| -0.95 | -5.99 | 0.05 | AP3B2 |
| -0.95 | -5.97 | 0.05 | TMX1 |
| -0.95 | -5.95 | 0.05 | PTGER3 |
| 0.95 | 5.91 | 0.05 | AGAP2 |
| -0.95 | -5.84 | 0.05 | PHTF1 |
| 0.94 | 5.72 | 0.05 | MED1 |
| -0.94 | -5.65 | 0.05 | CREBRF |
| 0.94 | 5.58 | 0.05 | SH3GL3 |
| 0.94 | 5.57 | 0.05 | KNL1 |
| -0.94 | -5.57 | 0.05 | RIPK4 |
| 0.94 | 5.55 | 0.05 | STXBP6 |
| -0.94 | -5.49 | 0.05 | MSANTD1 |
| 0.94 | 5.48 | 0.05 | IKZF1 |
| -0.94 | -5.42 | 0.05 | RSPH9 |
| -0.94 | -5.36 | 0.05 | SPNS2 |
| 0.94 | 5.35 | 0.05 | DCC |
| 0.94 | 5.32 | 0.05 | CABLES2 |
| 0.94 | 5.32 | 0.05 | SYBU |
| -0.94 | -5.30 | 0.05 | PSD4 |
| -0.94 | -5.29 | 0.05 | TEX21P |
| -0.93 | -5.27 | 0.05 | COL20A1 |
| 0.93 | 5.25 | 0.05 | NRM |
| -0.93 | -5.25 | 0.05 | AL139011.2 |
| 0.93 | 5.24 | 0.05 | GVQW2 |
| 0.93 | 5.24 | 0.05 | LINC00535 |
| -0.93 | -5.23 | 0.05 | AL109804.1 |
| -0.93 | -5.22 | 0.05 | MYL12A |
| 0.93 | 5.19 | 0.05 | USP16 |
| -0.93 | -5.18 | 0.05 | ALPK3 |
| 0.93 | 5.17 | 0.05 | LINC01301 |
| 0.93 | 5.13 | 0.05 | PRKG1 |
| -0.93 | -5.11 | 0.05 | RSPH9 |
| -0.93 | -5.10 | 0.05 | AC009053.3 |
| 0.93 | 5.09 | 0.05 | KCND3 |
| -0.93 | -5.08 | 0.05 | FOXR1 |
| -0.93 | -5.08 | 0.05 | C5orf22 |
| 0.93 | 5.06 | 0.05 | RAC2 |
| -0.93 | -5.05 | 0.05 | ADCY10P1 |
| -0.93 | -5.04 | 0.05 | RSPH9 |
| -0.93 | -5.02 | 0.05 | HORMAD2-AS1 |
| -0.93 | -5.01 | 0.05 | CICP14 |
| -0.93 | -4.98 | 0.05 | HORMAD2-AS1 |
| 0.93 | 4.98 | 0.05 | COA3 |
| 0.93 | 4.97 | 0.05 | TTC23 |
| 0.93 | 4.96 | 0.05 | DTX3L |
| -0.93 | -4.93 | 0.05 | RSPH9 |
| 0.93 | 4.92 | 0.05 | PRKG1 |
| 0.93 | 4.91 | 0.05 | GCNT4 |
| 0.93 | 4.90 | 0.05 | CABLES2 |
| -0.93 | -4.90 | 0.05 | NECAB3 |
| -0.92 | -4.83 | 0.05 | PRAME |
| -0.92 | -4.81 | 0.05 | SPTBN5 |
| 0.92 | 4.81 | 0.05 | AC012574.1 |
| -0.92 | -4.77 | 0.05 | CD93 |
| 0.92 | 4.76 | 0.05 | AC008758.5 |
| -0.92 | -4.75 | 0.05 | COL20A1 |
| 0.92 | 4.70 | 0.05 | TPK1 |
| -0.92 | -4.69 | 0.05 | IQCN |
| -0.92 | -4.69 | 0.05 | FASTKD2 |
| -0.92 | -4.67 | 0.05 | PM20D1 |
| -0.92 | -4.66 | 0.05 | SAXO1 |
| 0.92 | 4.62 | 0.05 | KCND3 |
| -0.92 | -4.61 | 0.05 | AC053527.1 |
| -0.92 | -4.60 | 0.05 | HIST3H3 |
| -0.92 | -4.57 | 0.05 | C9orf43 |
| -0.92 | -4.57 | 0.05 | DNHD1 |
| -0.92 | -4.55 | 0.05 | CPSF6 |
| 0.91 | 4.53 | 0.05 | CNNM2 |
| 0.91 | 4.52 | 0.05 | S1PR1 |
| -0.91 | -4.51 | 0.05 | ADCY10P1 |
| -0.91 | -4.49 | 0.05 | ITGA10 |
| -0.91 | -4.46 | 0.06 | ITGA10 |

* Table ordered by correlation coefficient (methylation vs gene expression)

**Supplementary Table 12. Top 100 probes that demonstrated a significant correlation between DNA methylation and gene expression at 24 hours.**

| corr_R | t | adjP | Gene_name |
| --- | --- | --- | --- |
| 1.00 | 32.80 | 0.01 | CTNNA3 |
| 1.00 | 29.23 | 0.01 | ANXA4 |
| 1.00 | 28.07 | 0.01 | ZNF101 |
| 1.00 | 25.52 | 0.01 | NRP1 |
| 1.00 | 24.22 | 0.01 | IMPDH2 |
| 1.00 | 23.90 | 0.01 | EPHA3 |
| 1.00 | 22.02 | 0.02 | ZNF232 |
| 1.00 | 20.62 | 0.02 | SNRNP25 |
| 1.00 | 20.15 | 0.02 | NMNAT3 |
| 0.99 | 19.63 | 0.02 | AC034111.1 |
| -0.99 | -19.21 | 0.02 | ANKDD1A |
| -0.99 | -18.64 | 0.02 | AC073517.1 |
| 0.99 | 17.93 | 0.02 | GTPBP3 |
| 0.99 | 17.49 | 0.02 | ARHGAP40 |
| 0.99 | 17.42 | 0.02 | AL359232.1 |
| -0.99 | -16.82 | 0.02 | SEC31B |
| 0.99 | 16.47 | 0.02 | EPHA3 |
| 0.99 | 16.18 | 0.02 | THOP1 |
| 0.99 | 16.12 | 0.02 | CCDC178 |
| 0.99 | 15.70 | 0.02 | LYPLAL1-DT |
| 0.99 | 15.69 | 0.02 | GAB2 |
| 0.99 | 15.52 | 0.02 | DNAJC11 |
| 0.99 | 15.50 | 0.02 | CD93 |
| 0.99 | 15.38 | 0.02 | PTPRT |
| 0.99 | 15.22 | 0.02 | PLUT |
| 0.99 | 15.13 | 0.02 | RGS16 |
| 0.99 | 15.01 | 0.02 | FLVCR1 |
| -0.99 | -14.86 | 0.02 | BCLAF3 |
| 0.99 | 14.76 | 0.02 | MEIS1 |
| -0.99 | -14.71 | 0.02 | SEC31B |
| -0.99 | -14.52 | 0.02 | LINC02591 |
| 0.99 | 14.36 | 0.02 | MAP7 |
| -0.99 | -14.14 | 0.02 | HOXA13 |
| 0.99 | 14.14 | 0.02 | KCND2 |
| 0.99 | 13.81 | 0.02 | TRIP13 |
| 0.99 | 13.66 | 0.02 | WNK2 |
| -0.99 | -13.59 | 0.02 | ADGRA3 |
| -0.99 | -13.58 | 0.02 | MYH11 |
| -0.99 | -13.58 | 0.02 | ITGB1-DT |
| 0.99 | 13.48 | 0.02 | GRB10 |
| 0.99 | 13.32 | 0.02 | RPL23 |
| 0.99 | 13.30 | 0.02 | NQO1 |
| 0.99 | 13.27 | 0.02 | IL16 |
| -0.99 | -13.25 | 0.02 | CTNND1 |
| 0.99 | 13.22 | 0.02 | ATP5MPL |
| 0.99 | 12.98 | 0.02 | ADGB |
| 0.99 | 12.93 | 0.02 | HAGHL |
| -0.99 | -12.89 | 0.02 | RTCB |
| -0.99 | -12.72 | 0.02 | ACER1 |
| -0.99 | -12.69 | 0.02 | TMPRSS13 |
| 0.99 | 12.68 | 0.02 | ZNF226 |
| -0.99 | -12.54 | 0.02 | AC022126.1 |
| -0.99 | -12.30 | 0.02 | CMPK1 |
| -0.99 | -12.15 | 0.02 | CTSG |
| 0.99 | 12.05 | 0.02 | KIAA1217 |
| 0.99 | 11.93 | 0.02 | LIMA1 |
| 0.99 | 11.73 | 0.02 | AP005482.1 |
| 0.99 | 11.72 | 0.02 | AGBL4 |
| 0.99 | 11.64 | 0.02 | SLIT3 |
| 0.99 | 11.62 | 0.02 | CELSR3 |
| -0.99 | -11.56 | 0.02 | PDE3A |
| 0.99 | 11.42 | 0.03 | MEIS1 |
| -0.98 | -11.37 | 0.03 | ACER1 |
| 0.98 | 11.32 | 0.03 | OPALIN |
| 0.98 | 11.29 | 0.03 | LEXM |
| 0.98 | 11.21 | 0.03 | EIF3A |
| 0.98 | 11.15 | 0.03 | CREB5 |
| 0.98 | 11.14 | 0.03 | RNH1 |
| -0.98 | -11.07 | 0.03 | ACER1 |
| -0.98 | -11.00 | 0.03 | SEC31B |
| -0.98 | -10.97 | 0.03 | SCUBE2 |
| 0.98 | 10.95 | 0.03 | NHP2 |
| -0.98 | -10.92 | 0.03 | SLC17A5 |
| 0.98 | 10.90 | 0.03 | ABCA4 |
| -0.98 | -10.87 | 0.03 | C3orf20 |
| 0.98 | 10.87 | 0.03 | WDR46 |
| 0.98 | 10.83 | 0.03 | BX255925.3 |
| 0.98 | 10.75 | 0.03 | CELF4 |
| -0.98 | -10.65 | 0.03 | GCNA |
| 0.98 | 10.52 | 0.03 | ASF1B |
| -0.98 | -10.51 | 0.03 | SEC31B |
| 0.98 | 10.49 | 0.03 | ZNF142 |
| 0.98 | 10.46 | 0.03 | LINC00958 |
| 0.98 | 10.42 | 0.03 | AGBL4 |
| 0.98 | 10.34 | 0.03 | TSPAN31 |
| 0.98 | 10.26 | 0.03 | HCN4 |
| 0.98 | 10.20 | 0.03 | RRP1B |
| 0.98 | 10.19 | 0.03 | AAAS |
| 0.98 | 10.14 | 0.03 | NHP2 |
| 0.98 | 10.13 | 0.03 | TBCD |
| -0.98 | -10.08 | 0.03 | CTNND1 |
| 0.98 | 9.93 | 0.03 | FRK |
| 0.98 | 9.84 | 0.03 | TOP3A |
| -0.98 | -9.84 | 0.03 | ONECUT2 |
| -0.98 | -9.79 | 0.03 | SNRPA1 |
| 0.98 | 9.79 | 0.03 | KIF1C |
| -0.98 | -9.78 | 0.03 | ANKDD1A |
| -0.98 | -9.74 | 0.03 | ARHGAP23 |
| 0.98 | 9.74 | 0.03 | PSMB3 |

* Table ordered by correlation coefficient (methylation vs gene expression)

**Supplementary Table 13: KEGG and GO pathway analysis results containing top hits from the differentially methylated CpGs from the 4-hour time point with a FDR of <0.05**

| Pathway | Description | n | DM | FDR |
| --- | --- | --- | --- | --- |
| path:hsa04910 | Insulin signalling pathway | 137 | 107 | < 0.001 |
| path:hsa04144 | Endocytosis | 249 | 178 | < 0.001 |
| path:hsa05170 | Human immunodeficiency virus 1 infection | 212 | 146 | < 0.001 |
| path:hsa04370 | VEGF signalling pathway | 59 | 51 | 0.001 |
| path:hsa04722 | Neurotrophin signalling pathway | 119 | 92 | 0.001 |
| path:hsa04660 | T cell receptor signalling pathway | 104 | 80 | 0.001 |
| path:hsa05131 | Shigellosis | 236 | 161 | 0.002 |
| path:hsa04666 | Fc gamma R-mediated phagocytosis | 93 | 72 | 0.002 |
| path:hsa05235 | PD-L1 expression and PD-1 checkpoint pathway in cancer | 89 | 68 | 0.002 |
| path:hsa01100 | Metabolic pathways | 1489 | 875 | 0.002 |
| path:hsa04150 | mTOR signalling pathway | 153 | 111 | 0.003 |
| path:hsa04071 | Sphingolipid signalling pathway | 119 | 88 | 0.003 |
| path:hsa04664 | Fc epsilon RI signalling pathway | 68 | 54 | 0.003 |
| path:hsa04210 | Apoptosis | 136 | 97 | 0.003 |
| path:hsa04010 | MAPK signalling pathway | 295 | 199 | 0.005 |
| path:hsa05215 | Prostate cancer | 97 | 73 | 0.005 |
| path:hsa05231 | Choline metabolism in cancer | 98 | 75 | 0.005 |
| path:hsa05221 | Acute myeloid leukemia | 67 | 53 | 0.007 |
| path:hsa04925 | Aldosterone synthesis and secretion | 98 | 72 | 0.012 |
| path:hsa04070 | Phosphatidylinositol signalling system | 99 | 74 | 0.014 |
| path:hsa04931 | Insulin resistance | 108 | 78 | 0.019 |
| path:hsa04072 | Phospholipase D signalling pathway | 148 | 104 | 0.019 |
| path:hsa05132 | Salmonella infection | 214 | 143 | 0.021 |
| path:hsa05225 | Hepatocellular carcinoma | 168 | 115 | 0.022 |
| path:hsa01230 | Biosynthesis of amino acids | 75 | 53 | 0.023 |
| path:hsa00564 | Glycerophospholipid metabolism | 98 | 70 | 0.027 |
| path:hsa04062 | Chemokine signalling pathway | 189 | 119 | 0.034 |
| path:hsa04152 | AMPK signalling pathway | 120 | 84 | 0.034 |
| path:hsa04218 | Cellular senescence | 160 | 108 | 0.038 |
| path:hsa04912 | GnRH signalling pathway | 93 | 66 | 0.04 |
| path:hsa04744 | Phototransduction | 28 | 22 | 0.04 |
| path:hsa04810 | Regulation of actin cytoskeleton | 213 | 141 | 0.04 |
| path:hsa05214 | Glioma | 75 | 55 | 0.04 |
| path:hsa05220 | Chronic myeloid leukemia | 76 | 56 | 0.04 |
| path:hsa04919 | Thyroid hormone signalling pathway | 119 | 84 | 0.04 |
| path:hsa01521 | EGFR tyrosine kinase inhibitor resistance | 79 | 59 | 0.04 |
| path:hsa04935 | Growth hormone synthesis, secretion and action | 119 | 84 | 0.04 |
| path:hsa05135 | Yersinia infection | 120 | 84 | 0.04 |
| path:hsa05211 | Renal cell carcinoma | 69 | 52 | 0.04 |
| path:hsa04625 | C-type lectin receptor signalling pathway | 104 | 72 | 0.04 |
| path:hsa05213 | Endometrial cancer | 58 | 44 | 0.04 |
| path:hsa05163 | Human cytomegalovirus infection | 225 | 143 | 0.041 |
| path:hsa00330 | Arginine and proline metabolism | 50 | 36 | 0.045 |
| path:hsa04964 | Proximal tubule bicarbonate reclamation | 23 | 19 | 0.047 |
| path:hsa04142 | Lysosome | 123 | 83 | 0.047 |
| path:hsa00562 | Inositol phosphate metabolism | 74 | 55 | 0.047 |
| path:hsa04140 | Autophagy - animal | 137 | 94 | 0.047 |
| path:hsa04929 | GnRH secretion | 64 | 48 | 0.047 |
| path:hsa05166 | Human T-cell leukemia virus 1 infection | 219 | 142 | 0.049 |
| path:hsa04630 | JAK-STAT signalling pathway | 162 | 99 | 0.049 |
| path:hsa04714 | Thermogenesis | 231 | 139 | 0.049 |
| path:hsa04662 | B cell receptor signalling pathway | 82 | 57 | 0.05 |
| GO:0005524 | ATP binding | 1461 | 376 | 0.001 |
| GO:0030554 | adenyl nucleotide binding | 1529 | 390 | 0.001 |
| GO:0032559 | adenyl ribonucleotide binding | 1518 | 388 | 0.001 |
| GO:0000166 | nucleotide binding | 2100 | 497 | 0.001 |
| GO:1901265 | nucleoside phosphate binding | 2101 | 497 | 0.001 |
| GO:0035639 | purine ribonucleoside triphosphate binding | 1793 | 432 | 0.001 |
| GO:0032555 | purine ribonucleotide binding | 1857 | 445 | 0.001 |
| GO:0017076 | purine nucleotide binding | 1870 | 447 | 0.001 |
| GO:0032553 | ribonucleotide binding | 1873 | 446 | 0.003 |
| GO:0036094 | small molecule binding | 2475 | 558 | 0.004 |
| GO:0008144 | drug binding | 1709 | 410 | 0.010 |
| GO:0043168 | anion binding | 2747 | 618 | 0.016 |

n, number of genes in the pathway

DM, number of genes that were differentially methylated

FDR, false discovery rate

Light grey shading represent results from the GO database

**Supplementary Table 14. GO pathway terms associated with genes that demonstrated a significant negative correlation between DNA methylation at the 4 hour time point and gene expression at the 24 hour time point.**

| Pathway | Description | n | DE | ontology | *p*value |
| --- | --- | --- | --- | --- | --- |
| GO:0007017 | microtubule-based process | 4 | 14 | BP | 0.006 |
| GO:0012506 | vesicle membrane | 3 | 8 | CC | 0.008 |
| GO:0030659 | cytoplasmic vesicle membrane | 3 | 8 | CC | 0.008 |
| GO:0006281 | DNA repair | 3 | 8 | BP | 0.009 |
| GO:0051656 | establishment of organelle localization | 3 | 11 | BP | 0.022 |
| GO:0031982 | vesicle | 6 | 44 | CC | 0.028 |
| GO:0006974 | cellular response to DNA damage stimulus | 3 | 12 | BP | 0.029 |
| GO:0051640 | organelle localization | 3 | 14 | BP | 0.043 |
| GO:0044433 | cytoplasmic vesicle part | 3 | 14 | CC | 0.044 |
| GO:0006302 | double-strand break repair | 2 | 6 | BP | 0.046 |
| GO:0000729 | DNA double-strand break processing | 4 | 6 | BP | 0.001 |
| GO:0048037 | cofactor binding | 13 | 55 | MF | 0.003 |
| GO:0004112 | cyclic-nucleotide phosphodiesterase activity | 3 | 6 | MF | 0.008 |
| GO:0004114 | 3',5'-cyclic-nucleotide phosphodiesterase activity | 3 | 6 | MF | 0.008 |
| GO:0071345 | cellular response to cytokine stimulus | 21 | 122 | BP | 0.009 |
| GO:0030532 | small nuclear ribonucleoprotein complex | 6 | 19 | CC | 0.010 |
| GO:0097525 | spliceosomal snRNP complex | 6 | 19 | CC | 0.010 |
| GO:0008081 | phosphoric diester hydrolase activity | 4 | 11 | MF | 0.011 |
| GO:0000795 | synaptonemal complex | 4 | 10 | CC | 0.012 |
| GO:0099086 | synaptonemal structure | 4 | 10 | CC | 0.012 |
| GO:0120114 | Sm-like protein family complex | 6 | 21 | CC | 0.017 |
| GO:0022408 | negative regulation of cell-cell adhesion | 6 | 22 | BP | 0.018 |
| GO:1990823 | response to leukemia inhibitory factor | 5 | 16 | BP | 0.019 |
| GO:1990830 | cellular response to leukemia inhibitory factor | 5 | 16 | BP | 0.019 |
| GO:0034728 | nucleosome organization | 11 | 49 | BP | 0.020 |
| GO:0005615 | extracellular space | 51 | 390 | CC | 0.020 |
| GO:0046928 | regulation of neurotransmitter secretion | 4 | 13 | BP | 0.020 |
| GO:0046906 | tetrapyrrole binding | 5 | 17 | MF | 0.022 |
| GO:0006333 | chromatin assembly or disassembly | 11 | 50 | BP | 0.022 |
| GO:0045637 | regulation of myeloid cell differentiation | 8 | 33 | BP | 0.023 |
| GO:0034097 | response to cytokine | 21 | 135 | BP | 0.028 |
| GO:0005685 | U1 snRNP | 3 | 7 | CC | 0.028 |
| GO:0031060 | regulation of histone methylation | 3 | 7 | BP | 0.029 |
| GO:0031062 | positive regulation of histone methylation | 3 | 7 | BP | 0.029 |
| GO:0031256 | leading edge membrane | 4 | 14 | CC | 0.030 |
| GO:0006334 | nucleosome assembly | 10 | 46 | BP | 0.032 |
| GO:0072523 | purine-containing compound catabolic process | 3 | 8 | BP | 0.033 |
| GO:0071005 | U2-type precatalytic spliceosome | 5 | 18 | CC | 0.034 |
| GO:0071011 | precatalytic spliceosome | 5 | 18 | CC | 0.034 |
| GO:0038111 | interleukin-7-mediated signaling pathway | 3 | 7 | BP | 0.034 |
| GO:0050866 | negative regulation of cell activation | 5 | 19 | BP | 0.036 |
| GO:1903035 | negative regulation of response to wounding | 4 | 14 | BP | 0.038 |
| GO:0050662 | coenzyme binding | 6 | 25 | MF | 0.038 |
| GO:0072376 | protein activation cascade | 3 | 8 | BP | 0.039 |
| GO:0000786 | Nucleosome | 9 | 40 | CC | 0.040 |
| GO:0020037 | heme binding | 4 | 14 | MF | 0.042 |
| GO:0031497 | chromatin assembly | 10 | 48 | BP | 0.042 |
| GO:0002526 | acute inflammatory response | 3 | 8 | BP | 0.042 |
| GO:0051287 | NAD binding | 3 | 8 | MF | 0.043 |
| GO:0071007 | U2-type catalytic step 2 spliceosome | 3 | 8 | CC | 0.043 |
| GO:0070062 | extracellular exosome | 39 | 294 | CC | 0.044 |
| GO:0032945 | negative regulation of mononuclear cell proliferation | 3 | 8 | BP | 0.045 |
| GO:0042130 | negative regulation of T cell proliferation | 3 | 8 | BP | 0.045 |
| GO:0050672 | negative regulation of lymphocyte proliferation | 3 | 8 | BP | 0.045 |
| GO:0070664 | negative regulation of leukocyte proliferation | 3 | 8 | BP | 0.045 |
| GO:0048786 | presynaptic active zone | 3 | 10 | CC | 0.045 |
| GO:0051290 | protein heterotetramerization | 4 | 12 | BP | 0.046 |
| GO:0045652 | regulation of megakaryocyte differentiation | 5 | 18 | BP | 0.046 |
| GO:0000183 | chromatin silencing at rDNA | 4 | 12 | BP | 0.046 |
| GO:0051588 | regulation of neurotransmitter transport | 4 | 16 | BP | 0.047 |
| GO:0098760 | response to interleukin-7 | 3 | 8 | BP | 0.049 |
| GO:0098761 | cellular response to interleukin-7 | 3 | 8 | BP | 0.049 |
| GO:0051291 | protein heterooligomerization | 5 | 20 | BP | 0.050 |

^n^, number of genes in the pathway

DE, number of genes that were differentially expressed

Light grey shading represents significant pathways detected at the 4 hr time point

**Supplementary Table 15. Potential disease processes associated with significantly correlated genes at 24 hours**

| Gene set ID | Description | N | Overlap | FDR |
| --- | --- | --- | --- | --- |
| 114480 | Breast cancer (BRCA1, BRIP1, BARD1) | 24 | 3 | 0.01 |
| 104300 | Alzheimer’s disease (APP, PAXIP1) | 11 | 2 | 0.04 |
| 252010 | Mitochondrial complex I deficiency (NDUFV1, TMEM126B) | 20 | 2 | 0.06 |
| 189960 | Tracheoesophageal fistula (BRIP1, RAD51C) | 21 | 2 | 0.06 |
| 601626 | Acute myeloid leukemia (NPM1, TERT) | 22 | 2 | 0.06 |
| 125853 | Noninsulin-dependent diabetes mellitus (PDX1, HMGA1) | 29 | 2 | 0.09 |

^N^ Total number of genes in gene set

^Overlap^ Number of genes present our dataset
