## Supplementary File 2 for "Site-specific decreases in DNA methylation in replicating cells following exposure to oxidative stress"

Validation of the Illumina EPIC array hits using bisulfite-based amplicon sequencing

The Illumina Infinium MethlationEpic beadChip array, is considered as the current gold standard in detection of DNA methylation at the genome-wide level. As such, alternative techniques do not have equivalent reliability, reproducibility or potential for high throughput screening. Therefore, many studies do not consider it applicable to validate the results of the methylation array using an inferior technique. Here, we applied an alternative DNA methylation detection technique referred to as bisulfite based amplicon sequencing (BSAS) for validation of a number of the top most significant differentially methylated CpGs observed in our study. BSAS was able to detect differential DNA methylation all investigated CpG sites, to a degree that correlated closely with the EPIC array, at all except one CpG.

| Probe ID | Gene | Genome region | PCR length | Forward | Tm | Reverse | Tm |
| --- | --- | --- | --- | --- | --- | --- | --- |
| cg05208399 | MAPK8IP3 | chr16:1816268-1815668 | 229 | TTTGTTGGGAAAGTGGAAGGTTAG | 57 | AACACAACCCACCAAAATAATACC | 55 |
| cg15047582 | TFAP2E | chr16:1814096-1813496 | 208 | GGTTTTGATTAATGTGGGTTGAATTT | 54 | CRCCCTTCCTCCTCTAAAC | 56 |
| cg10553894 | CPT1A | chr11:68550834-68550234 | 185 | TAGGTTTTGAGAGGAGGGTAT | 54 | ATAAAAACCTAAAAAAAAACCAAACC | 50 |
| cg13782884 | FAM46B | chr1:27339587-27338987 | 236 | GGTTGGGTAGGGGTTAAGG | 57 | AAACTCTACAAACCAATATCCGAAAC | 57 |
| cg05372828 | KCNAB2 | chr1:6111932-6111332 | 250 | TTAGGGTTAGTTATTTAGTTTGGGGTA | 56°C | TCTCCATATTCCTAACCTAAACAACC | 55°C |
| cg16852704 | TLX2 | chr2:74743543-74742943 | 148 | GGTTGTTTTTGTATTTGTAGTAGGA | 52°C | AACACTAAAACCACTTTATTATCCTC | 52°C |
|  |  | Illumina Sequencing barcode |  | TCGTCGGCAGCGTCAGATGTGTATAAGAGACAG |  | GTCTCGTGGGCTCGGAGATGTGTATAAGAGACAG |  |

**Table 1. DNA sequence of oligonucleotide primers used during PCR amplification of bisulfite converted DNA.**

Results

The differences between differential methylation in the treatment and control groups for each time point were investigated using BSAS and compared against the results obtained using the EPIC array (Tables 2 and 3). When comparing treatment and control samples at the four hour time point, there were no significant differences in DNA methylation after correction for multiple testing. However, prior to adjusting for multiple testing, all gene regions except for KCNAB2 demonstrated a trend towards consistent results. The logFC between the two methods were of similar magnitude, however, the larger variation observed with BSAS significantly reduced the statistical power. When assessed at the 72 hour time point there were no significant changes observed using either the BSAS or the EPIC methods, both of which had a similar logFC.

A Bland Altman analysis was used to compare the differences in Beta values from BSAS and the Illumina Epic array, against the mean value for the treatment and control groups (Figure 1). The observed differences for each group fell within the lines of agreement, however the treatment group showed a substantial negative bias, which was likely skewed by the lower sensitivity of BSAS in detecting unmethylated DNA (Figure 2).

**Table 2: CpG site differences from BSAS and the EPIC array methods at the 9 loci (differing levels of significance) between the treatment and control groups at the four hour time point**

| Gene | CHR | Position | Probe ID | logFC | *P* Value | adj*.P* Value | Epic Adj. *P* Value | Epic logFC |
| --- | --- | --- | --- | --- | --- | --- | --- | --- |
| MAPK8IP3 | 16 | 1815968 | cg05208399 | -1.76 | 0.03 | 0.14 | 0.00 | -1.8 |
| KCNAB2 | 1 | 6111671 | cg02630629 | -0.80 | 0.07 | 0.14 | 1.00 | -0.5 |
| TLX2 | 2 | 74743243 | cg16852704 | -1.14 | 0.07 | 0.14 | < 0.001 | -1.75 |
| FAM46B | 1 | 27339287 | cg13782884 | -0.55 | 0.08 | 0.14 | < 0.001 | -1.8 |
| TFAP2E | 1 | 36038869 | cg15047582 | -1.19 | 0.08 | 0.14 | 0.00 | -2 |
| KCNAB2 | 1 | 6111632 | cg05372828 | -0.99 | 0.11 | 0.16 | < 0.001 | -1.8 |
| FAM46B | 1 | 27339413 | cg24481381 | -0.30 | 0.16 | 0.21 | 0.09 | -1.2 |
| FAM46B | 1 | 27339314 | cg06606318 | -0.08 | 0.77 | 0.87 | 1.00 | -0.1 |
| FAM46B | 1 | 27339435 | cg22395192 | 0.02 | 0.89 | 0.89 | 0.08 | -0.75 |

**Table 3: CpG site differences from BSAS and the EPIC array methods at the 9 loci (differing levels of significance) between the treatment and control groups at the 72 hour time point**

| Gene | CHR | Position | Probe ID | logFC | P.Value | adj.P.Val | Epic Adj. *P* Value | Epic logFC |
| --- | --- | --- | --- | --- | --- | --- | --- | --- |
| TLX2 | 2 | 74743243 | cg16852704 | 1.07 | 0.09 | 0.56 | 0.39 | 0.33 |
| MAPK8IP3 | 16 | 1815968 | cg05208399 | -1.07 | 0.16 | 0.56 | 0.90 | 0.14 |
| FAM46B | 1 | 27339413 | cg24481381 | 0.28 | 0.20 | 0.56 | 0.10 | 0.90 |
| FAM46B | 1 | 27339435 | cg22395192 | -0.18 | 0.25 | 0.56 | 0.63 | 0.25 |
| FAM46B | 1 | 27339314 | cg06606318 | 0.19 | 0.49 | 0.80 | 0.94 | 0.03 |
| TFAP2E | 1 | 36038869 | cg15047582 | -0.39 | 0.53 | 0.80 | 0.54 | 0.44 |
| KCNAB2 | 1 | 6111671 | cg02630629 | 0.04 | 0.92 | 0.99 | 0.98 | -0.01 |
| KCNAB2 | 1 | 6111632 | cg05372828 | -0.02 | 0.97 | 0.99 | 0.89 | -0.07 |
| FAM46B | 1 | 27339287 | cg13782884 | 0.00 | 0.99 | 0.99 | 0.57 | 0.23 |

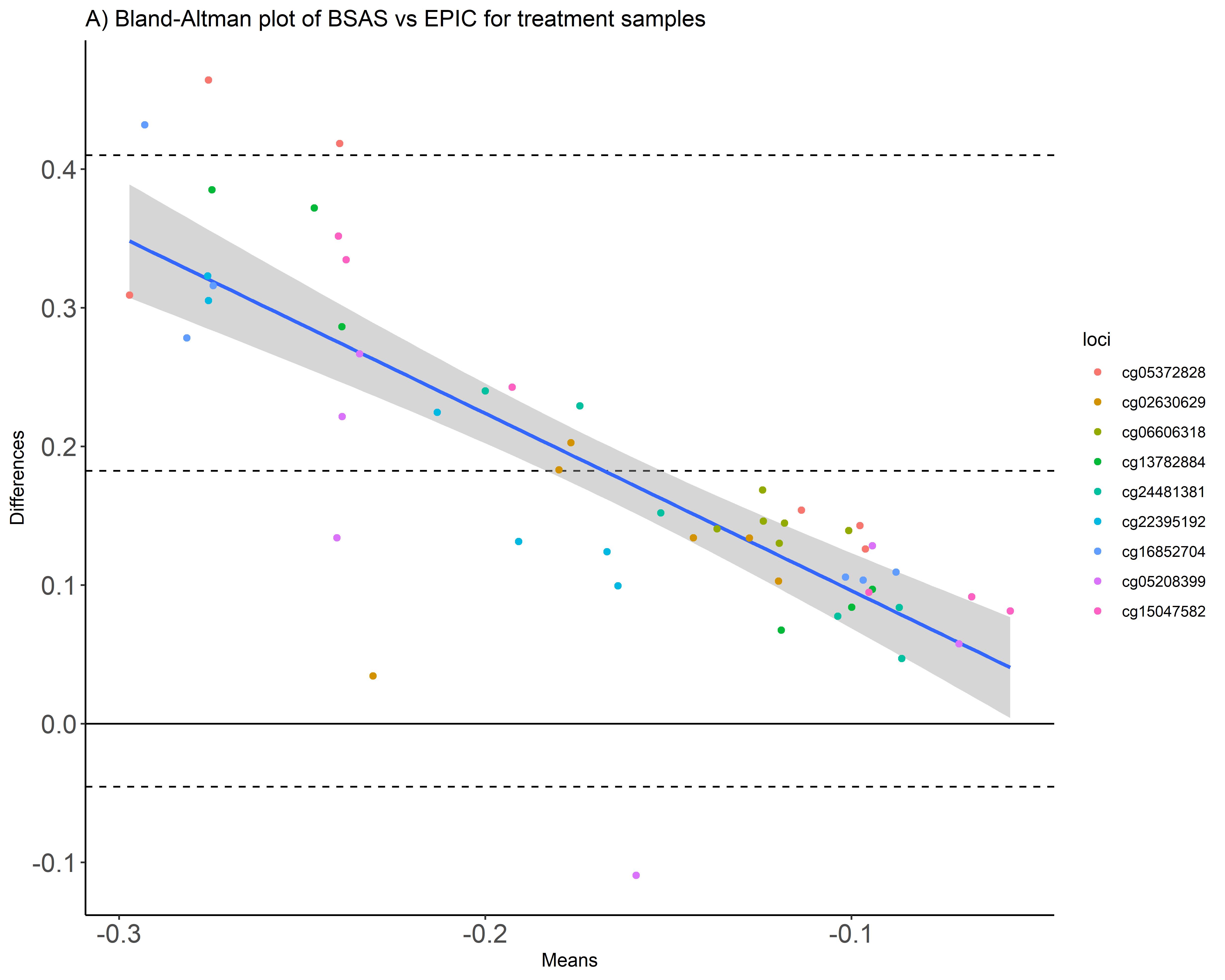

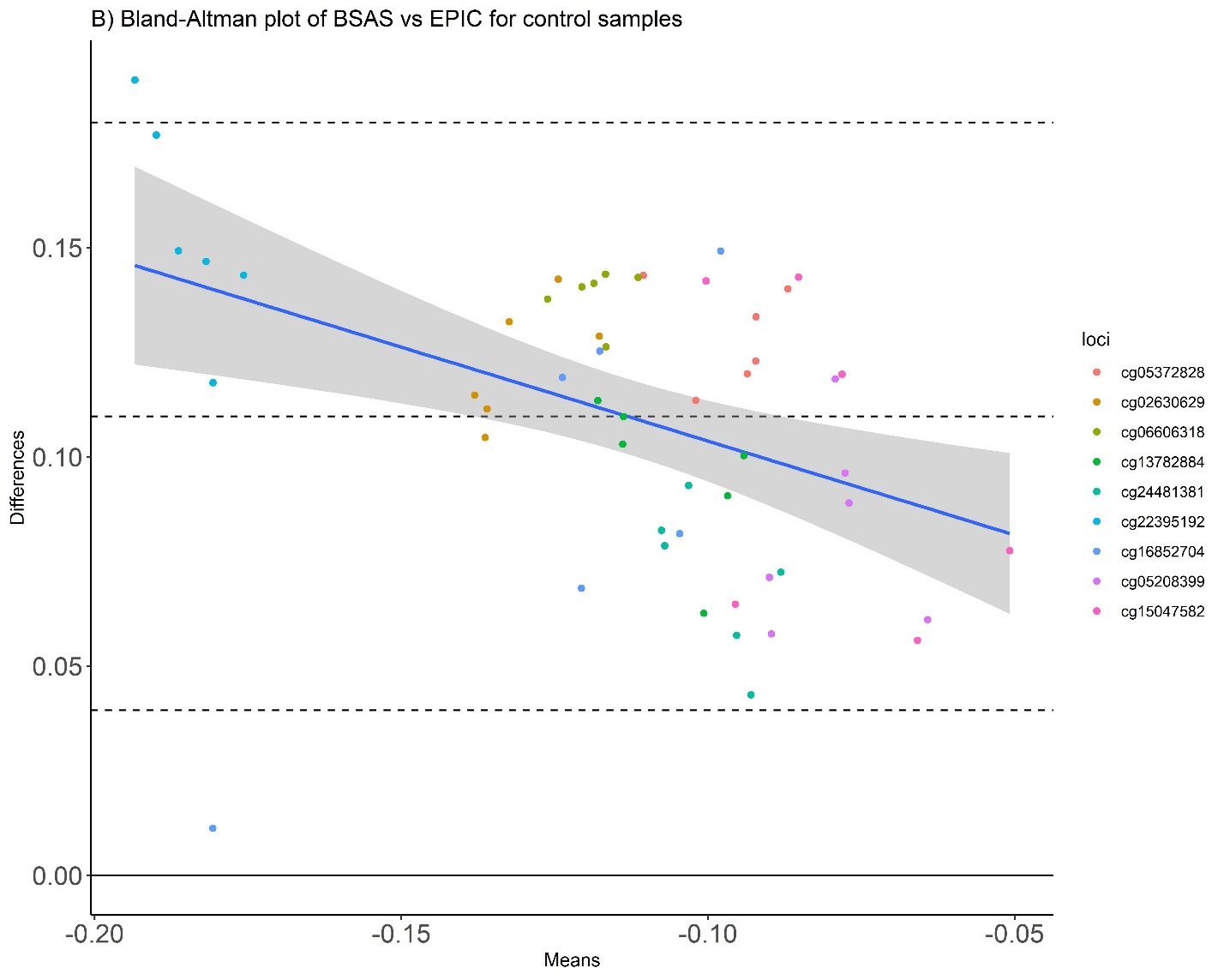
Figure 1. Bland Altman plots for treatment and control samples, using the mean, and differences between methylation values quantified by BSAS and by the EPIC array. A) Samples analysed from the treatment groups B) Samples analysed from the control groups. Each of the 9 CpGs were plotted, and the blue line demonstrates the proportional bias regression line, with standard error shaded in grey. The difference between Beta values is represented on the Y axis and the average (log transformed) Beta value is represented on the X axis. The top and bottom dashed lines represent +/- 2 standard deviation of the mean.

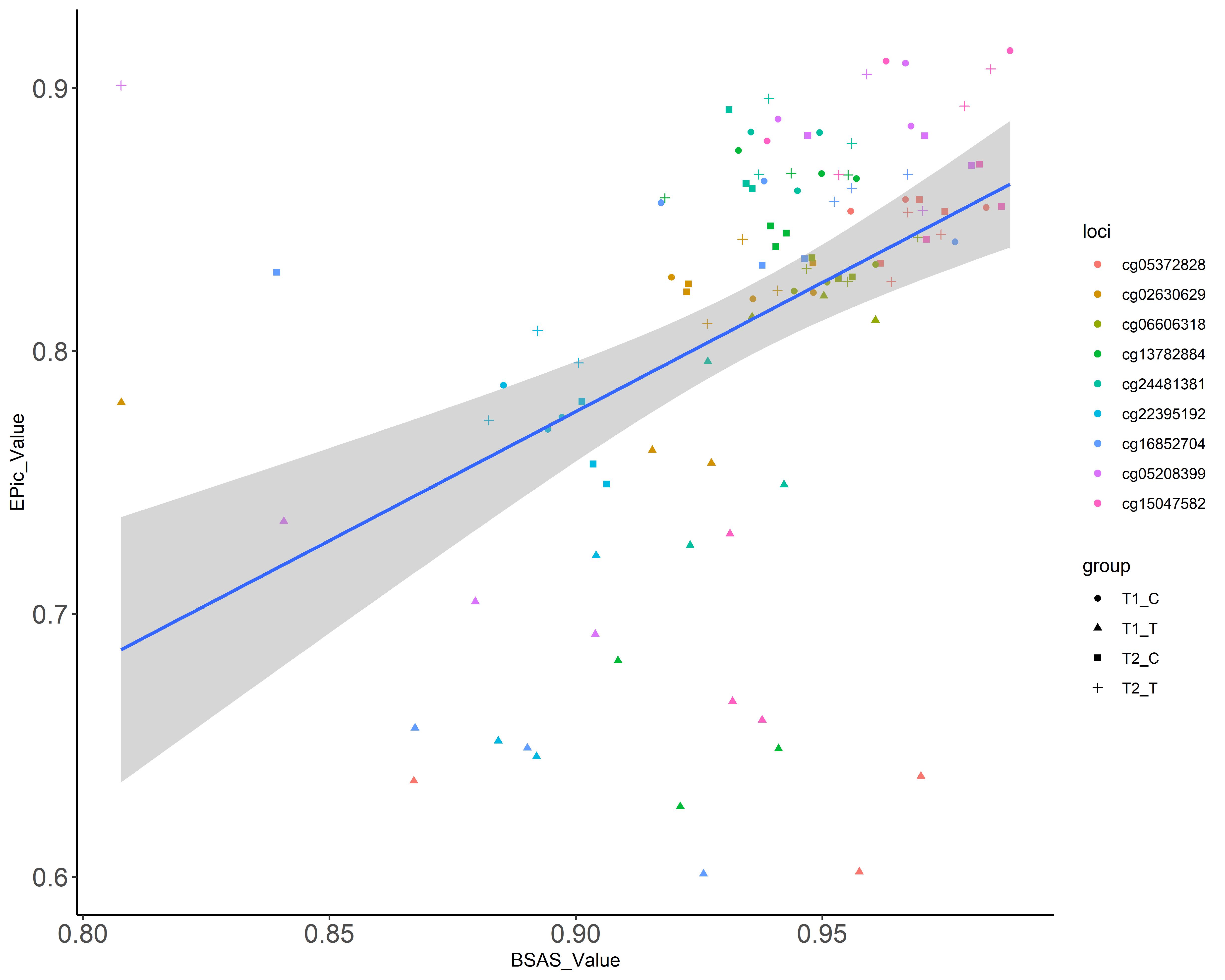

**Figure 2. Correlation between Beta values detected using BSAS and the EPIC array**. Values from the EPIC array are represented on the Y-axis and values from BSAS are represented on the X-axis. The regression line is demonstrated in blue, and the confidence intervals are represented by grey shading. T1_C are the control samples from the 4 hr time point, T1_T are the Treatment samples from the 4 hr time point, T1_C are the control samples from the 72 hr time point, and T2_T are the treatment samples from the 72 hr time point.
